# Population genetics, trait mapping and fungal pathogen surveillance using untargeted sequencing in timber rattlesnakes (*Crotalus horridus)*

**DOI:** 10.1101/2025.10.21.683524

**Authors:** Christopher Husted, Jacob Alonso, Ross Swofford, Anne G. Stengle, Lucas R. Moreira, Daniel E. Neafsey, Rachel A. Johnston, Eric Baitchman, Diane P. Genereux, Rachel Daniels, Elinor K. Karlsson

## Abstract

Timber rattlesnakes (*Crotalus horridus*) face escalating threats in the Northeastern Appalachians, including habitat fragmentation, human encroachment, and the fungal pathogen *Ophidiomyces ophiodiicola*. Using untargeted sequencing of DNA extracted from scale clips, we generated both host whole-genome and metagenomic data for 97 snakes from eight populations. Analysis of the snake genomes shows the populations surveyed exhibit relatively low levels of inbreeding and are genetically distinct, but that the degree of separation correlates only weakly with geographic distance. A genome-wide association analysis identified a locus associated with black-to-yellow color variation that contains an aldehyde dehydrogenase gene (*ALDH4A1*) related to genes involved in hair color differences in humans. Metagenomic analysis showed that *O. ophiodiicola* read counts were generally higher in snakes exhibiting clinical signs of Snake Fungal Disease, but some visually asymptomatic snakes had high pathogen loads.

Together, these findings highlight the dual utility of untargeted sequencing for population genetics and pathogen surveillance, providing a foundation for future studies of adaptation, disease dynamics, and conservation in this declining species.

## Introduction

The timber rattlesnake (*Crotalus horridus*) is a long-lived pit viper usually found in the deciduous forests and rugged terrain of the northeastern Appalachians. Once the northernmost rattlesnake, reaching as far as southern Ontario, the species has declined sharply because of habitat loss, road mortality, human persecution, and disease (Clark et al., 2011; Hohenlohe et al., 2021; Montague, 2022; A. Stengle, 2018) Today, it is regionally extinct in Canada, and in many U.S. states, including Michigan, Delaware, Maine, and Rhode Island (COSEWIC, 2023; Keyler, 2023; W. H. Martin et al., 2008), and listed as endangered in New Jersey, Ohio, Vermont, Connecticut, Massachusetts, Virginia, Indiana and New Hampshire, where only one population survives (Bushar et al., 2015; Clark et al., 2011; Furman, 2007). Timber rattlesnakes mature late and produce relatively few young, and the loss of even a small number of adults can significantly affect population survival(Heppell et al., 2000).

One of the most serious and growing threats to remaining timber rattlesnakes populations is Snake Fungal Disease, caused by the pathogen *Ophidiomyces ophiodiicola* (Allender, Raudabaugh, et al., 2015; Lorch et al., 2015, 2016). Affected snakes often develop skin lesions, may change their behavior, and have increased mortality (Allender et al., 2018; Lorch et al., 2016; McKenzie et al., 2021). However, the absence of lesions at a given time point does not necessarily indicate an absence of infection or disease(McKenzie et al., 2019). Lesions can temporarily resolve following a shed cycle, and recrudesce when the animal is stressed or cooled(Lind et al., 2023). The transmission dynamics are not yet well characterized, although infections can be vertically transmitted from mothers to offspring and the fungus can persist in the environment without a host (Haynes et al., 2020; A. G. Stengle et al., 2019).

Understanding the genetic diversity of surviving timber rattlesnake populations is critical for conservation. Previous studies on timber rattlesnake population genetics using microsatellites have shown a recent reduction in genetic diversity due to human encroachment and habitat fragmentation in New Jersey (Bushar et al., 2015), localized genetic structure among hibernacula (Clark et al., 2010), significant population bottlenecks in New Hampshire(Clark et al., 2011) and high levels of differentiation between populations (A. Stengle, 2018). However, these studies genotyped a relatively sparse set of markers across the genome, and lacked the resolution needed to detect fine-scale connectivity or link genetic variation to ecologically important traits.

A potentially powerful alternative approach is low-coverage whole-genome sequencing (skim-WGS). skim-WGS is an efficient, low-cost, and versatile method for generating dense genomic data from samples with limited DNA input (Fuentes-Pardo & Ruzzante, 2017; Homburger et al., 2019; Theissinger et al., 2023). It has previously been used to discover the genetic basis of traits relevant to health (Chat et al., 2021; Li et al., 2023). A key advantage of skim-WGS is its untargeted sequencing strategy, which captures all DNA in a sample, including host genomic DNA as well as microbial and fungal DNA (Adhikari et al., 2022; Lou et al., 2021). This comprehensive output (“whole-sample reads”) is especially valuable in timber rattlesnakes because of the threat posed by Snake Fungal Disease. Unlike amplicon-based methods, skim-WGS directly captures microbial DNA without primer bias, providing a more quantitative view of microbial community composition and enabling investigation of interactions among the host, the microbiome, and pathogens.

Here, we sequenced and analyzed DNA obtained from scale-clips from 111 timber rattlesnakes sampled across six populations in the Northeastern Appalachians to: (1) characterize population structure, inbreeding, and connectivity; (2) identify genomic variants associated with the yellow-to-black color morph which may play a role in thermoregulation (Allsteadt, 2003; Rhodes et al., 2023); and (3) explore geographic variation in skin-associated microbial communities, with a focus on *O. ophiodiicola*.

## Materials and Methods

### Sample collection and preparation

Timber rattlesnake scales were initially collected for a previous study between 2009 and 2014 and obtained as 2 mm scale clips preserved in 95% ethanol or air-dried (A. Stengle, 2018). DNA was extracted from scales using the DNA Extractor® FM Kit (Product Number: 295-58501, FUJIFILM Wako Pure Chemical Co., Richmond, VA). Each scale was removed from the sample, dried with a Kimwipe, and transferred to a 1.5 mL microcentrifuge tube. The manufacturer’s protocol and amounts for the Lysis, Enzyme-activated Reagent (EAR), and Protease Solution were used, with the following modifications: the tube was gently tapped to mix rather than vortexed, and the tube was incubated at 55°C with gentle rocking. After 2 hours of incubation, an additional 10 µL of Proteinase-K was added, and the incubation continued for 24 hours. If the scale was not completely digested, another 10 µL of Proteinase-K was added before proceeding and incubated for no longer than 24 hours. Following digestion, the sample was centrifuged at 16,000 x g for 5 minutes at room temperature, and the supernatant was transferred to a new 1.5 mL tube. DNA was precipitated by adding 250 µL of NaI and 450 µL of isopropanol in that order, followed by vortexing and incubation at room temperature for 15 minutes. The sample was then centrifuged and washed following the manufacturer’s protocol. A final incubation at 37°C for 5 minutes was performed to evaporate any residual ethanol before proceeding with downstream processing. A commercial sequencing company (Gencove Inc., New York, NY 10016) processed the final extracted samples for DNA-Seq library preparation and sequencing.

### Genomic data generation and processing

Whole-genome sequencing was performed on the MiSeq Illumina platform at 151 base pair read length, targeting 1x coverage. The raw fastq files were then trimmed using Cutadapt to remove low-quality base pairs and adaptors, discarding reads <20 base pairs post-trimming (M. Martin, 2011). The trimmed reads were aligned to both the *C. horridus* reference genome (ASM162548v1; 5,707 scaffolds; contig N50 = 2.3 Mb) (Rhoads et al., 2024) and the more contiguous eastern diamondback rattlesnake (*C. adamanteus*) reference genome (Cadamanteus_3dDNAHiC_1.2; 26 scaffolds; contig N50 = 67.5 Mb) using the Burrows-Wheeler Aligner (BWA). The lower contiguity of the *C. horridus* assembly negatively impacted mapping, so we used the *C. adamanteus* alignment for subsequent host variant calling. We note that a more contiguous reference for C. horridus was released after we completed these analyses (Roseman et al., 2025). Duplicate reads were removed using Picard Tools to minimize biases, and read groups were added while filtering out supplementary alignments. A similar alignment method was also performed in parallel to align the raw reads to the *O. ophiodiicola* genome (Ladner et al., 2022). After aligning the reads to the *O. ophiodiicola* genome, counts were normalized by dividing aligned reads by total sequenced reads. A Wilcoxon rank sum test was then performed to identify differences in normalized *O. ophiodiicola* read counts between individuals of the various sample populations.

### Population structure analysis

Genotyping and downstream analyses were performed using ANGSD (version: 0.935) (Korneliussen et al., 2014). Samples with coverage <0.051x were excluded (14/111), yielding 97 individuals for analysis. ANGSD parameters were set to obtain genotype likelihoods, computed using the SAMtools method (-GL 2), with filtered BAM files as input. Output options were enabled to obtain genotype probabilities, genotype likelihood files, and count data. Minor allele frequencies (-doMaf 1) and major and minor alleles were also estimated. Posterior probabilities of genotypes were calculated, keeping SNPs with a minimum of 90% (N=87) (-minInd 87) individuals represented and a minimum minor allele frequency of 0.05. Additional quality filters included a minimum mapping quality of 30, a minimum base quality of 20, and a stringent SNP p-value threshold of 1e-6. Principal component analysis (PCA) was conducted using PCAngsd, with default parameters and settings (Meisner & Albrechtsen, 2018). To validate the PCA results, particularly against potential sequencing coverage variability, the CollectWgsMetrics tool from the GATK platform was used (McKenna et al., 2010).

To visualize population structure, genotype likelihoods were initially estimated using ANGSD and analyzed with PCAngsd to create a covariance matrix (Meisner & Albrechtsen, 2018).

Principal components were then calculated in R using eigen decomposition of this matrix. The resulting eigenvectors represented sample coordinates, and the eigenvalues were used to calculate the percentage of variance explained by each component. Genetic structure was assessed and visualized using two complementary approaches. First, scores for the top principal components were plotted in ggplot2 (Wickham, 2016; Wilkinson, 2011). To partition variance by population identity, linear models were fit with town (factor) and mean coverage as predictors of each PC score. To quantify contributions of location and sequencing depth to a given PC, linear models were fit with town and mean coverage as predictors; sequential ANOVA estimated unique variance explained. Second, Uniform Manifold Approximation and Projection (UMAP; R package *uwot*) was used to identify finer-scale and potentially nonlinear structures among phenotyped individuals, employing the top 10 PCs from PCAngsd as input (*n_neighbors* = 15, *min_dist* = 0.5)(Melville, 2025).

Runs of homozygosity (ROH) were identified using bcftools roh, utilizing BCF files generated by ANGSD as input. Because bcftools roh is sensitive to low/uneven coverage, we applied a final coverage filter, retaining only samples above an outlier cutoff of median − 3x MAD of log10-coverage, back-transformed; in this dataset, the threshold was 0.241x. Identified ROH segments were converted into BED format, and pairwise comparisons between populations were performed using BEDTools. Specifically, the Jaccard index was calculated to measure ROH overlap between each population pair, alongside coverage metrics indicating the proportion of each population’s ROH segments overlapping with another population’s segments. A weighted Jaccard calculation was also performed using 100-kb and 1-Mb windows to evaluate ROH frequency across individuals. Sex chromosomes were excluded from the analysis because their inheritance patterns, such as hemizygosity and reduced recombination, can lead to biased ROH detection and artificially inflated homozygosity (Sun et al., 2023).

### GWAS for color phenotype

ANGSD-assoc (Jørsboe & Albrechtsen, 2022) was used to perform the GWAS for coloration phenotype, coded yellow vs. black (A. Stengle, 2018). BEAGLE-formatted genotype likelihoods were generated against the *C. adamanteus* reference. Parameters mirrored the population-structure analysis, except that loci were required in ∼75% of phenotyped individuals. Minor allele frequency estimation (-doMaf 4) and latent genotype modeling (-doAsso 4) were performed to explore the genetic associations with various phenotypic traits. This approach incorporated binary phenotypic data and adjusted for population structure using the first three principal components from the 49 sample results passed through pcangsd.

An additive model (-model 1) was used for animal color because additive genetic variance primarily contributes to the genetic architecture of complex traits, including color morphs (Hill et al., 2008). The ANGSD output was run through BEAGLE for imputation. To control for multiple testing, a genome-wide significance threshold was defined using the Bonferroni correction, based on a family-wise error rate of α = 0.05 and the total number of populations tested (n = 1,159,481). The resulting significance threshold was 4.31 x 10^-8^. The ANGSD-assoc output was processed and visualized in R using the ggplot2 package (Wickham, 2016; Wilkinson, 2011). To annotate significant loci with recognizable gene symbols, we lifted over the GTF for the *Crotalus tigris* genome assembly ASM1654583v1(Margres et al., 2021) to the *C. adamanteus* genomic coordinates using liftoff (Shumate & Salzberg, 2021). The *C. adamanteus* annotation file is available in the Dryad data repository for this paper.

### Microbiome identification and assessment

To comprehensively analyze the microbiome and identify differences in microbiome composition between sample populations, host reads were aligned to the *C. adamanteus* genome, and any reads that did not align were retained. Then, a custom reference database was created, which included the standard bacterial and fungal genomes from Kraken2 (Wood et al., 2019) and reference genomes of human and *O. ophiodiicola*, the fungus that causes Snake Fungal Disease. Kraken2 was run on the raw FASTQ files for each snake sample. To obtain more accurate genus-level abundance estimates, Kraken2 reports were processed using Bracken (Lu et al., 2017). Shannon diversity indices were calculated from Bracken-derived abundances using the vegan R package and compared across six geographic locations using Wilcoxon rank-sum tests (Oksanen et al., 2025).

To compare microbial communities among locations, between-sample composition and within-group variability were quantified. Additionally, Bray–Curtis distances were used to quantify between-sample differences in community composition. Bray–Curtis distances were computed from operational taxonomic unit (OTU) matrices. To test whether groups differed in within-group spread (beta-dispersion), we used PERMDISP (betadisper) with 999 permutations, reporting overall and pairwise tests. Comparisons were adjusted for multiple testing using the false discovery rate (FDR) method. Centroid separation among towns was tested using PERMANOVA (adonis2 function), with pairwise comparisons conducted via pairwise.adonis2, adjusting p-values with FDR correction.

## Results

### Study populations and sample collection

Our study included 111 timber rattlesnakes sampled from eight geographically distinct locations across the Northeastern Appalachians (Figure 1 A, Table S1). Four populations were located in the Berkshire–Taconic region: Mt. Washington, Massachusetts (MA; N = 40); Copake, New York (NY; N = 11); and two sites near Rutland, Vermont (VT), referred to as location 1 (L1; N = 26) and location 2 (L2; N = 10). The remaining populations were sampled from Westfield, MA (N = 13); Milton, MA (N = 5); Rockland County, NY (N = 2); and an undefined location in central Pennsylvania (N = 10). All sites occur within predominantly northern hardwood–mixed forests typical of the Northeastern Appalachians, a critical habitat for timber rattlesnakes (Eckert & Jesper, 2024). We had information on color phenotype (black vs. yellow morphs) for 50 individuals (with one individual filtered due to being too low coverage), and on the presence or absence of snake fungal disease for 36 individuals, all from Copake, NY (N_color_=11; N_disease_=9) or Mt. Washington, MA (N_color_=38; N_disease_=27) (Figure 1 B,C; Table S2).

**Figure 1.**
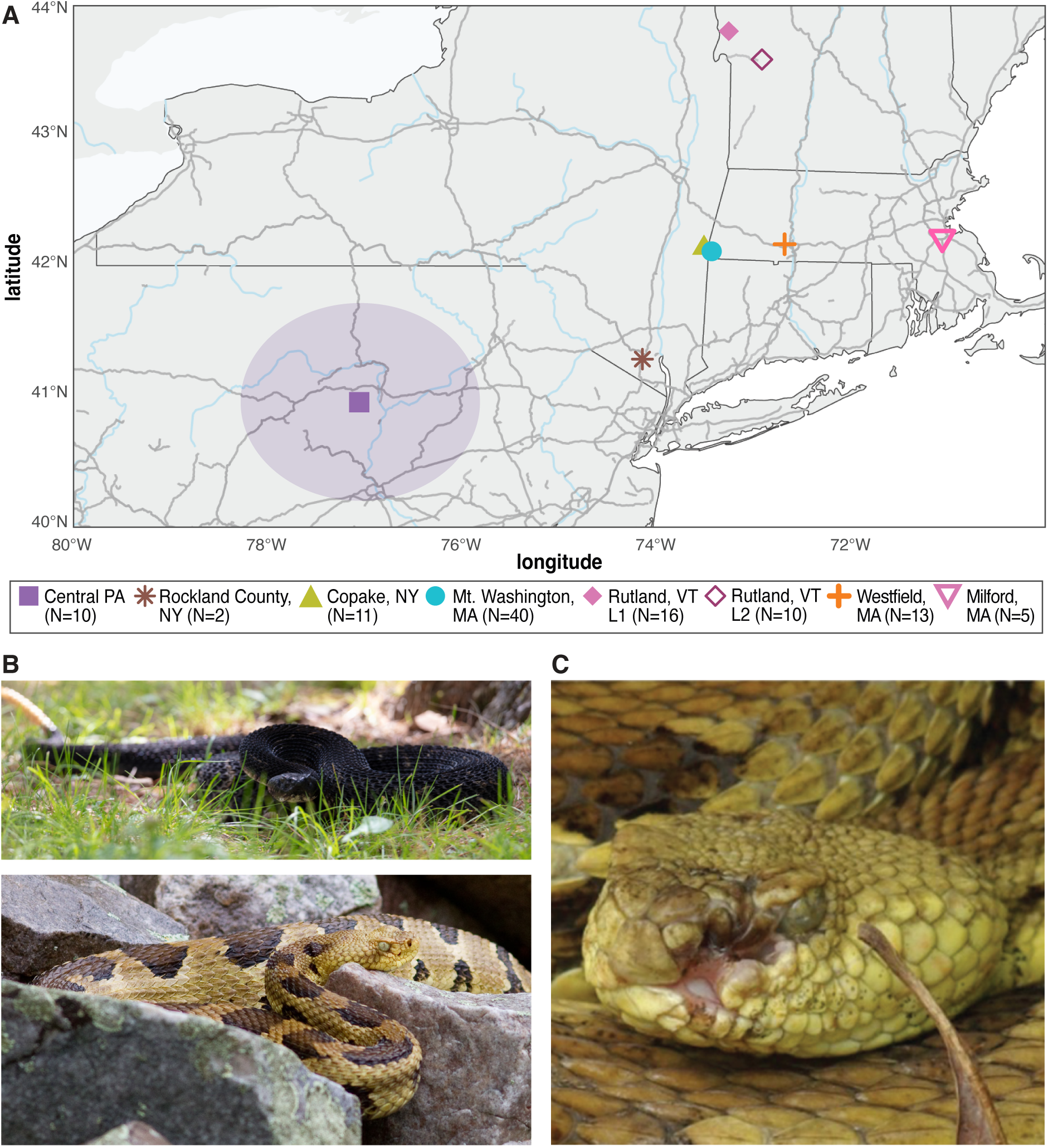
Snakes from eight geographically separated populations, some with phenotype information, were included in the study. (A) Map of Northeastern Appalachian sampling sites in Pennsylvania (PA), New York (NY), Massachusetts (MA) and Vermont (VT), with sample sizes indicated in the legend. Purple oval indicates that the exact locations of the sites in central PA are not well described. (B) Timber rattlesnake Crotalus horridus color morphs include black (top) and yellow (bottom). (C) Example of snake fungal disease (O. ophiodiicola) presenting as a lesion on the head. Photo credit (B) M. O’Neill, NPS, (C) Brian Gratwicke and (D) A. Stengle.

### Making a host-aligned sequencing dataset

DNA was extracted from scale clips obtained from live snakes for a previous study (A. Stengle, 2018) and sequenced on an Illumina platform, generating an average of 4.7 ±LJ3.0 Gb of raw sequencing data per individual(Table S1). Sequencing reads were aligned to both the *C. horridus* and the more contiguous *C. adamanteus* reference genomes, with an average of 76 ± 25% and 71 ± 23% of reads mapping, respectively, which is consistent with their relatively recent divergence(∼10.9 million years ago)(Kumar et al., 2022)(Table S3 and Table S4). We used the *C. adamanteus* alignment for host variant calling because its greater contiguity improved genotype likelihood estimates(Korneliussen et al., 2014).

The final host dataset included 97 samples with mean genome-wide coverage exceeding 0.05x (Table S5), representing six of the eight populations. Samples from Milton, MA, were excluded because none had sufficient host genome coverage, and Rockland, NY, was excluded from most analyses (host and microbiome) due to having only two individuals. The retained samples had mean alignment rates of 79% ± 8% and mean coverage of 1.03x ± 0.56x (range: 0.06– 3.62x) (Tables S3, S5). Coverage differed among populations, potentially reflecting differences in sample quality (24% of variance explained in a one-way ANOVA; p = 0.0004). We genotyped 381,321 high-confidence single-nucleotide polymorphisms (SNPs) per individual, distributed across the autosomal genome at an average spacing of ∼3,889 bp (0.257 SNPs/kb)(Korneliussen et al., 2014). This represents a marker density roughly an order of magnitude(∼8-13x) higher than typical dinucleotide microsatellite spacing in vertebrate genomes(Chen et al., 2005; Gibbs et al., 1998; Holycross et al., 2002; Villarreal et al., 1996).

### Fine-scale genetic structure and isolation-by-distance

#### Population differentiation

Historical gene flow among snake populations was only weakly correlated with geographic distance. The six populations included in the host-genome analyses had a mean pairwise separation of 211 ± 144 km (range: 5–450 km; Table S6). Pairwise weighted F_ST_, a measure of genetic differentiation between populations, was not significantly associated with geographic distance (Figure 2A; Table S6). Notably, the two populations near Rutland, VT, exhibited the highest differentiation (F_ST_ = 0.20) despite being less than 40 km apart, whereas the lowest differentiation occurred between Rutland (L1) and Westfield, MA, which are separated by ∼160 km.

**Figure 2.**
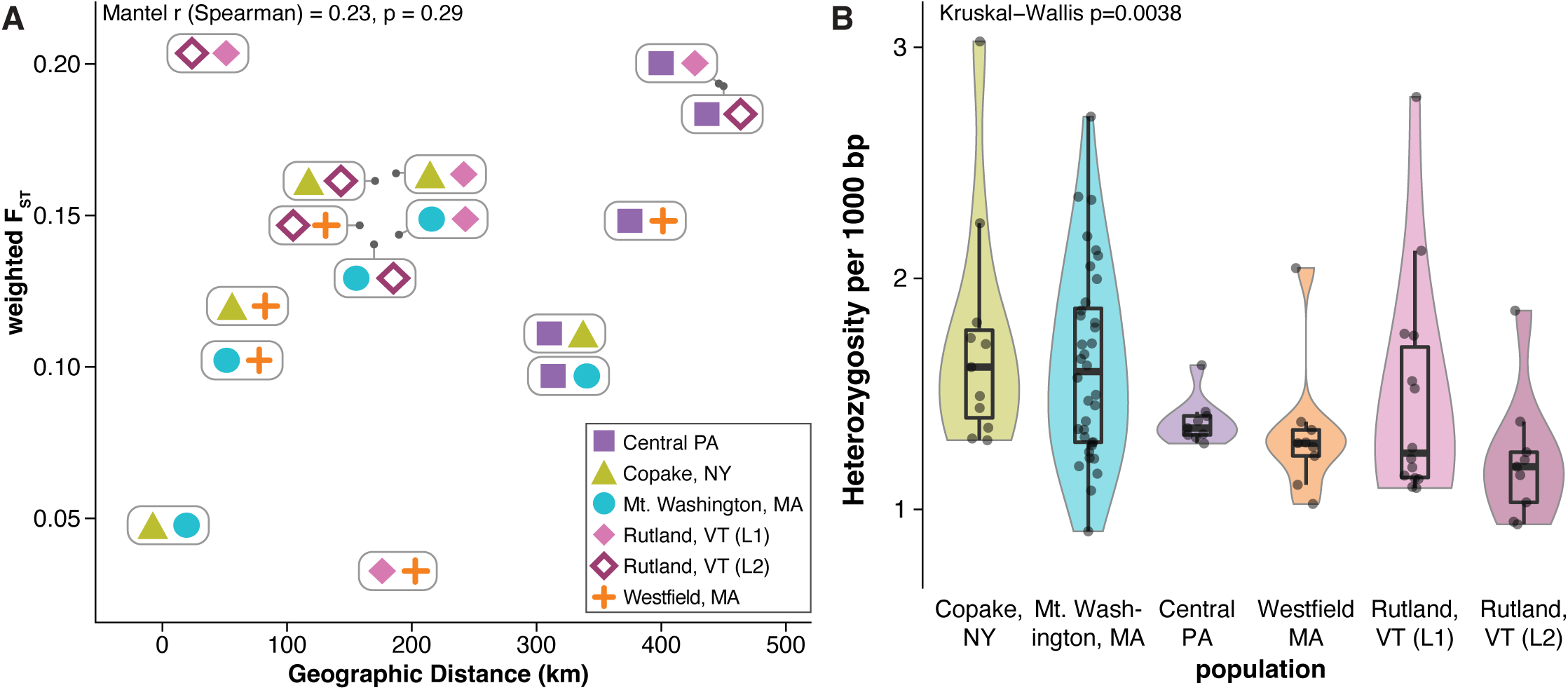
The five snake populations included in the snake genomic analysis are genetically differentiated. (A) Pairwise weighted F_ST_ between populations was not associated with geographic distance. (B) The number of heterozygous positions in individual snakes varies between populations. Violins are scaled to equal widths.

We also observed substantial within-population diversity. The average number of heterozygous calls ranged from 0.37 to 3.0 per 1,000 bp (Table S7), with Mt. Washington, MA, and Copake, NY, showing higher heterozygosity than other populations (Figure 2B). Similar patterns were found for nucleotide diversity (π), measured as the mean rate of pairwise heterozygosity per bp, which ranged from 0.0015 in Rutland, VT (L2) to 0.0023 in Mt. Washington, MA (Table S8).

#### Extent of homozygosity

Genome-wide runs of homozygosity (ROH) suggest that the five timber rattlesnake populations we surveyed have inbreeding levels similar to Western massasauga rattlesnakes and lower than endangered reptile populations. We measured each snake’s genomic inbreeding coefficient (F_ROH_) as the fraction of the genome in ROH longer than 100kb (Shafer & Kardos, 2025)(Figure 3A). Because this analysis was sensitive to very low coverage data, we applied more stringent coverage thresholds and retained 88 snakes in the analysis. Across all snakes, F_ROH_ is similar to the Western massasauga rattlesnakes (*Sistrurus tergeminus*) and lower than the threatened eastern massasauga rattlesnake (*Sistrurus catenatus*) and the endangered Chinese crocodile lizard (*Shinisaurus crocodilurus*)(Ochoa & Gibbs, 2021; Xie et al., 2022)(Figure 3B, Table S7). The burden of long ROH (>1Mb), which can signal more recent inbreeding (Ceballos et al., 2018; Kyriazis et al., 2025) was likewise comparable to western massasaugas, and possibly lower, and much lower than in eastern massasaugas and the endangered Komodo dragon (*Varanus komodoensis*)(Figure 3C)(Iannucci et al., 2021). These results suggest that, despite population fragmentation, timber rattlesnakes in the studied populations retain considerable genetic diversity and the potential for effective management.

**Figure 3.**
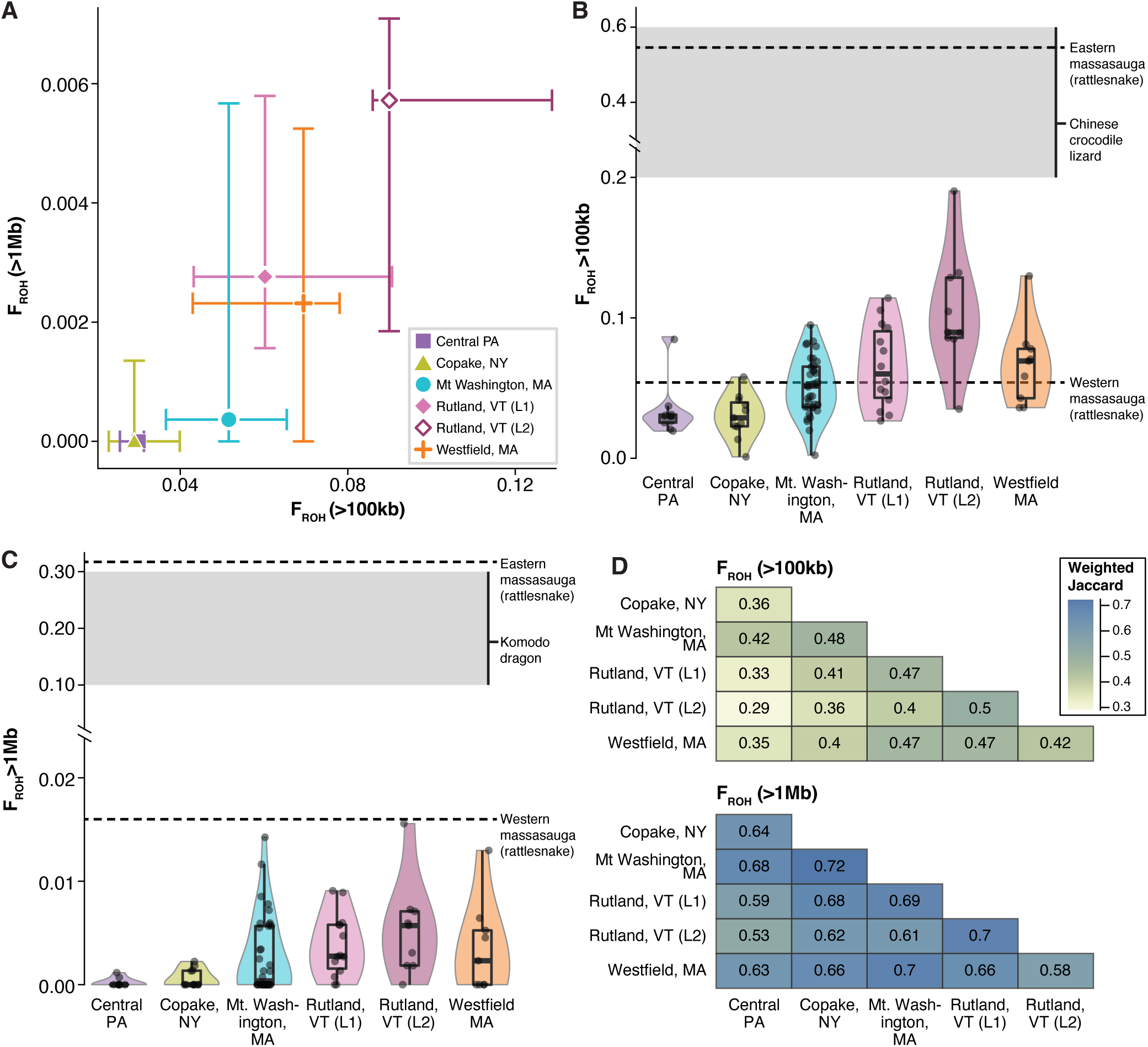
Inbreeding levels vary between populations and are lower than reported in other threatened reptiles. (A) Population medians of F_ROH>100kb_ (x) versus F_ROH>1Mb_ (y) with interquartile-range error bars. (B) Genomic inbreeding coefficients (F_ROH>100kb_) by population compared to other reptiles. (C) Long-tract genomic inbreeding coefficients (F_ROH>1Mb_) by population, with published ranges for other reptiles indicated by green shading and red line. Violins are scaled to equal widths in C and D. The eastern massasauga rattlesnake is listed as a federally threatened species, and the Chinese crocodile lizard and Komodo dragon are listed as endangered by the IUCN Red List. (D) Weighted Jaccard similarity heatmaps for ROH sharing among populations shows tracts > 1Mb (bottom) are more commonly shared across populations compared to all tracts > 100kb (top).

However, interpretation of these results is limited because genomic inbreeding estimates are scarce for reptiles and are largely available only for threatened species.

Long ROH are more commonly shared across populations than short ROH. To quantify this, we compared populations using a frequency-weighted Jaccard index, which measures the overlap of ROH while giving greater weight to those common across individuals. Jaccard values between populations ranged from 0.29-0.51 (mean = 0.41 +/-0.063) for ROH >100 kb, and increased to 0.53–0.72 (mean = 0.65 +/-0.052) for ROH >1 Mb (Figure 3D).

#### Population structure

We found no evidence of sex bias in our study populations. Timber rattlesnakes have a ZZ/ZW sex chromosome system in which females are heterogametic. We inferred the genetic sex of each snake based on the ratio of sequencing reads aligning to the W and Z chromosomes.

Males had W:Z ratios <0.16 and females had ratios > 0.33. Overall, our dataset has comparable numbers of females (N=44) and males (N=53; p_binomial_=0.42), and no difference in the ratio between populations (p_fisher_=0.50).

Individual timber rattlesnakes are genetically similar to other snakes from the same population, but not necessarily more closely related to snakes from geographically proximal populations. In a principal component analysis (PCA) of the whole genome, including the sex chromosomes, PC1 was strongly correlated with the W:Z read-depth ratio, and thus with genetically inferred sex (Figure 4A; Table S10). After removing sex chromosomes, population of origin explained most of the variance along both the first (76%, p = 8.3×10LJ^39^) and second (50%, p = 1.1×10^-18^) principal components, which together accounted for 3.26% and 1.64% of the total genomic variance, respectively (Figure 4B; Table S11). A small proportion of PC1 variation (14%, p = 4.12×10^-12^) and a larger proportion of PC2 variation (23%, p = 9.6×10^-14^) was attributable to sequencing depth(Table S12). Snakes from Vermont and Westfield, MA clustered apart from the other four populations, with those from Rutland, VT (L2) forming a tight cluster consistent with greater relatedness among individuals, whereas Rutland, VT (L1) formed a continuous cluster with Westfield, MA.

**Figure 4.**
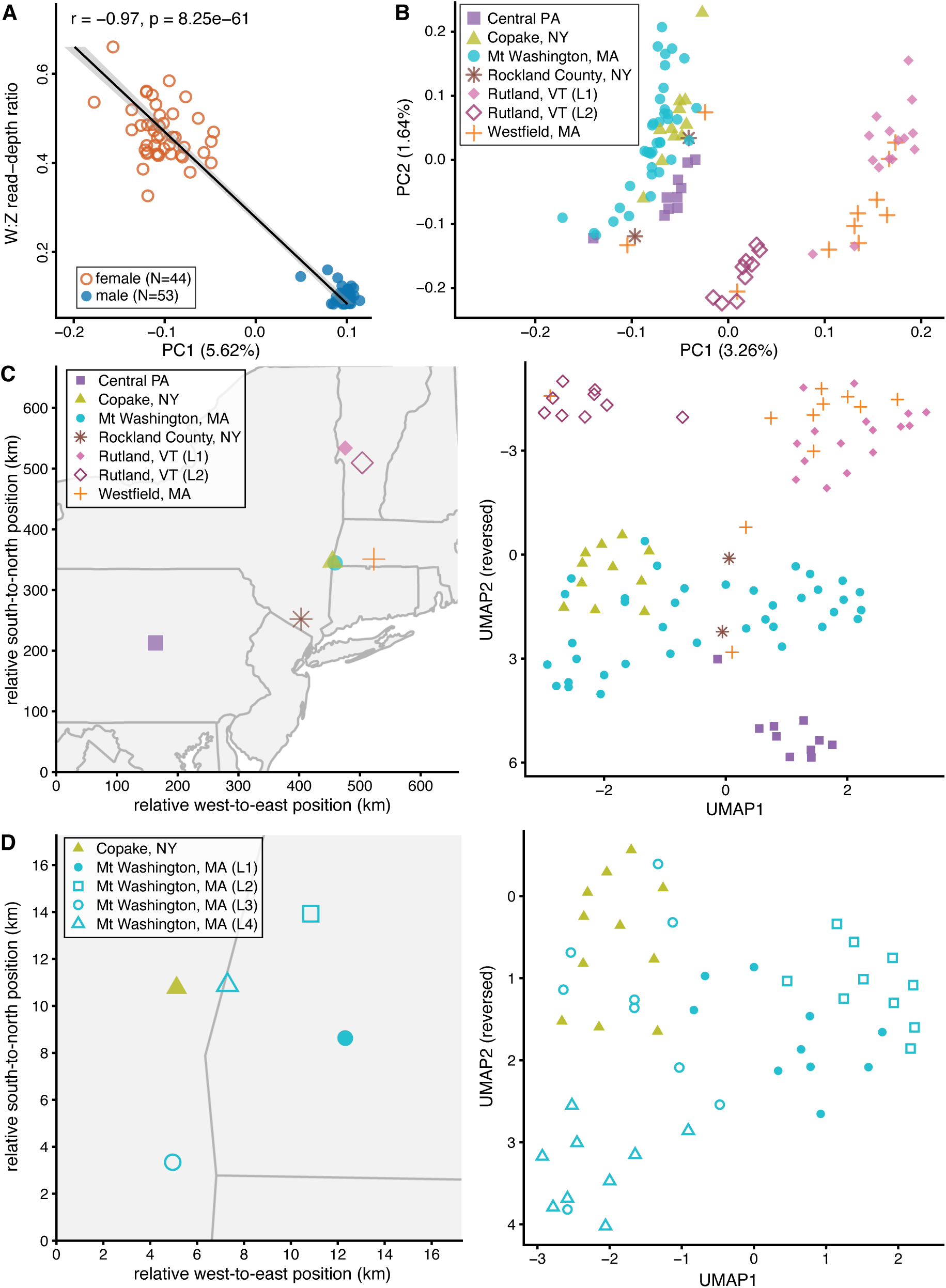
Host genetic analysis reveals sex and population structure in timber rattlesnakes. (A) Principal component analysis (PCA) of all chromosomes separates individuals along PC1 by sex. (B) PCA of autosomes only distinguishes populations from Vermont and one Massachusetts site from other populations along PC1. (C) Geographic relationships among populations (left) are partially mirrored in UMAP clustering (right). (D) Snakes from the same sampling sites within the Mt. Washington, MA and Copake, NY populations (left) are more closely related in UMAP space (right). During the active season, ranges of male and female snakes are ∼2 km^2^ and ∼0.3 km^2^ respectively (Reinert & Zappalorti, 1988).

To capture nonlinear relationships that would be missed by PCA, and visualize fine-scale population structure, we applied Uniform Manifold Approximation and Projection (UMAP), which revealed four main genetic clusters among timber rattlesnakes (Figure 4C; Table S11).

Population explained most of the variance in the UMAP embedding (55.2% for UMAP1 and 82.0% for UMAP2), whereas sequencing coverage explained very little (0.04% and 5.43%, respectively; Table S12). The two Rutland, VT subpopulations (L1 and L2) were genetically distinct despite their close proximity, with L1 snakes clustering together with those from Westfield, MA. As expected from their greater geographic separation, Central PA snakes formed a cluster distinct from the New York and New England populations. The Mt Washington, MA snakes were sampled from distinct four locations, and cluster by their fine-scale geographic sampling locations(Figure 4D; Table S11). A small number of snakes (<5%) clustered with populations outside their sampling location, suggesting rare cases of natural or human-mediated translocation.

### GWAS of colormorph

To investigate the genetic basis of scale color polymorphism, we performed a GWAS on 49 phenotyped timber rattlesnakes (24 yellow and 25 black morphs), treating color as a binary trait in a case–control framework. We tested for association using a likelihood ratio test under a logistic regression model implemented in ANGSD, which accommodates uncertainty in low-coverage data by calculating genotype likelihoods instead of relying on hard genotype calls (Korneliussen et al., 2014). The most associated region spanned six SNPs on chromosome 16 (top SNP p=3.3×10^-10^; Table S13) and contains two annotated genes: *LOC120313277* and *ALDH4A1*, both of which belong to a gene family previously implicated in human pigmentation (Figure 5) (Matsumoto et al., 2019) The major allele at each of the six associated SNPs was highly penetrant for the yellow morph, with 88.9–100% of individuals homozygous for the major allele exhibiting the yellow phenotype. In contrast, only 0–27.3% of individuals homozygous for the minor allele were yellow. Among heterozygotes, 33.3–47.4% were yellow, reflecting an intermediate frequency of the yellow morph compared to the two homozygous groups. As ANSGD outputs posterior genotype probabilities rather than discrete calls, we were unable to perform formal tests of inheritance models, which depend on confidently assigned genotypes.

**Figure 5.**
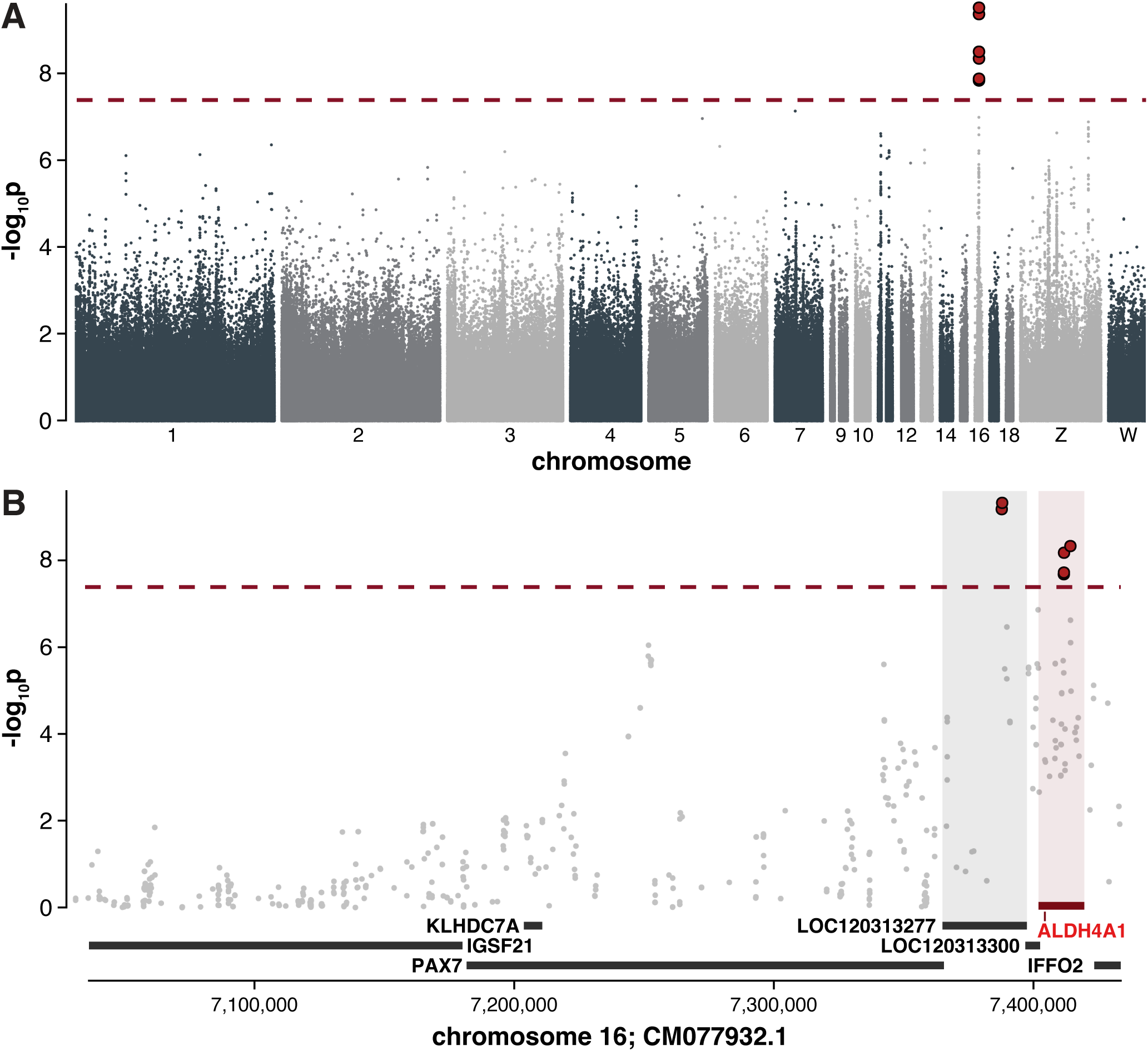
GWAS for color morph identifies a significantly associated locus on chromosome 16 overlapping a candidate gene. (A) Manhattan plot of genome-wide association results with the threshold for genome-wide significance marked by a dashed horizontal red line and all significantly associated variants shown as red circles. (B) The associated region on chromosome 16 shows that 4 of the 6 significant SNPs are in the gene ALDH4A1, with several other genes in the region.

### Microbiome diversity

Given the potential influence of microbial communities on skin health and disease susceptibility, we examined the microbiome of timber rattlesnake scale clips against two reference databases: one containing bacterial and viral genomes and another containing fungal genomes. Host reads were removed by aligning them to the *C. horridus* reference genome. Half of the non-host reads could be assigned to a taxon in the bacterial/viral database (50.8 ± 19.8%, range = 22.08– 98.7%; Table S14), and a much smaller percentage (on average, 4.3 ± 3.4%; range = 0.6– 30.3%) could be assigned using the fungal database.

We observed geographic structuring of the timber rattlesnake microbiome across the sampled populations. Samples from snakes from different populations were more different from one another than samples from the same population, as indicated by significantly lower Bray–Curtis distances (Wilcoxon signed-rank test, p = 6×10^-17^) (Figure S1A; Table S15). Individual snakes from different populations tend to more distinct microbiome compositions (PERMANOVA F = 7.45, R² = 0.313, p = 0.001) (Figure S1B), and some populations have more homogeneous microbiomes than others (PERMDISP F = 3.43, p = 0.006; Table S15) (Figure S1C). The alpha diversity of the microbiome, which measures within-sample richness and evenness, also differs between populations (Kruskal–Wallis χ²=21.504, p=0.0015; Figure S1D), and this result remained statistically significant in a restricted permutation test that randomized population labels 1,000 times while preserving sample sizes (permutation p<0.001; none as extreme as the observed value).

### Prevalence of snake fungal disease pathogen

Snakes with clinical evidence of Snake Fungal Disease carried higher sequencing read counts of the fungal pathogen *O. ophiodiicola*, suggesting that sequencing-based approaches could serve as a valuable tool for monitoring pathogen presence and population health. To quantify pathogen load, we mapped host-depleted metagenomic reads to the *O. ophiodiicola* reference genome and compared coverage-adjusted burden of *O. ophiodiicola* reads (“fungal burden”) to clinical evidence of infection, specifically the presence of skin lesions (Davy et al., 2021; Lorch et al., 2016). The overall distributions of fungal burden differed significantly between snakes with and without lesions (Kolmogorov–Smirnov test: D = 0.586, p = 0.022; Figure 6A; Table S16).

**Figure 6.**
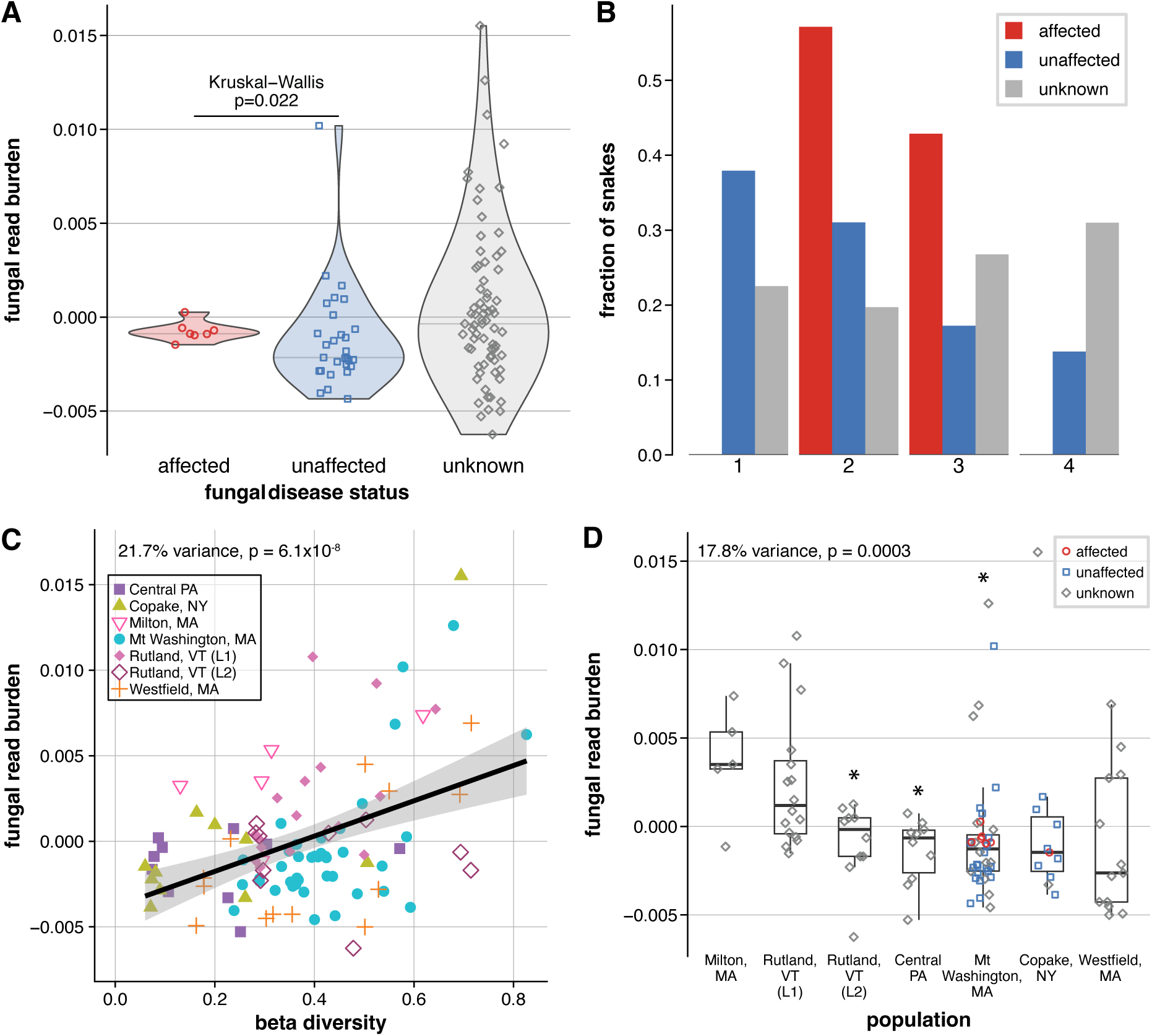
Snakes with clinical signs of Snake Fungal Disease have a higher burden of sequencing reads from the pathogen in scale-extracted DNA. (A) Fungal read burden differs among snakes with and without lesions consistent with Snake Fungal Disease. (B) Distribution of individuals across quartiles of fungal burden by status. Bar height indicates the fraction of snakes of a given disease status in each bin. (C) Fungal burden differs by population, but in a linear model that considers population, alpha diversity and beta diversity, only (D) beta diversity has a significant effect.

Snakes with lesions had a higher median fungal burden than snakes without lesions (Δmedian = 0.00127; one-sided, population-stratified permutation p = 0.029). Notably, none of the snakes with lesions fell within the lowest quartile of fungal burden (Figure 6B). Among snakes of unknown disease status (no information collected on presence or absence of lesions), 29 of 71 (41%) had burdens exceeding those observed in any snake with lesions.

Snakes with more unusual microbiome communities (greater β-diversity) tended to have higher *O. ophiodiicola* burdens. In a linear model including α-diversity, β-diversity, and population, these factors together explained approximately 40% of the variation in fungal burden (adjusted R² =0.349). β-diversity accounted for the largest portion of explained variance (21.7%) and was significantly and positively associated with fungal burden (β = 0.0131 ± 0.0022, p = 6.1×10^-8^; Figure 6C), indicating that snakes with microbiomes more distinct from others carried higher infection loads. Population explained an additional 17.8% of the variance (p = 3.0×10^-4^), with snakes from Rutland, VT (L2) (p=0.007), Mt. Washington, MA (p=0.003), and Westfield, MA (p=0.004) exhibiting lower fungal burdens than those from other sites (Figure 6D). In contrast, α-diversity (overall microbial diversity) contributed less than 1% of the explained variance and was not significantly associated with burden (p = 0.59).

## Discussion

We combined population genomics, trait association, and metagenomic profiling to examine the biogeography and conservation genetics of timber rattlesnakes (*Crotalus horridus*) in the Northeastern Appalachians. Using skim (∼1x) whole-genome sequencing of scale-clip extracted DNA, we simultaneously recovered genomic markers that permitted sex assignment, profiled host genetic variation, and characterized microbiome diversity. This untargeted approach demonstrates that a single skim-WGS library can yield valuable genetic and ecological information.

The value of untargeted sequencing increases when paired with individual and ecological metadata collected at the time of sampling. For example, for a subset of our snakes, color morph was recorded, enabling us to perform a GWAS that identified a locus on chromosome 16 containing *ALDH4A1*. Although this gene has not previously been linked to coloration in reptiles, it belongs to the aldehyde dehydrogenase family, which is functionally connected to pigment regulation in mammals and birds (Arbore et al., 2024; Kleszczynski & Slominski, 2013; Matsumoto et al., 2019). In snakes, pigmentation has typically been attributed to melanin pathway genes such as *TYR*, *TYRP1*, and *MC1R* (McNamara et al., 2021; Rosenblum et al., 2004). Our finding points to a possible alternative biochemical route, although further functional validation is required. The locus explains only part of the observed variation, consistent with polygenic architecture or environmental modulation (e.g., temperature-dependent expression).

By comparing sequencing coverage of the Z and W chromosomes, we were able to determine the genetic sex of each snake and assess the sex distribution in our populations. Sequencing-based sex determination is common in birds and has been used previously in snakes, but rarely in field-collected pit vipers. Instead, sexing often relies on imperfect morphology indicators (King, 1989; Rothe-Groleau et al., 2018) or invasive techniques such as probing or hemipenis eversion, which can stress or injure snakes (Fitch, 1960; Katz et al., 2020; Laszlo, 1975).

Consistent with previous work, we found no evidence of sex bias in timber rattlesnakes (Bauder et al., 2018; Brown & Maclean, 1983; Reinert & Zappalorti, 1988).

Sequencing is a particularly sensitive approach for detecting microbial pathogens. We detected intermediate or high levels of *O. ophiodiicola* DNA from scale clips in every snake with clinical signs of Snake Fungal Disease, highlighting the potential of untargeted genome sequencing for pathogen surveillance. This suggests scale clips could provide a less invasive alternative to tissue biopsies for detecting pathogen presence. While skin swabs are even less invasive and easier to collect (McKenzie et al., 2019), they often yield a lower quality and quantity of DNA, making it more difficult to detect the pathogen (Allender, Bunick, et al., 2015; Byrne et al., 2017). For example, only 37% of snakes with lesions tested positive for *O. ophiodiicola* by qPCR from swabs (Haynes et al., 2020), and culturing from swabs detected the pathogen in just 76% of symptomatic individuals (Lorch et al., 2016). The detection method rather than the sample type may be the key factor, as several studies have found no difference between scale clip and swab sampling(Baker & Allender, 2017; McKenzie et al., 2019). This raises the possibility that applying new sequencing technologies optimized for very low DNA yields could make swabs a reliable, fully noninvasive option for pathogen surveillance (Koda et al., 2023). Long-read sequencing would also help determine whether different *O. ophiodiicola* strains circulate among populations, and whether variation in strain distribution contributes to differences in disease severity (Koda et al., 2023).

We detected high levels of O. ophiodiicola in snakes without visible skin lesions, suggesting that untargeted sequencing may be a powerful and sensitive approach for monitoring pathogen prevalence and identifying subclinical infections. The high sensitivity of sequencing-based detection aligns with findings from environmental DNA and metagenomic studies in amphibians and fish (Hassan et al., 2022; Sahu et al., 2025). Consistent with earlier work showing that snakes with mild or no clinical signs often harbor lower pathogen loads (Allender, Bunick, et al., 2015; Haynes et al., 2020; Lorch et al., 2016; McKenzie et al., 2021), most snakes without lesions had lower fungal burdens than those with lesions. A few asymptomatic snakes, however, carried high pathogen loads, potentially reflecting infections that had not yet produced visible symptoms. Subclinical infections are known to occur (McKenzie et al., 2019), and disease severity is influenced by environmental conditions, with colder fall temperatures associated with more severe clinical signs (McCoy et al., 2017). Alternatively, O. ophiodiicola detected in asymptomatic snakes could reflect surface contamination, as DNA can leach into ethanol from external sources (Shokralla et al., 2010). Although our sampling protocol (a single scale clip per ethanol-filled tube) minimized cross-contamination, we cannot fully exclude the possibility of environmental or handling-related contamination. Histopathological examination would be necessary to confirm true infection.

Sequencing also captured the broader skin microbiome, allowing us to examine how fungal pathogen load relates to overall microbial community composition. Snakes with higher *O. ophiodiicola* loads tended to have higher β-diversity, indicating that their microbiome differs more from those of other snakes in this study. This pattern mirrors previous findings that *O. ophiodiicola* infection is associated with shifts in microbial community structure and increased variability among individuals (Romer et al., 2022, 2025; Walker et al., 2019). Together, these results suggest that infection disrupts the stability of the skin microbiome, potentially altering normal host–microbe interactions and leading to more heterogeneous microbial communities across individuals.

Our results highlight both challenges and opportunities for the conservation of timber rattlesnakes. On the positive side, the populations we surveyed still maintain reasonable levels of genetic diversity and show no evidence of extreme inbreeding, suggesting potential for long-term persistence under appropriate management. However, differentiation between geographically close populations, like the populations near Rutland VT, points to barriers that restrict dispersal and reduce gene flow, and extending this study to more relictual populations may reveal more severe inbreeding. Management strategies that enhance connectivity or consider translocations may help mitigate these risks, though such interventions may need to be balanced against the dangers of Snake Fungal Disease or other concerns.

Once widespread across the Northeastern U.S., timber rattlesnakes now persist only in fragmented refugia, where they face ongoing threats from habitat loss, human persecution, and Snake Fungal Disease. Our study demonstrates that untargeted sequencing, when paired with detailed ecological context, can generate rich insights about both hosts and pathogens from minimal tissue samples. This dual utility highlights the potential for integrating pathogen surveillance into population genomic studies, reducing costs and animal handling while increasing conservation impact. Moving forward, broader geographic and temporal sampling, coupled with comprehensive ecological data for each snake, will be essential for fully leveraging genomic technologies in the conservation and management of threatened rattlesnake populations.

## Data Availability

Sequence data have been deposited in the NCBI Sequence Read Archive under BioProject accession number PRJNA1346676. All associated data analysis files and metadata are available in the Dryad repository (DOI: 10.5061/dryad.m37pvmdg9).

## References

Adhikari, L., Shrestha, S., Wu, S., Crain, J., Gao, L., Evers, B., Wilson, D., Ju, Y., Koo, D.-H., Hucl, P., Pozniak, C., Walkowiak, S., Wang, X., Wu, J., Glaubitz, J. C., DeHaan, L., Friebe, B., & Poland, J. (2022). A high-throughput skim-sequencing approach for genotyping, dosage estimation and identifying translocations. Scientific Reports, 12(1), 17583. 10.1038/s41598-022-19858-2

Allender, M. C., Baker, S., Britton, M., & Kent, A. D. (2018). Snake fungal disease alters skin bacterial and fungal diversity in an endangered rattlesnake. Scientific Reports, 8(1), 12147. 10.1038/s41598-018-30709-x

Allender, M. C., Bunick, D., Dzhaman, E., Burrus, L., & Maddox, C. (2015). Development and use of a real-time polymerase chain reaction assay for the detection of Ophidiomyces ophiodiicola in snakes. *Journal of Veterinary Diagnostic Investigation: Official Publication of the American Association of Veterinary Laboratory Diagnosticians*, Inc, 27(2), 217–220. 10.1177/1040638715573983

Allender, M. C., Raudabaugh, D. B., Gleason, F. H., & Miller, A. N. (2015). The natural history, ecology, and epidemiology of Ophidiomyces ophiodiicola and its potential impact on free-ranging snake populations. Fungal Ecology, 17, 187–196. 10.1016/j.funeco.2015.05.003

Arbore, R., Barbosa, S., Brejcha, J., Ogawa, Y., Liu, Y., Nicolaï, M. P. J., Pereira, P., Sabatino, S. J., Cloutier, A., Poon, E. S. K., Marques, C. I., Andrade, P., Debruyn, G., Afonso, S., Afonso, R., Roy, S. G., Abdu, U., Lopes, R. J., Mojzeš, P.,… Carneiro, M. (2024). A molecular mechanism for bright color variation in parrots. *Science (New York*, N.Y.), 386(6721), eadp7710. 10.1126/science.adp7710

Baker, S., & Allender, M. (2017). Comparison of testing methods for snake fungal disease.

Bauder, J. M., Stengle, A. G., Jones, M., Marchand, M., Blodgett, D., Hess, B., Clifford, B., & Jenkins, C. L. (2018). Population Demographics, Monitoring, and Population Genetics of Timber Rattlesnakes in New England. Regional Conservation Needs (RCN) program. https://rcngrants.org/sites/default/files/final_reports/WMI_Timber_Rattlesnake_Report_final_2_9_18.pdf

Brown, W. S., & Maclean, F. M. (1983). Conspecific scent trailing by new born timber rattlesnake crotalus horridus. Herpetologica, 39(4), 430–436. https://www.jstor.org/stable/3892539?seq=1

Bushar, L. M., Bhatt, N., Dunlop, M. C., Schocklin, C., Malloy, M. A., & Reinert, H. K. (2015). Population Isolation and Genetic Subdivision of Timber Rattlesnakes (Crotalus horridus) in the New Jersey Pine Barrens. Herpetologica, 71(3), 203–211. 10.1655/HERPETOLOGICA-D-14-00030

Byrne, A. Q., Rothstein, A. P., Poorten, T. J., Erens, J., Settles, M. L., & Rosenblum, E. B. (2017). Unlocking the story in the swab: A new genotyping assay for the amphibian chytrid fungus Batrachochytrium dendrobatidis. Molecular Ecology Resources, 17(6), 1283–1292. 10.1111/1755-0998.12675

Ceballos, F. C., Joshi, P. K., Clark, D. W., Ramsay, M., & Wilson, J. F. (2018). Runs of homozygosity: windows into population history and trait architecture. Nature Reviews. Genetics, 19(4), 220–234. 10.1038/nrg.2017.109

Chat, V., Ferguson, R., Morales, L., & Kirchhoff, T. (2021). Ultra low-coverage whole-genome sequencing as an alternative to genotyping arrays in genome-wide association studies. Frontiers in Genetics, 12, 790445. 10.3389/fgene.2021.790445

Chen, K., Knorr, C., Bornemann-Kolatzki, K., Ren, J., Huang, L., Rohrer, G. A., & Brenig, B. (2005). Targeted oligonucleotide-mediated microsatellite identification (TOMMI) from large-insert library clones. BMC Genetics, 6(1), 54. 10.1186/1471-2156-6-54

Clark, R. W., Brown, W. S., Stechert, R., & Zamudio, K. R. (2010). Roads, interrupted dispersal, and genetic diversity in timber rattlesnakes. Conservation Biology: The Journal of the Society for Conservation Biology, 24(4), 1059–1069. 10.1111/j.1523-1739.2009.01439.x

Clark, R. W., Marchand, M. N., Clifford, B. J., Stechert, R., & Stephens, S. (2011). Decline of an isolated timber rattlesnake (*Crotalus horridus*) population: Interactions between climate change, disease, and loss of genetic diversity. Biological Conservation, 144(2), 886–891. 10.1016/j.biocon.2010.12.001

COSEWIC. (2023). Rapid Review of Classification on the Timber Rattlesnake Crotalus horridus in Canada. Committee on the Status of Endangered Wildlife in Canada. https://www.canada.ca/en/environment-climate-change/services/species-risk-public-registry/cosewic-assessments-status-reports/timber-rattlesnake-2023.html

Davy, C. M., Shirose, L., Campbell, D., Dillon, R., McKenzie, C., Nemeth, N., Braithwaite, T., Cai, H., Degazio, T., Dobbie, T., Egan, S., Fotherby, H., Litzgus, J. D., Manorome, P., Marks, S., Paterson, J. E., Sigler, L., Slavic, D., Slavik, E.,… Jardine, C. (2021). Revisiting ophidiomycosis (snake fungal disease) after a decade of targeted research. Frontiers in Veterinary Science, 8, 665805. 10.3389/fvets.2021.665805

Eckert, S. A., & Jesper, A. C. (2024). Home range, site fidelity, and movements of timber rattlesnakes (Crotalus horridus) in west-central Illinois. Animal Biotelemetry, 12(1), 1–16. 10.1186/s40317-023-00357-8

Fitch, H. S. (1960). Criteria for determining sex and breeding maturity in snakes. Herpetologica, 16, 49–51.

Fuentes-Pardo, A. P., & Ruzzante, D. E. (2017). Whole-genome sequencing approaches for conservation biology: Advantages, limitations and practical recommendations. Molecular Ecology, 26(20), 5369–5406. 10.1111/mec.14264

Furman, J. (2007). Timber rattlesnakes in Vermont & New York: Biology, history, and the fate of an endangered species. https://books.google.com/books?hl=en&lr=&id=CdOzBQAAQBAJ&oi=fnd&pg=PP1&dq=timber+rattlesnake+endangered&ots=RNLsxowrkB&sig=_6TCRbqgUIUsW7AT28Xb8XPD2Ns

Gibbs, H. L., Prior, K., & Parent, C. (1998). Brief communication. Characterization of DNA microsatellite loci from a threatened snake: the eastern massasauga rattlesnake (Sistrurus c. catenatus and their use in population studies. The Journal of Heredity, 89(2), 169–173. 10.1093/jhered/89.2.169

Hassan, S., Sabreena, Poczai, P., Ganai, B. A., Almalki, W. H., Gafur, A., & Sayyed, R. Z. (2022). Environmental DNA metabarcoding: A novel contrivance for documenting terrestrial biodiversity. Biology, 11(9), 1297. 10.3390/biology11091297

Haynes, E., Chandler, H. C., Stegenga, B. S., Adamovicz, L., Ospina, E., Zerpa-Catanho, D., Stevenson, D. J., & Allender, M. C. (2020). Ophidiomycosis surveillance of snakes in Georgia, USA reveals new host species and taxonomic associations with disease. Scientific Reports, 10(1), 10870. 10.1038/s41598-020-67800-1

Heppell, S. S., Caswell, H., & Crowder, L. B. (2000). Life histories and elasticity patterns: Perturbation analysis for species with minimal demographic data. Ecology, 81(3), 654–665. 10.1890/0012-9658(2000)081[0654:LHAEPP]2.0.CO;2

Hill, W. G., Goddard, M. E., & Visscher, P. M. (2008). Data and theory point to mainly additive genetic variance for complex traits. PLoS Genetics, 4(2), e1000008. 10.1371/journal.pgen.1000008

Hohenlohe, P. A., Funk, W. C., & Rajora, O. P. (2021). Population genomics for wildlife conservation and management. Molecular Ecology, 30(1), 62–82. 10.1111/mec.15720

Holycross, A. T., Douglas, M. E., Higbee, J. R., & Bogden, R. H. (2002). Isolation and characterization of microsatellite loci from a threatened rattlesnake (New Mexico RidgeLJnosed Rattlesnake, Crotalus willardi obscurus). Molecular Ecology Notes, 2(4), 537–539. 10.1046/j.1471-8286.2002.00310.x

Homburger, J. R., Neben, C. L., Mishne, G., Zhou, A. Y., Kathiresan, S., & Khera, A. V. (2019). Low coverage whole genome sequencing enables accurate assessment of common variants and calculation of genome-wide polygenic scores. Genome Medicine, 11(1), 74. 10.1186/s13073-019-0682-2

Iannucci, A., Benazzo, A., Natali, C., Arida, E. A., Zein, M. S. A., Jessop, T. S., Bertorelle, G., & Ciofi, C. (2021). Population structure, genomic diversity and demographic history of Komodo dragons inferred from whole-genome sequencing. Molecular Ecology, 30(23), 6309–6324. 10.1111/mec.16121

Jørsboe, E., & Albrechtsen, A. (2022). Efficient approaches for large-scale GWAS with genotype uncertainty. G3, 12(1), jkab385. 10.1093/g3journal/jkab385

Katz, A. D., Pearce, S., Melder, C., Sperry, J. H., & Davis, M. A. (2020). Molecular sexing is a viable alternative to probing for determining sex in the imperiled Louisiana Pine Snake (Pituophis ruthveni). Conservation Genetics Resources, 12(4), 537–539. 10.1007/s12686-020-01145-9

Keyler, D. E. (2023). Timber rattlesnake (Crotalus horridus): Biology, conservation, and envenomation in the Upper Mississippi River Valley (1982-2020). Toxicon: X, 19(100167), 100167. 10.1016/j.toxcx.2023.100167

King, R. B. (1989). Sexual dimorphism in snake tail length: sexual selection, natural selection, or morphological constraint? Biological Journal of the Linnean Society, 38(2), 133–154. 10.1111/j.1095-8312.1989.tb01570.x

Kleszczynski, K., & Slominski, A. T. (2013). Targeting ALDH1A1 to treat pigmentary disorders. Experimental Dermatology, 22(5), 316–317. 10.1111/exd.12134

Koda, S. A., McCauley, M., Farrell, J. A., Duffy, I. J., Duffy, F. G., Loesgen, S., Whilde, J., & Duffy, D. J. (2023). A novel eDNA approach for rare species monitoring: Application of long-read shotgun sequencing to Lynx rufus soil pawprints. Biological Conservation, 287(110315), 110315. 10.1016/j.biocon.2023.110315

Korneliussen, T. S., Albrechtsen, A., & Nielsen, R. (2014). ANGSD: Analysis of Next Generation Sequencing Data. BMC Bioinformatics, 15(1), 356. 10.1186/s12859-014-0356-4

Kumar, S., Suleski, M., Craig, J. M., Kasprowicz, A. E., Sanderford, M., Li, M., Stecher, G., & Hedges, S. B. (2022). TimeTree 5: An expanded resource for species divergence times. Molecular Biology and Evolution, 39(8), msac174. 10.1093/molbev/msac174

Kyriazis, C. C., Robinson, J. A., & Lohmueller, K. E. (2025). Long runs of homozygosity are reliable genomic markers of inbreeding depression. Trends in Ecology & Evolution. 10.1016/j.tree.2025.06.013

Ladner, J. T., Palmer, J. M., Ettinger, C. L., Stajich, J. E., Farrell, T. M., Glorioso, B. M., Lawson, B., Price, S. J., Stengle, A. G., Grear, D. A., & Lorch, J. M. (2022). The population genetics of the causative agent of snake fungal disease indicate recent introductions to the USA. PLoS Biology, 20(6), e3001676. 10.1371/journal.pbio.3001676

Laszlo, J. (1975). Probing as a practical method of sex recognition in snakes. The International Zoo Yearbook, 15(1), 178–179. 10.1111/j.1748-1090.1975.tb01393.x

Lind, C. M., Agugliaro, J., Lorch, J. M., & Farrell, T. M. (2023). Ophidiomycosis is related to seasonal patterns of reproduction, ecdysis, and thermoregulatory behavior in a freeLJliving snake species. Journal of Zoology (London, England: 1987), 319(1), 54–62. 10.1111/jzo.13024

Li, S., Yan, B., Li, T. K. T., Lu, J., Gu, Y., Tan, Y., Gong, F., Lam, T.-W., Xie, P., Wang, Y., Lin, G., & Luo, R. (2023). Ultra-low-coverage genome-wide association study-insights into gestational age using 17,844 embryo samples with preimplantation genetic testing. Genome Medicine, 15(1), 10. 10.1186/s13073-023-01158-7

Lorch, J. M., Knowles, S., Lankton, J. S., Michell, K., Edwards, J. L., Kapfer, J. M., Staffen, R. A., Wild, E. R., Schmidt, K. Z., Ballmann, A. E., Blodgett, D., Farrell, T. M., Glorioso, B. M., Last, L. A., Price, S. J., Schuler, K. L., Smith, C. E., Wellehan, J. F. X., Jr, & Blehert, D. S. (2016). Snake fungal disease: an emerging threat to wild snakes. Philosophical Transactions of the Royal Society of London. Series B, Biological Sciences, 371(1709). 10.1098/rstb.2015.0457

Lorch, J. M., Lankton, J., Werner, K., Falendysz, E. A., McCurley, K., & Blehert, D. S. (2015). Experimental infection of snakes with Ophidiomyces ophiodiicola causes pathological changes that typify snake fungal disease. mBio, 6(6), e01534–15. 10.1128/mBio.01534-15

Lou, R. N., Jacobs, A., Wilder, A. P., & Therkildsen, N. O. (2021). A beginner’s guide to low-coverage whole genome sequencing for population genomics. Molecular Ecology, 30(23), 5966–5993. 10.1111/mec.16077

Lu, J., Breitwieser, F. P., Thielen, P., & Salzberg, S. L. (2017). Bracken: estimating species abundance in metagenomics data. PeerJ. Computer Science, 3(e104), e104. 10.7717/peerj-cs.104

Margres, M. J., Rautsaw, R. M., Strickland, J. L., Mason, A. J., Schramer, T. D., Hofmann, E. P., Stiers, E., Ellsworth, S. A., Nystrom, G. S., Hogan, M. P., Bartlett, D. A., Colston, T. J., Gilbert, D. M., Rokyta, D. R., & Parkinson, C. L. (2021). The Tiger Rattlesnake genome reveals a complex genotype underlying a simple venom phenotype. Proceedings of the National Academy of Sciences, 118(4), e2014634118. 10.1073/pnas.2014634118

Martin, M. (2011). Cutadapt removes adapter sequences from high-throughput sequencing reads. EMBnet.journal, 17(1), 10–12. 10.14806/ej.17.1.200

Martin, W. H., Brown, W. S., Possardt, E. E., & Sealy, J. B. (2008). Biological variation, management units, and a conservation plan for the timber rattlesnake (Crotalus horridus). In W. K. Hayes, K. R. Beaman, M. D. Cardwell, & S. P. Bush (Eds.), The Biology of Rattlesnakes (pp. 447–462). Loma Linda University Press.

Matsumoto, A., Ito, S., Wakamatsu, K., Ichiba, M., Vasiliou, V., Akao, C., Song, B.-J., & Fujita, M. (2019). Ethanol induces skin hyperpigmentation in mice with aldehyde dehydrogenase 2 deficiency. Chemico-Biological Interactions, 302, 61–66. 10.1016/j.cbi.2019.01.035

McCoy, C. M., Lind, C. M., & Farrell, T. M. (2017). Environmental and physiological correlates of the severity of clinical signs of snake fungal disease in a population of pigmy rattlesnakes, Sistrurus miliarius. Conservation Physiology, 5(1), cow077. 10.1093/conphys/cow077

McKenna, A., Hanna, M., Banks, E., Sivachenko, A., Cibulskis, K., Kernytsky, A., Garimella, K., Altshuler, D., Gabriel, S., Daly, M., & DePristo, M. A. (2010). The Genome Analysis Toolkit: A MapReduce framework for analyzing next-generation DNA sequencing data. Genome Research, 20(9), 1297–1303. 10.1101/gr.107524.110

McKenzie, J. M., Price, S. J., Connette, G. M., Bonner, S. J., & Lorch, J. M. (2021). Effects of snake fungal disease on short-term survival, behavior, and movement in free-ranging snakes. Ecological Applications: A Publication of the Ecological Society of America, 31(2), e02251. 10.1002/eap.2251

McKenzie, J. M., Price, S. J., Fleckenstein, J. L., Drayer, A. N., Connette, G. M., Bohuski, E., & Lorch, J. M. (2019). Field diagnostics and seasonality of Ophidiomyces ophiodiicola in wild snake populations. EcoHealth, 16(1), 141–150. 10.1007/s10393-018-1384-8

McNamara, M. E., Rossi, V., Slater, T. S., Rogers, C. S., Ducrest, A.-L., Dubey, S., & Roulin, A. (2021). Decoding the evolution of melanin in vertebrates. Trends in Ecology & Evolution, 36(5), 430–443. 10.1016/j.tree.2020.12.012

Meisner, J., & Albrechtsen, A. (2018). Inferring Population Structure and Admixture Proportions in Low-Depth NGS Data. Genetics, 210(2), 719–731. 10.1534/genetics.118.301336

Melville, J. (2025). uwot: An R package implementing the UMAP dimensionality reduction method (Version 0.2.3). Github. https://github.com/jlmelville/uwot

Montague, G. (2022). Head-Starting and Conservation of Endangered Timber Rattlesnakes (Crotalus horridus horridus) at Roger Williams Park Zoo. Journal of Zoological and Botanical Gardens, 3(4), 581–585. 10.3390/jzbg3040043

Ochoa, A., & Gibbs, H. L. (2021). Genomic signatures of inbreeding and mutation load in a threatened rattlesnake. Molecular Ecology, 30(21), 5454–5469. 10.1111/mec.16147

Oksanen, J., Simpson, G. L., Blanchet, F. G., Kindt, R., Legendre, P., Minchin, P. R., O’Hara, R. B., Solymos, P., Stevens, M. H. H., Szoecs, E., Wagner, H., Barbour, M., Bedward, M., Bolker, B., Borcard, D., Borman, T., Carvalho, G., Chirico, M., De Caceres, M.,… Weedon, J. (2025). vegan: Community Ecology Package (Versions 2.7-1). https://CRAN.R-project.org/package=vegan

Reinert, H. K., & Zappalorti, R. T. (1988). Timber rattlesnakes (Crotalus horridus) of the pine barrens: Their movement patterns and habitat preference. Copeia, 1988(4), 964. 10.2307/1445720

Rhoads, D., Pummill, J., Beaupre, S. J., & Sanders, W. S. (2024). Crotalus horridus genome assembly ASM162548v2 [Dataset]. In Timber rattlesnake genome. https://www.ncbi.nlm.nih.gov/datasets/genome/GCA_001625485.2

Romer, A. S., Grinath, J. B., Moe, K. C., & Walker, D. M. (2022). Host microbiome responses to the Snake Fungal Disease pathogen (Ophidiomyces ophidiicola) are driven by changes in microbial richness. Scientific Reports, 12(1), 3078. 10.1038/s41598-022-07042-5

Romer, A. S., Grisnik, M., Dallas, J. W., Sutton, W., Murray, C. M., Hardman, R. H., Blanchard, T., Hanscom, R. J., Clark, R. W., Godwin, C., Alexander, N. R., Moe, K. C., Cobb, V. A., Eaker, J., Colvin, R., Thames, D., Ogle, C., Campbell, J., Frost, C.,… Walker, D. M. (2025). Effects of snake fungal disease (ophidiomycosis) on the skin microbiome across two major experimental scales. Conservation Biology: The Journal of the Society for Conservation Biology, 39(2), e14411. 10.1111/cobi.14411

Roseman, M. A., Mason, A. J., Bode, E. R., Bolton, P. E., Nachtigall, P. G., Peterman, W. E., & Gibbs, H. L. (2025). Insights from the timber rattlesnake (Crotalus horridus) genome for MHC gene architecture and evolution in threatened rattlesnakes. The Journal of Heredity, 116(5), 591–602. 10.1093/jhered/esae075

Rosenblum, E. B., Hoekstra, H. E., & Nachman, M. W. (2004). Adaptive reptile color variation and the evolution of the Mc1r gene. Evolution; International Journal of Organic Evolution, 58(8), 1794–1808. 10.1111/j.0014-3820.2004.tb00462.x

Rothe-Groleau, C., Rauter, C. M., & Fawcett, J. D. (2018). Morphological traits as indicators of sexual dimorphism in Prairie Rattlesnakes (Crotalus viridis). Transactions of the Nebraska Academy of Sciences and Affiliated Societies. 10.13014/k2x63k4g

Sahu, A., Singh, M., Amin, A., Malik, M. M., Qadri, S. N., Abubakr, A., Teja, S. S., Dar, S. A., & Ahmad, I. (2025). A systematic review on environmental DNA (eDNA) Science: An eco-friendly survey method for conservation and restoration of fragile ecosystems. Ecological Indicators, 173(113441), 113441. 10.1016/j.ecolind.2025.113441

Shafer, A. B. A., & Kardos, M. (2025). Runs of homozygosity and inferences in wild populations. Molecular Ecology, 34(3), e17641. 10.1111/mec.17641

Shokralla, S., Singer, G. A. C., & Hajibabaei, M. (2010). Direct PCR amplification and sequencing of specimens’ DNA from preservative ethanol. BioTechniques, 48(3), 233–234. 10.2144/000113362

Shumate, A., & Salzberg, S. L. (2021). Liftoff: accurate mapping of gene annotations. *Bioinformatics (Oxford*, England), 37(12), 1639–1643. 10.1093/bioinformatics/btaa1016

Stengle, A. (2018). Habitat selection, connectivity, and population genetics of a timber rattlesnake (Crotalus horridus) metapopulation in southwestern Massachusetts and New England (P. R. Sievert (ed.)) [PhD, University of Massachusetts Amherst]. 10.7275/11242468.0

Stengle, A. G., Farrell, T. M., Freitas, K. S., Lind, C. M., Price, S. J., Butler, B. O., Tadevosyan, T., Isidoro-Ayza, M., Taylor, D. R., Winzeler, M., & Lorch, J. M. (2019). Evidence of vertical transmission of the snake fungal pathogen Ophidiomyces ophiodiicola. Journal of Wildlife Diseases, 55(4), 961–964. 10.7589/2018-10-250

Sun, L., Wang, Z., Lu, T., Manolio, T. A., & Paterson, A. D. (2023). eXclusionarY: 10 years later, where are the sex chromosomes in GWASs? The American Journal of Human Genetics, 110(6), 903–912. 10.1016/j.ajhg.2023.04.009

Theissinger, K., Fernandes, C., Formenti, G., Bista, I., Berg, P. R., Bleidorn, C., Bombarely, A., Crottini, A., Gallo, G. R., Godoy, J. A., Jentoft, S., Malukiewicz, J., Mouton, A., Oomen, R. A., Paez, S., Palsbøll, P. J., Pampoulie, C., Ruiz-López, M. J., Secomandi, S,… European Reference Genome Atlas Consortium. (2023). How genomics can help biodiversity conservation. Trends in Genetics: TIG, 39(7), 545–559. 10.1016/j.tig.2023.01.005

Villarreal, X., Bricker, J., Reinert, H. K., Gelbert, L., & Bushar, L. M. (1996). Isolation and characterization of microsatellite loci for use in population genetic analysis in the timber rattlesnake, Crotalus horridus. The Journal of Heredity, 87(2), 152–155. 10.1093/oxfordjournals.jhered.a022973

Walker, D. M., Leys, J. E., Grisnik, M., Grajal-Puche, A., Murray, C. M., & Allender, M. C. (2019). Variability in snake skin microbial assemblages across spatial scales and disease states. The ISME Journal, 13(9), 2209–2222. 10.1038/s41396-019-0416-x

Wickham, H. (2016). ggplot2: Elegant Graphics for Data Analysis (2nd ed. 2016). Springer International Publishing: Imprint: Springer.

Wilkinson, L. (2011). ggplot2: Elegant Graphics for Data Analysis by WICKHAM, H. Biometrics, 67(2), 678–679. 10.1111/j.1541-0420.2011.01616.x

Wood, D. E., Lu, J., & Langmead, B. (2019). Improved metagenomic analysis with Kraken 2. Genome Biology, 20(1), 257. 10.1186/s13059-019-1891-0

Xie, H.-X., Liang, X.-X., Chen, Z.-Q., Li, W.-M., Mi, C.-R., Li, M., Wu, Z.-J., Zhou, X.-M., & Du, W.-G. (2022). Ancient demographics determine the effectiveness of genetic purging in endangered lizards. Molecular Biology and Evolution, 39(1), msab359. 10.1093/molbev/msab359

