## Supplemental Files for "Population genetics, trait mapping and fungal pathogen surveillance using untargeted sequencing in timber rattlesnakes (*Crotalus horridus)*"

**Figure S1. Microbiome diversity and dissimilarity across populations.**

(A) Paired comparison showing between-population distances exceed within-population distances (Wilcoxon signed-rank, $p=1.3\times10^{-16}$). (B) Within-population Bray–Curtis dissimilarities by population (PERMDISP p=0.001). (C) Between-population Bray–Curtis dissimilarities with overall community separation (PERMANOVA p=0.001). (D) Shannon diversity index by population (boxplots show median and IQR; Kruskal–Wallis, $p=7\times10^{-4}$).

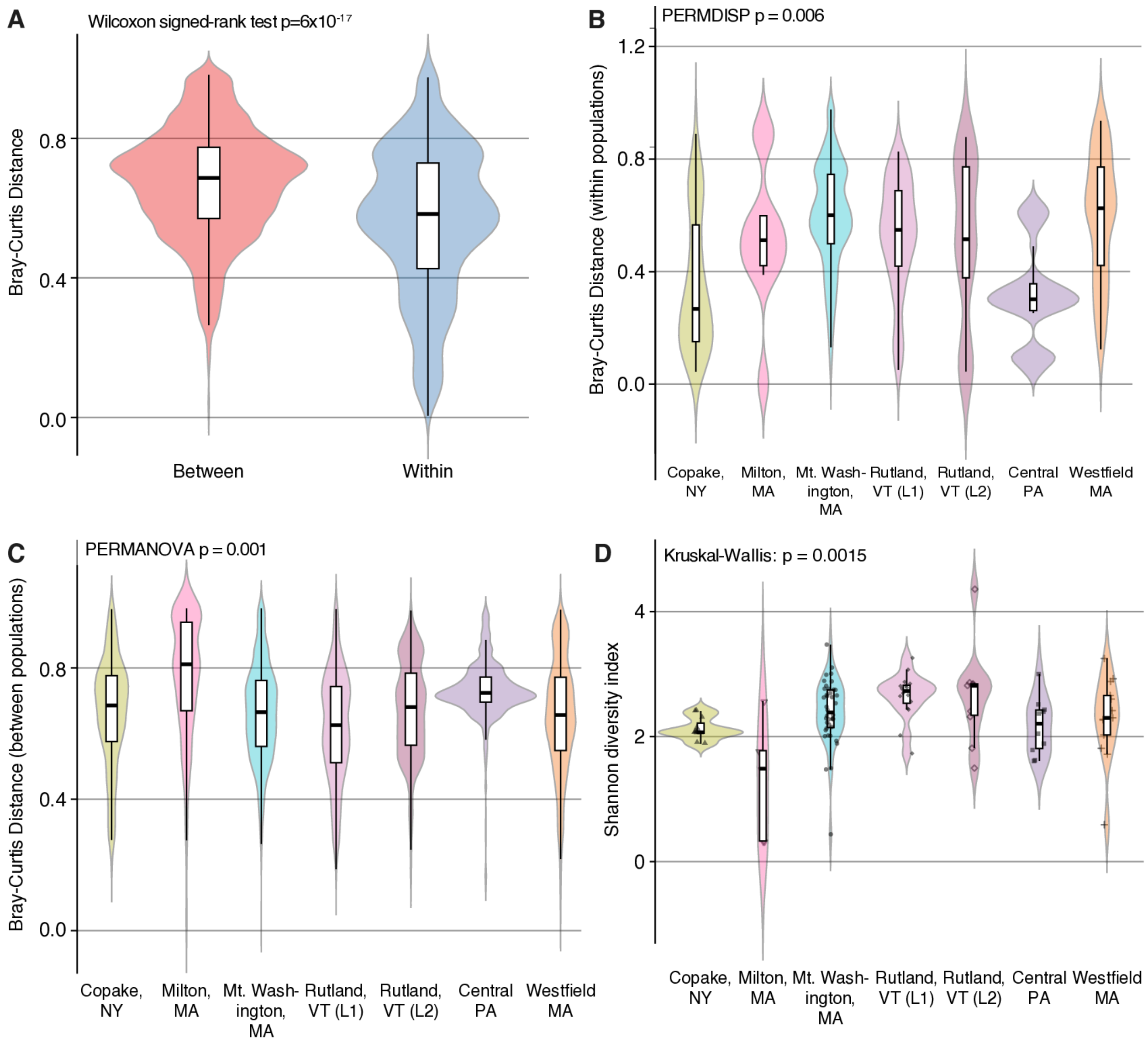

### Table S1. Sample metadata

Metadata for all sequenced timber rattlesnakes (n=111), including field ID ("snake"), color morph, population, sex proxy (WZ_ratio), and inferred sex, SFD status, and sequencing yield (Total_Gbp, Mean_Read_Length_bp).

| snake | Color | population | precise site | W/Z ratio | Inferred sex | lesions | Total Gbp | Mean Read Length (bp) |
|---|---|---|---|---|---|---|---|---|
| 09.06SFD | Y | Mt Washington MA | L4 | 0.085 | Male | Y | 4.9200 | 148.8 |
| 1 | NA | Central PA | NA | 0.485 | Female | no info | 5.4300 | 148.7 |
| 10 | NA | Central PA | NA | 0.510 | Female | no info | 3.0100 | 148.5 |
| 10.16 | Y | Mt Washington MA | L1 | 0.507 | Female | N | 4.2600 | 148.5 |
| 10.17 | B | Mt Washington MA | L1 | 0.116 | Male | N | 2.1800 | 148.3 |
| 10.21 | Y | Mt Washington MA | L1 | 0.110 | Male | Y | 2.2600 | 148.4 |
| 10.34 | B | Mt Washington MA | L1 | 0.400 | Female | N | 1.9300 | 148.4 |
| 10.4 | B | Copake NY | NA | 0.083 | Male | N | 5.7800 | 148.5 |
| 11 | NA | Central PA | NA | 0.102 | Male | no info | 0.5800 | 148.2 |
| 11.04 | Y | Mt Washington MA | L1 | 0.093 | Male | N | 3.7700 | 148.4 |
| 11.07 | B | Mt Washington MA | L4 | 0.084 | Male | Y | 4.4700 | 148.4 |
| 11.1 | Y | Mt Washington MA | L1 | 0.091 | Male | N | 0.6300 | 148.1 |
| 11.24 | B | Mt Washington MA | L1 | 0.093 | Male | N | 3.6500 | 148.4 |
| 11.28 | B | Mt Washington MA | L2 | 0.088 | Male | N | 5.5800 | 148.8 |
| 11.29 | Y | Mt Washington MA | L1 | 0.145 | Male | N | 4.3500 | 148.7 |
| 11.3 | Y | Mt Washington MA | L2 | 0.482 | Female | no info | 4.8500 | 148.4 |
| 11.31 | Y | Copake NY | NA | 0.432 | Female | N | 6.2000 | 148.8 |
| 11.32 | Y | Mt Washington MA | L2 | 0.457 | Female | N | 5.2400 | 148.4 |
| 11.35 | Y | Mt Washington MA | L2 | 0.142 | Male | no info | 5.2700 | 148.8 |
| 11.36 | Y | Mt Washington MA | L2 | 0.089 | Male | N | 4.4600 | 148.7 |
| 11.38 | Y | Mt Washington MA | L2 | 0.087 | Male | N | 5.1200 | 148.5 |
| 11.4 | Y | Mt Washington MA | L2 | 0.472 | Female | no info | 6.0500 | 148.5 |
| 11.41 | Y | Mt Washington MA | L2 | 0.087 | Male | Y | 6.0600 | 148.8 |
| 11.43 | B | Mt Washington MA | L1 | 0.489 | Female | N | 4.0400 | 148.4 |
| 11.44 | B | Copake NY | NA | 0.439 | Female | N | 5.8000 | 148.5 |
| 11.45 | B | Mt Washington MA | L4 | 0.486 | Female | N | 5.2300 | 148.4 |
| 11.46 | Y | Mt Washington MA | L2 | 0.091 | Male | N | 6.4300 | 148.6 |
| 11.52 | B | Copake NY | NA | 0.085 | Male | N | 5.9100 | 148.8 |
| 11.53 | B | Copake NY | NA | 0.417 | Female | N | 4.1700 | 148.4 |
| 11.54 | Y | Copake NY | NA | 0.107 | Male | no info | 5.9500 | 148.6 |
| 11.57 | Y | Copake NY | NA | 0.087 | Male | Y | 6.3200 | 148.6 |
| 11.58 | B | Copake NY | NA | 0.548 | Female | N | 2.3400 | 148.8 |
| 11.64 | B | Mt Washington MA | L4 | 0.117 | Male | N | 2.3800 | 148.3 |
| 11.68 | B | Copake NY | NA | 0.090 | Male | no info | 18.8100 | 148.9 |
| 11.73 | NA | Rockland County NY | L2 | 0.119 | Male | no info | 1.4400 | 148.4 |
| 11.74 | NA | Rockland County NY | L1 | 0.561 | Female | no info | 3.7300 | 148.4 |
| 12.11 | Y | Mt Washington MA | L3 | 0.584 | Female | no info | 1.8000 | 148.4 |
| 12.19 | B | Mt Washington MA | L3 | 0.559 | Female | no info | 4.3000 | 148.4 |
| 12.21 | B | Mt Washington MA | L4 | 0.088 | Male | Y | 4.3700 | 148.8 |
| 12.23 | Y | Mt Washington MA | L4 | 0.124 | Male | N | 5.2000 | 148.8 |
| 12.25 | Y | Mt Washington MA | L4 | 0.458 | Female | N | 4.3800 | 148.4 |
| 12.26 | B | Mt Washington MA | L3 | 0.441 | Female | no info | 4.5100 | 148.4 |
| 12.32 | Y | Mt Washington MA | L4 | 0.102 | Male | N | 4.2800 | 148.7 |
| 12.34 | Y | Mt Washington MA | L1 | 0.101 | Male | Y | 1.9200 | 148.6 |
| 12.39 | B | Mt Washington MA | L3 | 0.091 | Male | no info | 5.5500 | 148.4 |
| 12.4 | B | Mt Washington MA | L3 | 0.108 | Male | no info | 4.7200 | 148.8 |
| 12.43 | B | Mt Washington MA | L3 | 0.087 | Male | no info | 4.9200 | 148.5 |
| 12.48 | Y | Mt Washington MA | L2 | 0.084 | Male | N | 2.9400 | 148.2 |
| 12.49 | B | Mt Washington MA | L3 | 0.423 | Female | N | 1.7000 | 148.6 |
| 12.5 | B | Mt Washington MA | L3 | 0.469 | Female | no info | 0.6100 | 148.0 |
| 12.51 | B | Mt Washington MA | L3 | 0.216 | Male | no info | 0.0600 | 148.0 |
| 12.52 | Y | Mt Washington MA | L3 | 0.531 | Female | no info | 3.6200 | 148.7 |
| 12.61 | B | Copake NY | NA | 0.102 | Male | N | 4.7400 | 148.5 |
| 12.64 | B | Mt Washington MA | L4 | 0.552 | Female | N | 4.4700 | 148.6 |
| 12.71 | Y | Copake NY | NA | 0.084 | Male | N | 4.8200 | 148.8 |
| 12.73 | NA | Westfield MA | NA | 0.474 | Female | no info | 10.2200 | 148.7 |
| 12.74 | NA | Westfield MA | NA | 0.276 | Male | no info | 1.0700 | 148.8 |
| 12.75 | NA | Westfield MA | NA | 0.097 | Male | no info | 1.6700 | 148.5 |
| 162019 | NA | Milton MA | NA | 0.080 | Male | no info | 4.9700 | 148.6 |
| 186035 | NA | Milton MA | NA | 0.433 | Female | no info | 0.2300 | 147.4 |
| 186036 | NA | Milton MA | NA | 0.488 | Female | no info | 0.2300 | 148.0 |
| 186037 | NA | Milton MA | NA | 0.099 | Male | no info | 0.1700 | 147.3 |
| 19192 | NA | Milton MA | NA | 0.088 | Male | no info | 7.7900 | 148.7 |
| 2 | NA | Central PA | NA | 0.094 | Male | no info | 3.3900 | 148.3 |
| 27544 | NA | Mt Washington MA (1928) | Historic | 0.087 | Male | no info | 5.3300 | 148.6 |
| 3 | NA | Central PA | NA | 0.457 | Female | no info | 3.8400 | 148.7 |
| 5 | NA | Central PA | NA | 0.085 | Male | no info | 4.4800 | 148.4 |
| 6 | NA | Central PA | NA | 0.090 | Male | no info | 3.8900 | 148.4 |
| 6439 | NA | Ipswich MA (1896) | Historic | 0.134 | Male | no info | 1.9900 | 148.6 |
| 6462 | NA | Holyoke MA (1886) | Historic | 0.089 | Male | no info | 6.9600 | 148.6 |
| 7 | NA | Central PA | NA | 0.094 | Male | no info | 4.9200 | 148.8 |
| 77 | NA | Westfield MA | NA | 0.420 | Female | no info | 3.7400 | 148.6 |
| 78 | NA | Westfield MA | NA | 0.386 | Female | no info | 3.7200 | 148.6 |
| 8 | NA | Central PA | NA | 0.435 | Female | no info | 3.5300 | 148.3 |
| 80 | NA | Westfield MA | NA | 0.448 | Female | no info | 0.7800 | 148.1 |
| 81 | NA | Westfield MA | NA | 0.215 | Male | no info | 5.4800 | 148.7 |
| 82303 | NA | Mt Washington MA | L5 | 0.078 | Male | no info | 3.8900 | 148.6 |
| 84 | NA | Westfield MA | NA | 0.089 | Male | no info | 3.6200 | 148.6 |
| 9 | NA | Central PA | NA | 0.085 | Male | no info | 6.2400 | 148.8 |
| 90 | NA | Westfield MA | NA | 0.484 | Female | no info | 5.7700 | 148.6 |
| 91 | NA | Westfield MA | NA | 0.098 | Male | no info | 3.5800 | 148.5 |
| 92 | NA | Westfield MA | NA | 0.095 | Male | no info | 2.0100 | 148.4 |
| 93 | NA | Westfield MA | NA | 0.087 | Male | no info | 5.4500 | 148.6 |
| 99 | NA | Westfield MA | NA | 0.462 | Female | no info | 4.3100 | 148.5 |
| R-4404 | NA | Lynn MA (1878) | Historic | 0.090 | Male | no info | 3.7200 | 148.4 |
| VT1 | NA | Rutland VT L1 | NA | 0.160 | Male | no info | 18.7400 | 148.9 |
| VT10 | NA | Rutland VT L1 | NA | 0.536 | Female | no info | 17.4400 | 148.8 |
| VT100 | NA | Rutland VT L2 | NA | 0.379 | Female | no info | 3.7100 | 148.2 |
| VT101 | NA | Rutland VT L2 | NA | 0.084 | Male | no info | 6.3300 | 148.8 |
| VT102 | NA | Rutland VT L2 | NA | 0.326 | Female | no info | 3.7000 | 148.6 |
| VT103 | NA | Rutland VT L2 | NA | 0.421 | Female | no info | 6.2200 | 148.8 |
| VT104 | NA | Rutland VT L2 | NA | 0.416 | Female | no info | 5.4400 | 148.5 |
| VT105 | NA | Rutland VT L2 | NA | 0.139 | Male | no info | 6.9900 | 148.8 |
| VT107 | NA | Rutland VT L2 | NA | 0.424 | Female | no info | 6.0800 | 148.8 |
| VT109 | NA | Rutland VT L2 | NA | 0.083 | Male | no info | 5.1800 | 148.5 |
| VT110 | NA | Rutland VT L2 | NA | 0.419 | Female | no info | 5.4200 | 148.5 |
| VT111 | NA | Rutland VT L2 | NA | 0.413 | Female | no info | 3.3200 | 148.7 |
| VT12 | NA | Rutland VT L1 | NA | 0.519 | Female | no info | 4.8600 | 148.8 |
| VT16 | NA | Rutland VT L1 | NA | 0.439 | Female | no info | 7.0000 | 148.8 |

**Table S1. Sample metadata**

Metadata for all sequenced timber rattlesnakes (n=111), including field ID (“snake”), color morph, population, sex proxy (WZ_ratio), and inferred sex, SFD status, and sequencing yield (Total_Gbp, Mean_Read_Length_bp).

| snake | Color | population | precise site | W/Z ratio | Inferred sex | lesions | Total Gbp | Mean Read Length (bp) |
|---|---|---|---|---|---|---|---|---|
| VT20 | NA | Rutland VT L1 | NA | 0.124 | Male | no info | 1.5200 | 148.1 |
| VT24 | NA | Rutland VT L1 | NA | 0.086 | Male | no info | 5.7800 | 148.5 |
| VT30 | NA | Rutland VT L1 | NA | 0.099 | Male | no info | 9.2800 | 148.6 |
| VT34 | NA | Rutland VT L1 | NA | 0.083 | Male | no info | 6.5000 | 148.6 |
| VT4 | NA | Rutland VT L1 | NA | 0.087 | Male | no info | 5.8700 | 148.5 |
| VT40 | NA | Rutland VT L1 | NA | 0.500 | Female | no info | 1.6900 | 148.3 |
| VT44 | NA | Rutland VT L1 | NA | 0.420 | Female | no info | 5.9900 | 148.8 |
| VT50 | NA | Rutland VT L1 | NA | 0.087 | Male | no info | 6.5200 | 148.8 |
| VT52 | NA | Rutland VT L1 | NA | 0.083 | Male | no info | 5.1900 | 148.4 |
| VT56 | NA | Rutland VT L1 | NA | 0.091 | Male | no info | 5.8600 | 148.5 |
| VT70 | NA | Rutland VT L1 | NA | 0.660 | Female | no info | 5.4300 | 148.8 |
| VT8 | NA | Rutland VT L1 | NA | 0.429 | Female | no info | 6.7700 | 148.8 |

**Table S2. Population counts & lesion status**

Per-population counts summarizing sample size, color morph totals, and snake fungal disease lesion status (observed / none / missing).

| population | n | Color: black | Color: yellow | fungal lesions observed | no fungal lesions | no data on fungal lesions | host genome coverage >0.05x | inferred female | inferred male |
|---|---|---|---|---|---|---|---|---|---|
| Mt Washington, MA | 40 | 18 | 21 | 6 | 21 | 13 | 38 | 15 | 23 |
| Copake NY | 11 | 7 | 4 | 1 | 8 | 2 | 11 | 4 | 7 |
| Milton MA | 5 | 0 | 0 | 0 | 0 | 5 | 0 | 0 | 0 |
| Rockland County NY | 2 | 0 | 0 | 0 | 0 | 2 | 2 | 1 | 1 |
| Rutland VT | 26 | 0 | 0 | 0 | 0 | 26 | 25 | 14 | 11 |
| Central PA | 10 | 0 | 0 | 0 | 0 | 10 | 10 | 4 | 6 |
| Westfield MA | 13 | 0 | 0 | 0 | 0 | 13 | 11 | 6 | 5 |

### Table S3. Alignment metrics to C. adamanteus

Per-sample mapping summary against a non-conspecific pit viper reference, including raw reads/pairs, primary mapped/Q30 reads, properly paired reads.

| snake | raw reads total | raw pairs | is paired | raw r1 | raw r2 | primary total in bam | primary mapped | primary mapped q30 | primary pp reads | percent input captured | pct mapped vs fastq | pct mapped q30 vs fastq | pct properly paired pair vs fastq |
|---|---|---|---|---|---|---|---|---|---|---|---|---|---|
| VT110 | 36,500,422 | 18,250,211 | 1 | 18,250,211 | 18,250,211 | 30,630,829 | 30,630,829 | 29,256,107 | 30,630,829 | 83.9200 | 83.9200 | 80.1500 | 83.9200 |
| VT111 | 22,350,592 | 11,175,296 | 1 | 11,175,296 | 11,175,296 | 18,329,316 | 18,329,316 | 17,518,839 | 18,329,316 | 82.0100 | 82.0100 | 78.3800 | 82.0100 |
| 12.11 | 12,126,054 | 6,063,027 | 1 | 6,063,027 | 6,063,027 | 8,397,838 | 8,397,838 | 7,782,837 | 8,397,838 | 69.2500 | 69.2500 | 64.1800 | 69.2500 |
| 19192 | 52,363,920 | 26,181,960 | 1 | 26,181,960 | 26,181,960 | 1,733,080 | 1,733,080 | 1,671,020 | 1,733,080 | 3.3100 | 3.3100 | 3.1900 | 3.3100 |
| 12.32 | 28,762,130 | 14,381,065 | 1 | 14,381,065 | 14,381,065 | 22,652,270 | 22,652,270 | 21,384,686 | 22,652,270 | 78.7600 | 78.7600 | 74.3500 | 78.7600 |
| 12.19 | 28,984,816 | 14,492,408 | 1 | 14,492,408 | 14,492,408 | 22,273,984 | 22,273,984 | 20,700,266 | 22,273,984 | 76.8500 | 76.8500 | 71.4200 | 76.8500 |
| 12.34 | 12,923,180 | 6,461,590 | 1 | 6,461,590 | 6,461,590 | 8,743,497 | 8,743,497 | 8,264,683 | 8,743,497 | 67.6600 | 67.6600 | 63.9500 | 67.6600 |
| 11.1 | 4,243,028 | 2,121,514 | 1 | 2,121,514 | 2,121,514 | 2,366,091 | 2,366,091 | 2,260,418 | 2,366,091 | 55.7600 | 55.7600 | 53.2700 | 55.7600 |
| 12.5 | 4,130,382 | 2,065,191 | 1 | 2,065,191 | 2,065,191 | 2,904,906 | 2,904,906 | 2,745,142 | 2,904,906 | 70.3300 | 70.3300 | 66.4600 | 70.3300 |
| 12.51 | 433,638 | 216,819 | 1 | 216,819 | 216,819 | 86,504 | 86,504 | 78,793 | 86,504 | 19.9500 | 19.9500 | 18.1700 | 19.9500 |
| 12.52 | 24,351,056 | 12,175,528 | 1 | 12,175,528 | 12,175,528 | 18,976,998 | 18,976,998 | 17,815,787 | 18,976,998 | 77.9300 | 77.9300 | 73.1600 | 77.9300 |
| 12.39 | 37,417,822 | 18,708,911 | 1 | 18,708,911 | 18,708,911 | 31,116,702 | 31,116,702 | 29,648,366 | 31,116,702 | 83.1600 | 83.1600 | 79.2400 | 83.1600 |
| 11.3 | 32,714,970 | 16,357,485 | 1 | 16,357,485 | 16,357,485 | 26,491,415 | 26,491,415 | 25,055,539 | 26,491,415 | 80.9800 | 80.9800 | 76.5900 | 80.9800 |
| 11.31 | 41,656,298 | 20,828,149 | 1 | 20,828,149 | 20,828,149 | 33,867,107 | 33,867,107 | 32,350,875 | 33,867,107 | 81.3000 | 81.3000 | 77.6600 | 81.3000 |
| 12.71 | 32,422,910 | 16,211,455 | 1 | 16,211,455 | 16,211,455 | 26,888,865 | 26,888,865 | 25,709,463 | 26,888,865 | 82.9300 | 82.9300 | 79.2900 | 82.9300 |
| 11.32 | 35,281,448 | 17,640,724 | 1 | 17,640,724 | 17,640,724 | 28,650,376 | 28,650,376 | 27,148,891 | 28,650,376 | 81.2100 | 81.2100 | 76.9500 | 81.2100 |
| 12.73 | 68,706,304 | 34,353,152 | 1 | 34,353,152 | 34,353,152 | 48,972,116 | 48,972,116 | 46,016,055 | 48,972,116 | 71.2800 | 71.2800 | 66.9800 | 71.2800 |
| 12.74 | 7,174,116 | 3,587,058 | 1 | 3,587,058 | 3,587,058 | 681,081 | 681,081 | 627,122 | 681,081 | 9.4900 | 9.4900 | 8.7400 | 9.4900 |
| 11.35 | 35,433,382 | 17,716,691 | 1 | 17,716,691 | 17,716,691 | 25,918,802 | 25,918,802 | 23,723,140 | 25,918,802 | 73.1500 | 73.1500 | 66.9500 | 73.1500 |
| 12.75 | 11,234,822 | 5,617,411 | 1 | 5,617,411 | 5,617,411 | 6,707,923 | 6,707,923 | 6,393,163 | 6,707,923 | 59.7100 | 59.7100 | 56.9000 | 59.7100 |
| 11.36 | 30,008,552 | 15,004,276 | 1 | 15,004,276 | 15,004,276 | 24,566,280 | 24,566,280 | 23,382,189 | 24,566,280 | 81.8600 | 81.8600 | 77.9200 | 81.8600 |
| 11.38 | 34,509,168 | 17,254,584 | 1 | 17,254,584 | 17,254,584 | 28,736,835 | 28,736,835 | 27,421,265 | 28,736,835 | 83.2700 | 83.2700 | 79.4600 | 83.2700 |
| 1 | 36,540,180 | 18,270,090 | 1 | 18,270,090 | 18,270,090 | 29,789,678 | 29,789,678 | 28,127,972 | 29,789,678 | 81.5300 | 81.5300 | 76.9800 | 81.5300 |
| 11.52 | 39,696,028 | 19,848,014 | 1 | 19,848,014 | 19,848,014 | 33,487,048 | 33,487,048 | 32,029,714 | 33,487,048 | 84.3600 | 84.3600 | 80.6900 | 84.3600 |
| 2 | 22,864,266 | 11,432,133 | 1 | 11,432,133 | 11,432,133 | 18,950,136 | 18,950,136 | 18,052,403 | 18,950,136 | 82.8800 | 82.8800 | 78.9500 | 82.8800 |
| 11.53 | 28,070,230 | 14,035,115 | 1 | 14,035,115 | 14,035,115 | 23,350,005 | 23,350,005 | 22,281,170 | 23,350,005 | 83.1800 | 83.1800 | 79.3800 | 83.1800 |
| VT10 | 117,181,354 | 58,590,677 | 1 | 58,590,677 | 58,590,677 | 90,980,221 | 90,980,221 | 85,138,353 | 90,980,221 | 77.6400 | 77.6400 | 72.6600 | 77.6400 |
| 3 | 25,818,460 | 12,909,230 | 1 | 12,909,230 | 12,909,230 | 21,272,849 | 21,272,849 | 20,205,597 | 21,272,849 | 82.3900 | 82.3900 | 78.2600 | 82.3900 |
| 11.54 | 40,030,232 | 20,015,116 | 1 | 20,015,116 | 20,015,116 | 31,731,195 | 31,731,195 | 29,971,496 | 31,731,195 | 79.2700 | 79.2700 | 74.8700 | 79.2700 |
| 10.16 | 28,682,340 | 14,341,170 | 1 | 14,341,170 | 14,341,170 | 22,844,070 | 22,844,070 | 21,534,188 | 22,844,070 | 79.6500 | 79.6500 | 75.0800 | 79.6500 |
| VT12 | 32,678,204 | 16,339,102 | 1 | 16,339,102 | 16,339,102 | 22,751,051 | 22,751,051 | 21,096,340 | 22,751,051 | 69.6200 | 69.6200 | 64.5600 | 69.6200 |
| 5 | 30,204,526 | 15,102,263 | 1 | 15,102,263 | 15,102,263 | 25,584,127 | 25,584,127 | 24,506,659 | 25,584,127 | 84.7000 | 84.7000 | 81.1400 | 84.7000 |
| 10.17 | 14,711,078 | 7,355,539 | 1 | 7,355,539 | 7,355,539 | 10,595,463 | 10,595,463 | 9,903,351 | 10,595,463 | 72.0200 | 72.0200 | 67.3200 | 72.0200 |
| 80 | 5,293,652 | 2,646,826 | 1 | 2,646,826 | 2,646,826 | 4,257,884 | 4,257,884 | 4,052,997 | 4,257,884 | 80.4300 | 80.4300 | 76.5600 | 80.4300 |
| 6 | 26,207,930 | 13,103,527 | 1 | 13,104,403 | 13,103,527 | 22,627,233 | 22,627,233 | 21,699,043 | 22,627,233 | 86.3400 | 86.3400 | 82.8000 | 86.3400 |
| 09.06SFD | 33,061,458 | 16,530,729 | 1 | 16,530,729 | 16,530,729 | 27,876,010 | 27,876,010 | 26,669,907 | 27,876,010 | 84.3200 | 84.3200 | 80.6700 | 84.3200 |
| 11.57 | 42,519,270 | 21,259,635 | 1 | 21,259,635 | 21,259,635 | 35,503,268 | 35,503,268 | 33,911,537 | 35,503,268 | 83.5000 | 83.5000 | 79.7600 | 83.5000 |
| 81 | 36,880,144 | 18,440,072 | 1 | 18,440,072 | 18,440,072 | 2,842,841 | 2,842,841 | 2,725,350 | 2,842,841 | 7.7100 | 7.7100 | 7.3900 | 7.7100 |
| 7 | 33,041,008 | 16,520,504 | 1 | 16,520,504 | 16,520,504 | 27,329,561 | 27,329,561 | 26,059,385 | 27,329,561 | 82.7100 | 82.7100 | 78.8700 | 82.7100 |
| 11.58 | 15,751,880 | 7,875,940 | 1 | 7,875,940 | 7,875,940 | 8,096,283 | 8,096,283 | 7,500,506 | 8,096,283 | 51.4000 | 51.4000 | 47.6200 | 51.4000 |
| 8 | 23,797,712 | 11,898,856 | 1 | 11,898,856 | 11,898,856 | 19,539,998 | 19,539,998 | 18,613,409 | 19,539,998 | 82.1100 | 82.1100 | 78.2200 | 82.1100 |
| VT16 | 47,050,854 | 23,525,427 | 1 | 23,525,427 | 23,525,427 | 38,880,230 | 38,880,230 | 37,026,081 | 38,880,230 | 82.6300 | 82.6300 | 78.6900 | 82.6300 |
| 9 | 41,928,712 | 20,964,356 | 1 | 20,964,356 | 20,964,356 | 35,452,493 | 35,452,493 | 33,932,728 | 35,452,493 | 84.5500 | 84.5500 | 80.9300 | 84.5500 |
| 11.73 | 9,709,736 | 4,854,868 | 1 | 4,854,868 | 4,854,868 | 4,504,356 | 4,504,356 | 4,163,903 | 4,504,356 | 46.3900 | 46.3900 | 42.8800 | 46.3900 |
| 10.34 | 12,980,004 | 6,490,002 | 1 | 6,490,002 | 6,490,002 | 10,454,726 | 10,454,726 | 10,008,614 | 10,454,726 | 80.5400 | 80.5400 | 77.1100 | 80.5400 |
| VT30 | 62,425,848 | 31,212,924 | 1 | 31,212,924 | 31,212,924 | 50,795,372 | 50,795,372 | 48,228,637 | 50,795,372 | 81.3700 | 81.3700 | 77.2600 | 81.3700 |
| 84 | 24,338,190 | 12,169,095 | 1 | 12,169,095 | 12,169,095 | 20,095,544 | 20,095,544 | 19,203,151 | 20,095,544 | 82.5700 | 82.5700 | 78.9000 | 82.5700 |
| 11.74 | 25,149,744 | 12,574,872 | 1 | 12,574,872 | 12,574,872 | 16,441,230 | 16,441,230 | 15,054,389 | 16,441,230 | 65.3700 | 65.3700 | 59.8600 | 65.3700 |
| VT34 | 43,729,060 | 21,864,530 | 1 | 21,864,530 | 21,864,530 | 36,706,723 | 36,706,723 | 35,136,274 | 36,706,723 | 83.9400 | 83.9400 | 80.3500 | 83.9400 |
| 6439 | 13,362,788 | 6,681,394 | 1 | 6,681,394 | 6,681,394 | 590,509 | 590,509 | 571,252 | 590,509 | 4.4200 | 4.4200 | 4.2700 | 4.4200 |
| VT50 | 43,798,602 | 21,899,301 | 1 | 21,899,301 | 21,899,301 | 36,970,364 | 36,970,364 | 35,353,224 | 36,970,364 | 84.4100 | 84.4100 | 80.7200 | 84.4100 |
| VT52 | 34,943,248 | 17,471,624 | 1 | 17,471,624 | 17,471,624 | 29,064,160 | 29,064,160 | 27,851,942 | 29,064,160 | 83.1800 | 83.1800 | 79.7100 | 83.1800 |
| VT56 | 39,478,594 | 19,739,297 | 1 | 19,739,297 | 19,739,297 | 32,732,854 | 32,732,854 | 31,175,421 | 32,732,854 | 82.9100 | 82.9100 | 78.9700 | 82.9100 |
| VT70 | 36,501,706 | 18,250,853 | 1 | 18,250,853 | 18,250,853 | 25,778,841 | 25,778,841 | 23,538,193 | 25,778,841 | 70.6200 | 70.6200 | 64.4900 | 70.6200 |
| 82303 | 26,182,190 | 13,090,802 | 1 | 13,091,388 | 13,090,802 | 1,523,674 | 1,523,674 | 1,456,685 | 1,523,674 | 5.8200 | 5.8200 | 5.5600 | 5.8200 |
| 186035 | 1,575,600 | 787,800 | 1 | 787,800 | 787,800 | 1,097,698 | 1,097,698 | 1,042,465 | 1,097,698 | 69.6700 | 69.6700 | 66.1600 | 69.6700 |
| VT100 | 25,033,995 | 12,516,633 | 1 | 12,516,633 | 12,517,362 | 15,601,923 | 15,601,923 | 14,886,553 | 15,601,923 | 62.3200 | 62.3200 | 59.4700 | 62.3200 |
| 186036 | 1,549,182 | 774,591 | 1 | 774,591 | 774,591 | 893,680 | 893,680 | 842,944 | 893,680 | 57.6900 | 57.6900 | 54.4100 | 57.6900 |
| VT101 | 42,517,450 | 21,258,725 | 1 | 21,258,725 | 21,258,725 | 35,696,718 | 35,696,718 | 34,115,674 | 35,696,718 | 83.9600 | 83.9600 | 80.2400 | 83.9600 |
| 186037 | 1,180,434 | 590,217 | 1 | 590,217 | 590,217 | 654,210 | 654,210 | 617,623 | 654,210 | 55.4200 | 55.4200 | 52.3200 | 55.4200 |
| VT102 | 24,905,246 | 12,451,950 | 1 | 12,451,950 | 12,453,296 | 23,256,621 | 23,256,621 | 22,891,171 | 23,256,621 | 93.3800 | 93.3800 | 91.9100 | 93.3900 |
| VT103 | 41,827,002 | 20,913,501 | 1 | 20,913,501 | 20,913,501 | 35,021,238 | 35,021,238 | 33,417,011 | 35,021,238 | 83.7300 | 83.7300 | 79.8900 | 83.7300 |
| VT104 | 36,661,266 | 18,330,633 | 1 | 18,330,633 | 18,330,633 | 30,715,910 | 30,715,910 | 29,345,366 | 30,715,910 | 83.7800 | 83.7800 | 80.0400 | 83.7800 |
| VT105 | 46,991,776 | 23,495,888 | 1 | 23,495,888 | 23,495,888 | 2,499,139 | 2,499,139 | 2,304,248 | 2,499,139 | 5.3200 | 5.3200 | 4.9000 | 5.3200 |
| VT107 | 40,873,946 | 20,436,973 | 1 | 20,436,973 | 20,436,973 | 34,303,630 | 34,303,630 | 32,742,789 | 34,303,630 | 83.9300 | 83.9300 | 80.1100 | 83.9300 |
| VT109 | 34,852,314 | 17,426,157 | 1 | 17,426,157 | 17,426,157 | 29,585,020 | 29,585,020 | 28,344,018 | 29,585,020 | 84.8900 | 84.8900 | 81.3300 | 84.8900 |
| 12.21 | 29,337,764 | 14,668,882 | 1 | 14,668,882 | 14,668,882 | 24,337,276 | 24,337,276 | 23,243,704 | 24,337,276 | 82.9600 | 82.9600 | 79.2300 | 82.9600 |
| 12.23 | 34,959,560 | 17,479,780 | 1 | 17,479,780 | 17,479,780 | 26,560,938 | 26,560,938 | 24,754,751 | 26,560,938 | 75.9800 | 75.9800 | 70.8100 | 75.9800 |
| 12.25 | 29,488,730 | 14,744,365 | 1 | 14,744,365 | 14,744,365 | 24,164,011 | 24,164,011 | 22,945,821 | 24,164,011 | 81.9400 | 81.9400 | 77.8100 | 81.9400 |
| 12.26 | 30,395,994 | 15,197,997 | 1 | 15,197,997 | 15,197,997 | 25,248,939 | 25,248,939 | 24,041,960 | 25,248,939 | 83.0700 | 83.0700 | 79.1000 | 83.0700 |
| 10 | 20,242,036 | 10,121,018 | 1 | 10,121,018 | 10,121,018 | 15,611,161 | 15,611,161 | 14,662,492 | 15,611,161 | 77.1200 | 77.1200 | 72.4400 | 77.1200 |
| 12.4 | 31,702,668 | 15,851,334 | 1 | 15,851,334 | 15,851,334 | 24,838,049 | 24,838,049 | 23,430,848 | 24,838,049 | 78.3500 | 78.3500 | 73.9100 | 78.3500 |
| 11 | 3,945,284 | 1,972,642 | 1 | 1,972,642 | 1,972,642 | 2,930,335 | 2,930,335 | 2,790,973 | 2,930,335 | 74.2700 | 74.2700 | 70.7400 | 74.2700 |
| R-4404 | 25,071,376 | 12,535,398 | 1 | 12,535,398 | 12,535,978 | 1,108,594 | 1,108,594 | 1,074,312 | 1,108,594 | 4.4200 | 4.4200 | 4.2900 | 4.4200 |
| 12.43 | 33,157,842 | 16,578,921 | 1 | 16,578,921 | 16,578,921 | 27,749,680 | 27,749,680 | 26,500,769 | 27,749,680 | 83.6900 | 83.6900 | 79.9200 | 83.6900 |
| 11.04 | 25,425,380 | 12,712,690 | 1 | 12,712,690 | 12,712,690 | 20,900,547 | 20,900,547 | 19,869,687 | 20,900,547 | 82.2000 | 82.2000 | 78.1500 | 82.2000 |
| 11.07 | 30,088,014 | 15,044,007 | 1 | 15,044,007 | 15,044,007 | 25,359,607 | 25,359,607 | 24,300,317 | 25,359,607 | 84.2800 | 84.2800 | 80.7600 | 84.2800 |
| 12.61 | 31,947,908 | 15,973,954 | 1 | 15,973,954 | 15,973,954 | 25,437,449 | 25,437,449 | 24,031,456 | 25,437,449 | 79.6200 | 79.6200 | 75.2200 | 79.6200 |
| 12.48 | 19,862,948 | 9,931,474 | 1 | 9,931,474 | 9,931,474 | 16,347,961 | 16,347,961 | 15,621,688 | 16,347,961 | 82.3000 | 82.3000 | 78.6500 | 82.3000 |
| 11.4 | 40,722,168 | 20,361,084 | 1 | 20,361,084 | 20,361,084 | 33,189,176 | 33,189,176 | 31,383,983 | 33,189,176 | 81.5000 | 81.5000 | 77.0700 | 81.5000 |
| 162019 | 33,423,630 | 16,711,815 | 1 | 16,711,815 | 16,711,815 | 787,330 | 787,330 | 758,939 | 787,330 | 2.3600 | 2.3600 | 2.2700 | 2.3600 |
| 12.49 | 11,465,026 | 5,732,513 | 1 | 5,732,513 | 5,732,513 | 8,677,404 | 8,677,404 | 8,223,100 | 8,677,404 | 75.6900 | 75.6900 | 71.7200 | 75.6900 |
| 11.24 | 24,623,176 | 12,311,588 | 1 | 12,311,588 | 12,311,588 | 20,386,724 | 20,386,724 | 19,428,791 | 20,386,724 | 82.7900 | 82.7900 | 78.9000 | 82.7900 |
| 12.64 | 30,113,684 | 15,056,842 | 1 | 15,056,842 | 15,056,842 | 22,443,494 | 22,443,494 | 20,941,214 | 22,443,494 | 74.5300 | 74.5300 | 69.5400 | 74.5300 |
| 27544 | 35,890,692 | 17,945,346 | 1 | 17,945,346 | 17,945,346 | 1,141,006 | 1,141,006 | 1,102,183 | 1,141,006 | 3.1800 | 3.1800 | 3.0700 | 3.1800 |
| 11.28 | 37,510,038 | 18,755,019 | 1 | 18,755,019 | 18,755,019 | 31,324,653 | 31,324,653 | 29,918,353 | 31,324,653 | 83.5100 | 83.5100 | 79.7600 | 83.5100 |
| 11.41 | 40,710,252 | 20,355,126 | 1 | 20,355,126 | 20,355,126 | 34,038,334 | 34,038,334 | 32,494,298 | 34,038,334 | 83.6100 | 83.6100 | 79.8200 | 83.6100 |
| 11.29 | 29,259,462 | 14,629,731 | 1 | 14,629,731 | 14,629,731 | 23,776,636 | 23,776,636 | 22,505,866 | 23,776,636 | 81.2600 | 81.2600 | 76.9200 | 81.2600 |
| VT1 | 125,878,582 | 62,939,291 | 1 | 62,939,291 | 62,939,291 | 79,559,459 | 79,559,459 | 73,442,603 | 79,559,459 | 63.2000 | 63.2000 | 58.3400 | 63.2000 |
| 11.43 | 27,193,348 | 13,596,674 | 1 | 13,596,674 | 13,596,674 | 21,687,737 | 21,687,737 | 20,447,276 | 21,687,737 | 79.7500 | 79.7500 | 75.1900 | 79.7500 |
| 11.44 | 39,028,780 | 19,514,390 | 1 | 19,514,390 | 19,514,390 | 32,271,344 | 32,271,344 | 30,689,779 | 32,271,344 | 82.6900 | 82.6900 | 78.6300 | 82.6900 |
| 11.45 | 35,240,178 | 17,620,089 | 1 | 17,620,089 | 17,620,089 | 28,157,635 | 28,157,635 | 26,521,750 | 28,157,635 | 79.9000 | 79.9000 | 75.2600 | 79.9000 |
| VT4 | 39,522,840 | 19,761,420 | 1 | 19,761,420 | 19,761,420 | 32,972,997 | 32,972,997 | 31,479,687 | 32,972,997 | 83.4300 | 83.4300 | 79.6500 | 83.4300 |
| 11.46 | 43,277,470 | 21,638,735 | 1 | 21,638,735 | 21,638,735 | 36,412,977 | 36,412,977 | 34,806,913 | 36,412,977 | 84.1400 | 84.1400 | 80.4300 | 84.1400 |
| 10.21 | 15,249,038 | 7,624,519 | 1 | 7,624,519 | 7,624,519 | 10,157,114 | 10,157,114 | 9,596,129 | 10,157,114 | 66.6100 | 66.6100 | 62.9300 | 66.6100 |
| VT8 | 45,507,146 | 22,753,573 | 1 | 22,753,573 | 22,753,573 | 37,966,461 | 37,966,461 | 36,177,938 | 37,966,461 | 83.4300 | 83.4300 | 79.5000 | 83.4300 |
| VT20 | 10,284,758 | 5,142,379 | 1 | 5,142,379 | 5,142,379 | 6,666,278 | 6,666,278 | 6,224,859 | 6,666,278 | 64.8200 | 64.8200 | 60.5300 | 64.8200 |
| 11.64 | 16,018,750 | 8,009,375 | 1 | 8,009,375 | 8,009,375 | 11,538,598 | 11,538,598 | 10,756,237 | 11,538,598 | 72.0300 | 72.0300 | 67.1500 | 72.0300 |

### Table S3. Alignment metrics to C. adamanteus

Per-sample mapping summary against a non-conspecific pit viper reference, including raw reads/pairs, primary mapped/Q30 reads, properly paired reads.

| snake | raw reads total | raw pairs | is paired | raw r1 | raw r2 | primary total in bam | primary mapped | primary mapped q30 | primary pp reads | percent input captured | pct mapped vs fastq | pct mapped q30 vs fastq | pct properly paired pair vs fastq |
|---|---|---|---|---|---|---|---|---|---|---|---|---|---|
| 10.4 | 38,940,546 | 19,470,273 | 1 | 19,470,273 | 19,470,273 | 32,715,554 | 32,715,554 | 31,331,376 | 32,715,554 | 84.0100 | 84.0100 | 80.4600 | 84.0100 |
| 90 | 38,805,910 | 19,402,955 | 1 | 19,402,955 | 19,402,955 | 31,723,484 | 31,723,484 | 30,047,631 | 31,723,484 | 81.7500 | 81.7500 | 77.4300 | 81.7500 |
| 77 | 25,153,460 | 12,576,171 | 1 | 12,576,171 | 12,577,289 | 22,390,316 | 22,390,316 | 21,708,531 | 22,390,316 | 89.0100 | 89.0100 | 86.3000 | 89.0200 |
| VT24 | 38,913,786 | 19,456,893 | 1 | 19,456,893 | 19,456,893 | 32,475,357 | 32,475,357 | 31,002,835 | 32,475,357 | 83.4500 | 83.4500 | 79.6700 | 83.4500 |
| 91 | 24,101,706 | 12,050,853 | 1 | 12,050,853 | 12,050,853 | 19,305,969 | 19,305,969 | 18,303,901 | 19,305,969 | 80.1000 | 80.1000 | 75.9400 | 80.1000 |
| 78 | 25,040,951 | 12,520,011 | 1 | 12,520,011 | 12,520,940 | 23,127,248 | 23,127,248 | 22,668,521 | 23,127,248 | 92.3600 | 92.3600 | 90.5300 | 92.3600 |
| 11.68 | 126,330,410 | 63,165,205 | 1 | 63,165,205 | 63,165,205 | 105,701,097 | 105,701,097 | 100,777,682 | 105,701,097 | 83.6700 | 83.6700 | 79.7700 | 83.6700 |
| 92 | 13,546,924 | 6,773,462 | 1 | 6,773,462 | 6,773,462 | 10,977,022 | 10,977,022 | 10,452,784 | 10,977,022 | 81.0300 | 81.0300 | 77.1600 | 81.0300 |
| 93 | 36,702,800 | 18,350,215 | 1 | 18,350,215 | 18,352,585 | 30,850,193 | 30,850,193 | 29,439,958 | 30,850,193 | 84.0500 | 84.0500 | 80.2100 | 84.0600 |
| VT40 | 11,413,826 | 5,706,913 | 1 | 5,706,913 | 5,706,913 | 8,699,750 | 8,699,750 | 8,222,626 | 8,699,750 | 76.2200 | 76.2200 | 72.0400 | 76.2200 |
| VT44 | 40,257,834 | 20,128,917 | 1 | 20,128,917 | 20,128,917 | 33,394,898 | 33,394,898 | 31,883,684 | 33,394,898 | 82.9500 | 82.9500 | 79.2000 | 82.9500 |
| 99 | 29,045,652 | 14,522,826 | 1 | 14,522,826 | 14,522,826 | 23,809,829 | 23,809,829 | 22,607,127 | 23,809,829 | 81.9700 | 81.9700 | 77.8300 | 81.9700 |
| 6462 | 46,817,378 | 23,408,689 | 1 | 23,408,689 | 23,408,689 | 1,670,387 | 1,670,387 | 1,612,362 | 1,670,387 | 3.5700 | 3.5700 | 3.4400 | 3.5700 |

**Table S4. Alignment metrics to C. horridus**

Per-sample host alignment summary to the timber rattlesnake reference, mirroring S3 fields to enable direct comparison of mapping rates and Q30 retention by sample.

| snake | raw reads total | raw pairs | is paired | raw r1 | raw r2 | primary total in bam | primary mapped | primary mapped q30 | primary pp reads | percent input captured | pct mapped vs fastq | pct mapped q30 vs fastq | pct properly paired pair vs fastq |
|---|---|---|---|---|---|---|---|---|---|---|---|---|---|
| VT110 | 36,500,422 | 18,250,211 | 1 | 18,250,211 | 18,250,211 | 32,762,477 | 32,762,477 | 31,904,260 | 32,762,477 | 89.7600 | 89.7600 | 87.4100 | 89.7600 |
| VT111 | 22,350,592 | 11,175,296 | 1 | 11,175,296 | 11,175,296 | 19,395,058 | 19,395,058 | 18,926,227 | 19,395,058 | 86.7800 | 86.7800 | 84.6800 | 86.7800 |
| 12.11 | 12,126,054 | 6,063,027 | 1 | 6,063,027 | 6,063,027 | 8,824,715 | 8,824,715 | 8,589,966 | 8,824,715 | 72.7700 | 72.7700 | 70.8400 | 72.7700 |
| 19192 | 52,363,920 | 26,181,960 | 1 | 26,181,960 | 26,181,960 | 2,171,162 | 2,171,162 | 2,111,998 | 2,171,162 | 4.1500 | 4.1500 | 4.0300 | 4.1500 |
| 12.32 | 28,762,130 | 14,381,065 | 1 | 14,381,065 | 14,381,065 | 24,305,732 | 24,305,732 | 23,535,491 | 24,305,732 | 84.5100 | 84.5100 | 81.8300 | 84.5100 |
| 12.19 | 28,984,816 | 14,492,408 | 1 | 14,492,408 | 14,492,408 | 24,405,617 | 24,405,617 | 23,450,519 | 24,405,617 | 84.2000 | 84.2000 | 80.9100 | 84.2000 |
| 12.34 | 12,923,180 | 6,461,590 | 1 | 6,461,590 | 6,461,590 | 9,339,963 | 9,339,963 | 9,062,845 | 9,339,963 | 72.2700 | 72.2700 | 70.1300 | 72.2700 |
| 11.1 | 4,243,028 | 2,121,514 | 1 | 2,121,514 | 2,121,514 | 2,486,069 | 2,486,069 | 2,424,491 | 2,486,069 | 58.5900 | 58.5900 | 57.1400 | 58.5900 |
| 12.5 | 4,130,382 | 2,065,191 | 1 | 2,065,191 | 2,065,191 | 3,085,285 | 3,085,285 | 2,997,712 | 3,085,285 | 74.7000 | 74.7000 | 72.5800 | 74.7000 |
| 12.51 | 433,638 | 216,819 | 1 | 216,819 | 216,819 | 89,456 | 89,456 | 82,491 | 89,456 | 20.6300 | 20.6300 | 19.0200 | 20.6300 |
| 12.52 | 24,351,056 | 12,175,528 | 1 | 12,175,528 | 12,175,528 | 20,494,758 | 20,494,758 | 19,819,110 | 20,494,758 | 84.1600 | 84.1600 | 81.3900 | 84.1600 |
| 12.39 | 37,417,822 | 18,708,911 | 1 | 18,708,911 | 18,708,911 | 33,301,216 | 33,301,216 | 32,412,451 | 33,301,216 | 89.0000 | 89.0000 | 86.6200 | 89.0000 |
| 11.3 | 32,714,970 | 16,357,485 | 1 | 16,357,485 | 16,357,485 | 28,557,231 | 28,557,231 | 27,692,476 | 28,557,231 | 87.2900 | 87.2900 | 84.6500 | 87.2900 |
| 11.31 | 41,656,298 | 20,828,149 | 1 | 20,828,149 | 20,828,149 | 35,958,603 | 35,958,603 | 35,025,186 | 35,958,603 | 86.3200 | 86.3200 | 84.0800 | 86.3200 |
| 12.71 | 32,422,910 | 16,211,455 | 1 | 16,211,455 | 16,211,455 | 28,744,587 | 28,744,587 | 28,031,778 | 28,744,587 | 88.6600 | 88.6600 | 86.4600 | 88.6600 |
| 11.32 | 35,281,448 | 17,640,724 | 1 | 17,640,724 | 17,640,724 | 30,726,572 | 30,726,572 | 29,826,020 | 30,726,572 | 87.0900 | 87.0900 | 84.5400 | 87.0900 |
| 12.73 | 68,706,304 | 34,353,152 | 1 | 34,353,152 | 34,353,152 | 52,266,490 | 52,266,490 | 50,464,608 | 52,266,490 | 76.0700 | 76.0700 | 73.4500 | 76.0700 |
| 12.74 | 7,174,116 | 3,587,058 | 1 | 3,587,058 | 3,587,058 | 700,076 | 700,076 | 653,931 | 700,076 | 9.7600 | 9.7600 | 9.1200 | 9.7600 |
| 11.35 | 35,433,382 | 17,716,691 | 1 | 17,716,691 | 17,716,691 | 28,736,953 | 28,736,953 | 27,387,557 | 28,736,953 | 81.1000 | 81.1000 | 77.2900 | 81.1000 |
| 12.75 | 11,234,822 | 5,617,411 | 1 | 5,617,411 | 5,617,411 | 7,069,033 | 7,069,033 | 6,891,177 | 7,069,033 | 62.9200 | 62.9200 | 61.3400 | 62.9200 |
| 11.36 | 30,008,552 | 15,004,276 | 1 | 15,004,276 | 15,004,276 | 26,226,093 | 26,226,093 | 25,513,706 | 26,226,093 | 87.4000 | 87.4000 | 85.0200 | 87.4000 |
| 11.38 | 34,509,168 | 17,254,584 | 1 | 17,254,584 | 17,254,584 | 30,800,046 | 30,800,046 | 29,975,869 | 30,800,046 | 89.2500 | 89.2500 | 86.8600 | 89.2500 |
| 1 | 36,540,180 | 18,270,090 | 1 | 18,270,090 | 18,270,090 | 32,125,480 | 32,125,480 | 31,127,766 | 32,125,480 | 87.9200 | 87.9200 | 85.1900 | 87.9200 |
| 11.52 | 39,696,028 | 19,848,014 | 1 | 19,848,014 | 19,848,014 | 35,712,518 | 35,712,518 | 34,833,717 | 35,712,518 | 89.9600 | 89.9600 | 87.7500 | 89.9600 |
| 2 | 22,864,266 | 11,432,133 | 1 | 11,432,133 | 11,432,133 | 20,203,522 | 20,203,522 | 19,691,146 | 20,203,522 | 88.3600 | 88.3600 | 86.1200 | 88.3600 |
| 11.53 | 28,070,230 | 14,035,115 | 1 | 14,035,115 | 14,035,115 | 24,894,712 | 24,894,712 | 24,253,418 | 24,894,712 | 88.6900 | 88.6900 | 86.4000 | 88.6900 |
| VT10 | 117,181,354 | 58,590,677 | 1 | 58,590,677 | 58,590,677 | 95,015,002 | 95,015,002 | 92,488,688 | 95,015,002 | 81.0800 | 81.0800 | 78.9300 | 81.0800 |
| 3 | 25,818,460 | 12,909,230 | 1 | 12,909,230 | 12,909,230 | 22,731,283 | 22,731,283 | 22,115,868 | 22,731,283 | 88.0400 | 88.0400 | 85.6600 | 88.0400 |
| 11.54 | 40,030,232 | 20,015,116 | 1 | 20,015,116 | 20,015,116 | 34,382,721 | 34,382,721 | 33,302,769 | 34,382,721 | 85.8900 | 85.8900 | 83.1900 | 85.8900 |
| 10.16 | 28,682,340 | 14,341,170 | 1 | 14,341,170 | 14,341,170 | 24,608,546 | 24,608,546 | 23,841,787 | 24,608,546 | 85.8000 | 85.8000 | 83.1200 | 85.8000 |
| VT12 | 32,678,204 | 16,339,102 | 1 | 16,339,102 | 16,339,102 | 24,250,823 | 24,250,823 | 23,459,472 | 24,250,823 | 74.2100 | 74.2100 | 71.7900 | 74.2100 |
| 5 | 30,204,526 | 15,102,263 | 1 | 15,102,263 | 15,102,263 | 27,245,692 | 27,245,692 | 26,612,720 | 27,245,692 | 90.2000 | 90.2000 | 88.1100 | 90.2000 |
| 10.17 | 14,711,078 | 7,355,539 | 1 | 7,355,539 | 7,355,539 | 11,473,481 | 11,473,481 | 11,060,169 | 11,473,481 | 77.9900 | 77.9900 | 75.1800 | 77.9900 |
| 80 | 5,293,652 | 2,646,826 | 1 | 2,646,826 | 2,646,826 | 4,527,451 | 4,527,451 | 4,410,026 | 4,527,451 | 85.5300 | 85.5300 | 83.3100 | 85.5300 |
| 6 | 26,207,930 | 13,103,527 | 1 | 13,104,403 | 13,103,527 | 24,410,334 | 24,410,334 | 23,912,624 | 24,410,334 | 93.1400 | 93.1400 | 91.2400 | 93.1400 |
| 09.06SFD | 33,061,458 | 16,530,729 | 1 | 16,530,729 | 16,530,729 | 29,712,824 | 29,712,824 | 28,988,335 | 29,712,824 | 89.8700 | 89.8700 | 87.6800 | 89.8700 |
| 11.57 | 42,519,270 | 21,259,635 | 1 | 21,259,635 | 21,259,635 | 38,023,327 | 38,023,327 | 37,030,549 | 38,023,327 | 89.4300 | 89.4300 | 87.0900 | 89.4300 |
| 81 | 36,880,144 | 18,440,072 | 1 | 18,440,072 | 18,440,072 | 2,890,180 | 2,890,180 | 2,787,806 | 2,890,180 | 7.8400 | 7.8400 | 7.5600 | 7.8400 |
| 7 | 33,041,008 | 16,520,504 | 1 | 16,520,504 | 16,520,504 | 29,140,821 | 29,140,821 | 28,408,375 | 29,140,821 | 88.2000 | 88.2000 | 85.9800 | 88.2000 |
| 11.58 | 15,751,880 | 7,875,940 | 1 | 7,875,940 | 7,875,940 | 8,771,620 | 8,771,620 | 8,431,662 | 8,771,620 | 55.6900 | 55.6900 | 53.5300 | 55.6900 |
| 8 | 23,797,712 | 11,898,856 | 1 | 11,898,856 | 11,898,856 | 20,824,478 | 20,824,478 | 20,293,270 | 20,824,478 | 87.5100 | 87.5100 | 85.2700 | 87.5100 |
| VT16 | 47,050,854 | 23,525,427 | 1 | 23,525,427 | 23,525,427 | 41,703,798 | 41,703,798 | 40,546,962 | 41,703,798 | 88.6400 | 88.6400 | 86.1800 | 88.6400 |
| 9 | 41,928,712 | 20,964,356 | 1 | 20,964,356 | 20,964,356 | 37,881,141 | 37,881,141 | 36,976,358 | 37,881,141 | 90.3500 | 90.3500 | 88.1900 | 90.3500 |
| 11.73 | 9,709,736 | 4,854,868 | 1 | 4,854,868 | 4,854,868 | 4,849,040 | 4,849,040 | 4,631,028 | 4,849,040 | 49.9400 | 49.9400 | 47.6900 | 49.9400 |
| 10.34 | 12,980,004 | 6,490,002 | 1 | 6,490,002 | 6,490,002 | 11,047,932 | 11,047,932 | 10,793,812 | 11,047,932 | 85.1200 | 85.1200 | 83.1600 | 85.1200 |
| VT30 | 62,425,848 | 31,212,924 | 1 | 31,212,924 | 31,212,924 | 53,821,147 | 53,821,147 | 52,457,777 | 53,821,147 | 86.2200 | 86.2200 | 84.0300 | 86.2200 |
| 84 | 24,338,190 | 12,169,095 | 1 | 12,169,095 | 12,169,095 | 21,487,217 | 21,487,217 | 20,939,161 | 21,487,217 | 88.2900 | 88.2900 | 86.0300 | 88.2900 |
| 11.74 | 25,149,744 | 12,574,872 | 1 | 12,574,872 | 12,574,872 | 17,950,947 | 17,950,947 | 17,097,801 | 17,950,947 | 71.3800 | 71.3800 | 67.9800 | 71.3800 |
| VT34 | 43,729,060 | 21,864,530 | 1 | 21,864,530 | 21,864,530 | 39,231,421 | 39,231,421 | 38,246,973 | 39,231,421 | 89.7100 | 89.7100 | 87.4600 | 89.7100 |
| 6439 | 13,362,788 | 6,681,394 | 1 | 6,681,394 | 6,681,394 | 707,956 | 707,956 | 689,758 | 707,956 | 5.3000 | 5.3000 | 5.1600 | 5.3000 |
| VT50 | 43,798,602 | 21,899,301 | 1 | 21,899,301 | 21,899,301 | 39,216,693 | 39,216,693 | 38,284,193 | 39,216,693 | 89.5400 | 89.5400 | 87.4100 | 89.5400 |
| VT52 | 34,943,248 | 17,471,624 | 1 | 17,471,624 | 17,471,624 | 30,916,709 | 30,916,709 | 30,170,589 | 30,916,709 | 88.4800 | 88.4800 | 86.3400 | 88.4800 |
| VT56 | 39,478,594 | 19,739,297 | 1 | 19,739,297 | 19,739,297 | 35,223,614 | 35,223,614 | 34,243,652 | 35,223,614 | 89.2200 | 89.2200 | 86.7400 | 89.2200 |
| VT70 | 36,501,706 | 18,250,853 | 1 | 18,250,853 | 18,250,853 | 36,501,706 | 35,071,055 | 26,998,025 | 33,212,216 | 100.0000 | 96.0800 | 73.9600 | 90.9900 |
| 82303 | 26,182,190 | 13,090,802 | 1 | 13,091,388 | 13,090,802 | 1,911,700 | 1,911,700 | 1,848,031 | 1,911,700 | 7.3000 | 7.3000 | 7.0600 | 7.3000 |
| 186035 | 1,575,600 | 787,800 | 1 | 787,800 | 787,800 | 1,155,767 | 1,155,767 | 1,121,634 | 1,155,767 | 73.3500 | 73.3500 | 71.1900 | 73.3500 |
| VT100 | 25,033,995 | 12,516,633 | 1 | 12,516,633 | 12,517,362 | 16,666,269 | 16,666,269 | 16,207,830 | 16,666,269 | 66.5700 | 66.5700 | 64.7400 | 66.5800 |
| 186036 | 1,549,182 | 774,591 | 1 | 774,591 | 774,591 | 948,814 | 948,814 | 917,777 | 948,814 | 61.2500 | 61.2500 | 59.2400 | 61.2500 |
| VT101 | 42,517,450 | 21,258,725 | 1 | 21,258,725 | 21,258,725 | 38,147,952 | 38,147,952 | 37,164,308 | 38,147,952 | 89.7200 | 89.7200 | 87.4100 | 89.7200 |
| 186037 | 1,180,434 | 590,217 | 1 | 590,217 | 590,217 | 695,488 | 695,488 | 673,873 | 695,488 | 58.9200 | 58.9200 | 57.0900 | 58.9200 |
| VT102 | 24,905,246 | 12,451,950 | 1 | 12,451,950 | 12,453,296 | 24,321,499 | 24,321,499 | 24,185,144 | 24,321,499 | 97.6600 | 97.6600 | 97.1100 | 97.6600 |
| VT103 | 41,827,002 | 20,913,501 | 1 | 20,913,501 | 20,913,501 | 37,501,601 | 37,501,601 | 36,500,560 | 37,501,601 | 89.6600 | 89.6600 | 87.2700 | 89.6600 |
| VT104 | 36,661,266 | 18,330,633 | 1 | 18,330,633 | 18,330,633 | 32,780,658 | 32,780,658 | 31,929,980 | 32,780,658 | 89.4100 | 89.4100 | 87.0900 | 89.4100 |
| VT105 | 46,991,776 | 23,495,888 | 1 | 23,495,888 | 23,495,888 | 2,737,066 | 2,737,066 | 2,547,708 | 2,737,066 | 5.8200 | 5.8200 | 5.4200 | 5.8200 |
| VT107 | 40,873,946 | 20,436,973 | 1 | 20,436,973 | 20,436,973 | 36,664,440 | 36,664,440 | 35,706,213 | 36,664,440 | 89.7000 | 89.7000 | 87.3600 | 89.7000 |
| VT109 | 34,852,314 | 17,426,157 | 1 | 17,426,157 | 17,426,157 | 31,522,870 | 31,522,870 | 30,758,711 | 31,522,870 | 90.4500 | 90.4500 | 88.2500 | 90.4500 |
| 12.21 | 29,337,764 | 14,668,882 | 1 | 14,668,882 | 14,668,882 | 25,930,403 | 25,930,403 | 25,278,606 | 25,930,403 | 88.3900 | 88.3900 | 86.1600 | 88.3900 |
| 12.23 | 34,959,560 | 17,479,780 | 1 | 17,479,780 | 17,479,780 | 29,091,549 | 29,091,549 | 27,998,645 | 29,091,549 | 83.2100 | 83.2100 | 80.0900 | 83.2100 |
| 12.25 | 29,488,730 | 14,744,365 | 1 | 14,744,365 | 14,744,365 | 25,932,737 | 25,932,737 | 25,191,416 | 25,932,737 | 87.9400 | 87.9400 | 85.4300 | 87.9400 |
| 12.26 | 30,395,994 | 15,197,997 | 1 | 15,197,997 | 15,197,997 | 27,063,599 | 27,063,599 | 26,332,706 | 27,063,599 | 89.0400 | 89.0400 | 86.6300 | 89.0400 |
| 10 | 20,242,036 | 10,121,018 | 1 | 10,121,018 | 10,121,018 | 16,892,286 | 16,892,286 | 16,332,048 | 16,892,286 | 83.4500 | 83.4500 | 80.6800 | 83.4500 |
| 12.4 | 31,702,668 | 15,851,334 | 1 | 15,851,334 | 15,851,334 | 26,788,839 | 26,788,839 | 25,961,654 | 26,788,839 | 84.5000 | 84.5000 | 81.8900 | 84.5000 |
| 11 | 3,945,284 | 1,972,642 | 1 | 1,972,642 | 1,972,642 | 3,100,785 | 3,100,785 | 3,021,539 | 3,100,785 | 78.5900 | 78.5900 | 76.5900 | 78.5900 |
| R-4404 | 25,071,376 | 12,535,398 | 1 | 12,535,398 | 12,535,978 | 1,382,282 | 1,382,282 | 1,351,416 | 1,382,282 | 5.5100 | 5.5100 | 5.3900 | 5.5100 |
| 12.43 | 33,157,842 | 16,578,921 | 1 | 16,578,921 | 16,578,921 | 29,680,985 | 29,680,985 | 28,918,251 | 29,680,985 | 89.5100 | 89.5100 | 87.2100 | 89.5100 |
| 11.04 | 25,425,380 | 12,712,690 | 1 | 12,712,690 | 12,712,690 | 22,363,533 | 22,363,533 | 21,750,109 | 22,363,533 | 87.9600 | 87.9600 | 85.5400 | 87.9600 |
| 11.07 | 30,088,014 | 15,044,007 | 1 | 15,044,007 | 15,044,007 | 26,956,930 | 26,956,930 | 26,315,959 | 26,956,930 | 89.5900 | 89.5900 | 87.4600 | 89.5900 |
| 12.61 | 31,947,908 | 15,973,954 | 1 | 15,973,954 | 15,973,954 | 27,553,866 | 27,553,866 | 26,687,217 | 27,553,866 | 86.2500 | 86.2500 | 83.5300 | 86.2500 |
| 12.48 | 19,862,948 | 9,931,474 | 1 | 9,931,474 | 9,931,474 | 17,331,193 | 17,331,193 | 16,913,856 | 17,331,193 | 87.2500 | 87.2500 | 85.1500 | 87.2500 |
| 11.4 | 40,722,168 | 20,361,084 | 1 | 20,361,084 | 20,361,084 | 35,863,103 | 35,863,103 | 34,741,385 | 35,863,103 | 88.0700 | 88.0700 | 85.3100 | 88.0700 |
| 162019 | 33,423,630 | 16,711,815 | 1 | 16,711,815 | 16,711,815 | 995,658 | 995,658 | 968,706 | 995,658 | 2.9800 | 2.9800 | 2.9000 | 2.9800 |
| 12.49 | 11,465,026 | 5,732,513 | 1 | 5,732,513 | 5,732,513 | 9,258,856 | 9,258,856 | 8,985,959 | 9,258,856 | 80.7600 | 80.7600 | 78.3800 | 80.7600 |
| 11.24 | 24,623,176 | 12,311,588 | 1 | 12,311,588 | 12,311,588 | 21,725,308 | 21,725,308 | 21,161,681 | 21,725,308 | 88.2300 | 88.2300 | 85.9400 | 88.2300 |
| 12.64 | 30,113,684 | 15,056,842 | 1 | 15,056,842 | 15,056,842 | 24,571,275 | 24,571,275 | 23,670,306 | 24,571,275 | 81.6000 | 81.6000 | 78.6000 | 81.6000 |
| 27544 | 35,890,692 | 17,945,346 | 1 | 17,945,346 | 17,945,346 | 1,424,797 | 1,424,797 | 1,387,398 | 1,424,797 | 3.9700 | 3.9700 | 3.8700 | 3.9700 |
| 11.28 | 37,510,038 | 18,755,019 | 1 | 18,755,019 | 18,755,019 | 33,450,422 | 33,450,422 | 32,591,714 | 33,450,422 | 89.1800 | 89.1800 | 86.8900 | 89.1800 |
| 11.41 | 40,710,252 | 20,355,126 | 1 | 20,355,126 | 20,355,126 | 36,417,733 | 36,417,733 | 35,472,452 | 36,417,733 | 89.4600 | 89.4600 | 87.1300 | 89.4600 |
| 11.29 | 29,259,462 | 14,629,731 | 1 | 14,629,731 | 14,629,731 | 25,337,291 | 25,337,291 | 24,596,248 | 25,337,291 | 86.6000 | 86.6000 | 84.0600 | 86.6000 |
| VT1 | 125,878,582 | 62,939,291 | 1 | 62,939,291 | 62,939,291 | 83,912,710 | 83,912,710 | 81,060,626 | 83,912,710 | 66.6600 | 66.6600 | 64.4000 | 66.6600 |
| 11.43 | 27,193,348 | 13,596,674 | 1 | 13,596,674 | 13,596,674 | 23,388,515 | 23,388,515 | 22,636,298 | 23,388,515 | 86.0100 | 86.0100 | 83.2400 | 86.0100 |
| 11.44 | 39,028,780 | 19,514,390 | 1 | 19,514,390 | 19,514,390 | 34,647,213 | 34,647,213 | 33,679,986 | 34,647,213 | 88.7700 | 88.7700 | 86.3000 | 88.7700 |
| 11.45 | 35,240,178 | 17,620,089 | 1 | 17,620,089 | 17,620,089 | 30,600,988 | 30,600,988 | 29,561,059 | 30,600,988 | 86.8400 | 86.8400 | 83.8800 | 86.8400 |
| VT4 | 39,522,840 | 19,761,420 | 1 | 19,761,420 | 19,761,420 | 35,281,426 | 35,281,426 | 34,348,547 | 35,281,426 | 89.2700 | 89.2700 | 86.9100 | 89.2700 |
| 11.46 | 43,277,470 | 21,638,735 | 1 | 21,638,735 | 21,638,735 | 38,841,259 | 38,841,259 | 37,853,966 | 38,841,259 | 89.7500 | 89.7500 | 87.4700 | 89.7500 |
| 10.21 | 15,249,038 | 7,624,519 | 1 | 7,624,519 | 7,624,519 | 10,905,072 | 10,905,072 | 10,588,328 | 10,905,072 | 71.5100 | 71.5100 | 69.4400 | 71.5100 |
| VT8 | 45,507,146 | 22,753,573 | 1 | 22,753,573 | 22,753,573 | 45,507,146 | 45,319,626 | 40,012,970 | 44,216,668 | 100.0000 | 99.5900 | 87.9300 | 97.1600 |
| VT20 | 10,284,758 | 5,142,379 | 1 | 5,142,379 | 5,142,379 | 7,146,886 | 7,146,886 | 6,911,267 | 7,146,886 | 69.4900 | 69.4900 | 67.2000 | 69.4900 |
| 11.64 | 16,018,750 | 8,009,375 | 1 | 8,009,375 | 8,009,375 | 12,531,649 | 12,531,649 | 12,064,757 | 12,531,649 | 78.2300 | 78.2300 | 75.3200 | 78.2300 |

### Table S4. Alignment metrics to C. horridus

Per-sample host alignment summary to the timber rattlesnake reference, mirroring S3 fields to enable direct comparison of mapping rates and Q30 retention by sample.

| snake | raw reads total | raw pairs | is paired | raw r1 | raw r2 | primary total in bam | primary mapped | primary mapped q30 | primary pp reads | percent input captured | pct mapped vs fastq | pct mapped q30 vs fastq | pct properly paired pair vs fastq |
|---|---|---|---|---|---|---|---|---|---|---|---|---|---|
| 10.4 | 38,940,546 | 19,470,273 | 1 | 19,470,273 | 19,470,273 | 35,011,202 | 35,011,202 | 34,144,616 | 35,011,202 | 89.9100 | 89.9100 | 87.6800 | 89.9100 |
| 90 | 38,805,910 | 19,402,955 | 1 | 19,402,955 | 19,402,955 | 34,197,856 | 34,197,856 | 33,150,144 | 34,197,856 | 88.1300 | 88.1300 | 85.4300 | 88.1300 |
| 77 | 25,153,460 | 12,576,171 | 1 | 12,576,171 | 12,577,289 | 23,899,270 | 23,899,270 | 23,569,606 | 23,899,270 | 95.0100 | 95.0100 | 93.7000 | 95.0200 |
| VT24 | 38,913,786 | 19,456,893 | 1 | 19,456,893 | 19,456,893 | 34,876,595 | 34,876,595 | 33,934,434 | 34,876,595 | 89.6300 | 89.6300 | 87.2000 | 89.6300 |
| 91 | 24,101,706 | 12,050,853 | 1 | 12,050,853 | 12,050,853 | 20,818,044 | 20,818,044 | 20,193,671 | 20,818,044 | 86.3800 | 86.3800 | 83.7900 | 86.3800 |
| 78 | 25,040,951 | 12,520,011 | 1 | 12,520,011 | 12,520,940 | 24,339,788 | 24,339,788 | 24,165,950 | 24,339,788 | 97.2000 | 97.2000 | 96.5100 | 97.2000 |
| 11.68 | 126,330,410 | 63,165,205 | 1 | 63,165,205 | 63,165,205 | 110,935,267 | 110,935,267 | 108,631,622 | 110,935,267 | 87.8100 | 87.8100 | 85.9900 | 87.8100 |
| 92 | 13,546,924 | 6,773,462 | 1 | 6,773,462 | 6,773,462 | 11,749,855 | 11,749,855 | 11,434,855 | 11,749,855 | 86.7300 | 86.7300 | 84.4100 | 86.7300 |
| 93 | 36,702,800 | 18,350,215 | 1 | 18,350,215 | 18,352,585 | 33,152,850 | 33,152,850 | 32,274,505 | 33,152,850 | 90.3300 | 90.3300 | 87.9300 | 90.3300 |
| VT40 | 11,413,826 | 5,706,913 | 1 | 5,706,913 | 5,706,913 | 9,238,890 | 9,238,890 | 8,985,171 | 9,238,890 | 80.9400 | 80.9400 | 78.7200 | 80.9400 |
| VT44 | 40,257,834 | 20,128,917 | 1 | 20,128,917 | 20,128,917 | 35,710,508 | 35,710,508 | 34,762,677 | 35,710,508 | 88.7000 | 88.7000 | 86.3500 | 88.7000 |
| 99 | 29,045,652 | 14,522,826 | 1 | 14,522,826 | 14,522,826 | 25,597,942 | 25,597,942 | 24,860,915 | 25,597,942 | 88.1300 | 88.1300 | 85.5900 | 88.1300 |
| 6462 | 46,817,378 | 23,408,689 | 1 | 23,408,689 | 23,408,689 | 2,081,230 | 2,081,230 | 2,026,227 | 2,081,230 | 4.4500 | 4.4500 | 4.3300 | 4.4500 |

**Table S5. Coverage & run information**

Per-sample read count, mean coverage (x), and sequencing run metadata (Run_ID, Lane, platform), with a flag indicating inclusion in host-genome analyses.

| snake | Read Count | Mean Coverage | Run ID | Lane | Retained for host analysis? |
|---|---|---|---|---|---|
| 1 | 18,270,090 | 1.2138 | 623 | 5 | TRUE |
| 2 | 11,432,133 | 0.6599 | 623 | 4 | TRUE |
| 5 | 15,102,263 | 1.0696 | 623 | 4 | TRUE |
| 6 | 13,104,403 | 0.9393 | 623 | 4 | TRUE |
| 7 | 16,520,504 | 1.0125 | 623 | 5 | TRUE |
| 8 | 11,898,856 | 0.7072 | 623 | 4 | TRUE |
| 9 | 20,964,356 | 1.5173 | 623 | 5 | TRUE |
| 10 | 10,121,018 | 0.5256 | 623 | 6 | TRUE |
| 10.16 | 14,341,170 | 0.7510 | 623 | 6 | TRUE |
| 10.17 | 7,355,539 | 0.3848 | 623 | 4 | TRUE |
| 10.21 | 7,624,519 | 0.3167 | 623 | 6 | TRUE |
| 10.34 | 6,490,002 | 0.3403 | 623 | 5 | TRUE |
| 10.4 | 19,470,273 | 1.5659 | 623 | 4 | TRUE |
| 11 | 1,972,642 | 0.0940 | 623 | 6 | TRUE |
| 11.04 | 12,712,690 | 0.7656 | 623 | 4 | TRUE |
| 11.07 | 15,044,007 | 1.1887 | 623 | 4 | TRUE |
| 11.1 | 2,121,514 | 0.0666 | 623 | 5 | TRUE |
| 11.24 | 12,311,588 | 0.7612 | 623 | 4 | TRUE |
| 11.28 | 18,755,019 | 1.2288 | 623 | 5 | TRUE |
| 11.29 | 14,629,731 | 0.8179 | 623 | 5 | TRUE |
| 11.3 | 16,357,485 | 1.1736 | 623 | 4 | TRUE |
| 11.31 | 20,828,149 | 1.6137 | 623 | 5 | TRUE |
| 11.32 | 17,640,724 | 1.1632 | 623 | 4 | TRUE |
| 11.35 | 17,716,691 | 1.0342 | 623 | 5 | TRUE |
| 11.36 | 15,004,276 | 0.8689 | 623 | 5 | TRUE |
| 11.38 | 17,254,584 | 1.2252 | 623 | 4 | TRUE |
| 11.4 | 20,361,084 | 1.3316 | 623 | 6 | TRUE |
| 11.41 | 20,355,126 | 1.4808 | 623 | 5 | TRUE |
| 11.43 | 13,596,674 | 0.8844 | 623 | 4 | TRUE |
| 11.44 | 19,514,390 | 1.5938 | 623 | 4 | TRUE |
| 11.45 | 17,620,089 | 1.3862 | 623 | 4 | TRUE |
| 11.46 | 21,638,735 | 1.5070 | 623 | 6 | TRUE |
| 11.52 | 19,848,014 | 1.4265 | 623 | 5 | TRUE |
| 11.53 | 14,035,115 | 0.9370 | 623 | 4 | TRUE |
| 11.54 | 20,015,116 | 1.3131 | 623 | 6 | TRUE |
| 11.57 | 21,259,635 | 1.6122 | 623 | 6 | TRUE |
| 11.58 | 7,875,940 | 0.2710 | 623 | 5 | TRUE |
| 11.64 | 8,009,375 | 0.4429 | 623 | 4 | TRUE |
| 11.68 | 63,165,205 | 3.6178 | 562 | 2 | TRUE |
| 11.73 | 4,854,868 | 0.1446 | 623 | 6 | TRUE |
| 11.74 | 12,574,872 | 0.5700 | 623 | 6 | TRUE |
| 12.11 | 6,063,027 | 0.2772 | 623 | 6 | TRUE |
| 12.19 | 14,492,408 | 0.9373 | 623 | 4 | TRUE |
| 12.21 | 14,668,882 | 0.9623 | 623 | 5 | TRUE |
| 12.23 | 17,479,780 | 1.1068 | 623 | 5 | TRUE |
| 12.25 | 14,744,365 | 1.0648 | 623 | 4 | TRUE |
| 12.26 | 15,197,997 | 1.1715 | 623 | 4 | TRUE |
| 12.32 | 14,381,065 | 0.8943 | 623 | 5 | TRUE |
| 12.34 | 6,461,590 | 0.2936 | 623 | 5 | TRUE |
| 12.39 | 18,708,911 | 1.4202 | 623 | 4 | TRUE |
| 12.4 | 15,851,334 | 0.9336 | 623 | 5 | TRUE |
| 12.43 | 16,578,921 | 1.0395 | 623 | 6 | TRUE |
| 12.48 | 9,931,474 | 0.5822 | 623 | 4 | TRUE |
| 12.49 | 5,732,513 | 0.3114 | 623 | 5 | TRUE |
| 12.5 | 2,065,191 | 0.0898 | 623 | 5 | TRUE |
| 12.51 | 216,819 | 0.0023 | 623 | 4 | FALSE |
| 12.52 | 12,175,528 | 0.6570 | 623 | 5 | TRUE |
| 12.61 | 15,973,954 | 1.0740 | 623 | 4 | TRUE |
| 12.64 | 15,056,842 | 0.8384 | 623 | 6 | TRUE |
| 12.71 | 16,211,455 | 1.0847 | 623 | 5 | TRUE |
| 12.73 | 34,353,152 | 1.0515 | 562 | 2 | TRUE |
| 12.74 | 3,587,058 | 0.0079 | 562 | 2 | FALSE |
| 12.75 | 5,617,411 | 0.1683 | 562 | 2 | TRUE |
| 77 | 12,576,171 | 0.8310 | 623 | 6 | TRUE |
| 78 | 12,520,011 | 1.0301 | 623 | 6 | TRUE |
| 80 | 2,646,826 | 0.1684 | 623 | 6 | TRUE |
| 81 | 18,440,072 | 0.0411 | 623 | 6 | FALSE |
| 84 | 12,169,095 | 0.8004 | 623 | 6 | TRUE |

**Table S5. Coverage & run information**

Per-sample read count, mean coverage (x), and sequencing run metadata (Run_ID, Lane, platform), with a flag indicating inclusion in host-genome analyses.

| snake | Read Count | Mean Coverage | Run ID | Lane | Retained for host analysis? |
|---|---|---|---|---|---|
| 90 | 19,402,955 | 1.4616 | 623 | 6 | TRUE |
| 91 | 12,050,853 | 0.7209 | 623 | 6 | TRUE |
| 92 | 6,773,462 | 0.4211 | 623 | 6 | TRUE |
| 93 | 18,350,215 | 1.4288 | 623 | 6 | TRUE |
| 99 | 14,522,826 | 0.9241 | 623 | 6 | TRUE |
| 6439 | 6,681,394 | 0.0024 | 623 | 6 | FALSE |
| 6462 | 23,408,689 | 0.0063 | 623 | 6 | FALSE |
| 19192 | 26,181,960 | 0.0064 | 623 | 6 | FALSE |
| 27544 | 17,945,346 | 0.0043 | 623 | 6 | FALSE |
| 82303 | 13,091,388 | 0.0057 | 623 | 6 | FALSE |
| 162019 | 16,711,815 | 0.0029 | 623 | 6 | FALSE |
| 186035 | 787,800 | 0.0324 | 623 | 6 | FALSE |
| 186036 | 774,591 | 0.0275 | 623 | 6 | FALSE |
| 186037 | 590,217 | 0.0193 | 623 | 6 | FALSE |
| 09.06SFD | 16,530,729 | 1.2303 | 623 | 5 | TRUE |
| 3 | 12,909,230 | 0.7633 | 623 | 5 | TRUE |
| R-4404 | 12,535,398 | 0.0040 | 623 | 6 | FALSE |
| VT1 | 62,939,291 | 1.7031 | 562 | 2 | TRUE |
| VT10 | 58,590,677 | 1.9742 | 562 | 2 | TRUE |
| VT100 | 12,516,633 | 0.6697 | 623 | 4 | TRUE |
| VT101 | 21,258,725 | 1.6909 | 623 | 5 | TRUE |
| VT102 | 12,451,950 | 1.0458 | 623 | 6 | TRUE |
| VT103 | 20,913,501 | 1.7012 | 623 | 5 | TRUE |
| VT104 | 18,330,633 | 1.4682 | 623 | 4 | TRUE |
| VT105 | 23,495,888 | 0.0121 | 623 | 6 | FALSE |
| VT107 | 20,436,973 | 1.4978 | 623 | 5 | TRUE |
| VT109 | 17,426,157 | 1.3148 | 623 | 4 | TRUE |
| VT110 | 18,250,211 | 1.3239 | 623 | 4 | TRUE |
| VT111 | 11,175,296 | 0.6236 | 623 | 5 | TRUE |
| VT12 | 16,339,102 | 0.9394 | 623 | 5 | TRUE |
| VT16 | 23,525,427 | 1.9258 | 623 | 5 | TRUE |
| VT20 | 5,142,379 | 0.1364 | 562 | 2 | TRUE |
| VT24 | 19,456,893 | 1.7415 | 623 | 4 | TRUE |
| VT30 | 31,212,924 | 1.3658 | 562 | 2 | TRUE |
| VT34 | 21,864,530 | 1.6948 | 623 | 6 | TRUE |
| VT4 | 19,761,420 | 1.6619 | 623 | 4 | TRUE |
| VT40 | 5,706,913 | 0.1939 | 562 | 2 | TRUE |
| VT44 | 20,128,917 | 1.6395 | 623 | 5 | TRUE |
| VT50 | 21,899,301 | 1.0789 | 562 | 2 | TRUE |
| VT52 | 17,471,624 | 1.3858 | 623 | 4 | TRUE |
| VT56 | 19,739,297 | 1.7069 | 623 | 4 | TRUE |
| VT70 | 18,250,853 | 0.5917 | 562 | 2 | TRUE |
| VT8 | 22,753,573 | 1.9752 | 623 | 5 | TRUE |

**Table S6. Pairwise population differentiation**

Genome-wide weighted FST for each population pair, with great-circle distances (km) included for isolation-by-distance analyses.

| PopA | PopB | FST unweighted | FST Weighted | Distance km |
|---|---|---|---|---|
| Rutland VT L1 | Westfield MA | 0.0323 | 0.0326 | 188.7000 |
| Copake NY | Mt Washington MA | 0.0140 | 0.0475 | 4.6000 |
| Central PA | Mt Washington MA | 0.0286 | 0.0970 | 325.7000 |
| Mt Washington MA | Westfield MA | 0.0269 | 0.1021 | 64.0000 |
| Central PA | Copake NY | 0.0453 | 0.1105 | 322.8000 |
| Copake NY | Westfield MA | 0.0397 | 0.1201 | 67.9000 |
| Mt Washington MA | Rutland VT L2 | 0.0172 | 0.1405 | 170.5000 |
| Mt Washington MA | Rutland VT L1 | 0.0161 | 0.1433 | 189.7000 |
| Rutland VT L2 | Westfield MA | 0.0466 | 0.1467 | 159.7000 |
| Central PA | Westfield MA | 0.0508 | 0.1481 | 387.4000 |
| Copake NY | Rutland VT L2 | 0.0437 | 0.1614 | 169.6000 |
| Copake NY | Rutland VT L1 | 0.0445 | 0.1640 | 188.1000 |
| Central PA | Rutland VT L2 | 0.0644 | 0.1924 | 450.3000 |
| Central PA | Rutland VT L1 | 0.0712 | 0.1934 | 446.5000 |
| Rutland VT L1 | Rutland VT L2 | 0.0503 | 0.2034 | 36.5000 |

**Table S7. Individual heterozygosity and ROH burden**

Per-individual heterozygosity (genome-wide and autosomes), genomic inbreeding from ROH (FROH>100kb and FROH>1Mb), plus population labels and per-1000bp summaries.

| snake | het sites | total sites | het per 1000 | F ROH 100kb | F ROH 1Mb | het sites (XY excluded) | total sites (XY excluded) | het per 1000 (XY excluded) | het rate (XY excluded) |
|---|---|---|---|---|---|---|---|---|---|
| 09.06SFD | 1064962.9158 | 922796440.0216 | 1.1541 | 0.09502895 | 0.01425527 | 941596.1682 | 837423581.1037 | 1.1244 | 0.0011 |
| 1 | 1208217.9015 | 860453028.9808 | 1.4042 | 0.08667827 | 0 | 1003553.2583 | 784037066.9685 | 1.2800 | 0.0013 |
| 10 | 835283.1604 | 514468309.8503 | 1.6236 | 0.02087751 | 0 | 698472.3196 | 470218357.8937 | 1.4854 | 0.0015 |
| 10.16 | 1515454.5356 | 647815133.8910 | 2.3393 | 0.02923279 | 0 | 1302075.0708 | 591568180.8278 | 2.2011 | 0.0022 |
| 10.17 | 705598.5303 | 390416979.0379 | 1.8073 | 0.04433498 | 0 | 618848.2618 | 353417301.9674 | 1.7510 | 0.0018 |
| 10.21 | 684286.5191 | 322627108.9508 | 2.1210 | 0.0315904 | 0 | 597886.8957 | 291555076.9684 | 2.0507 | 0.0021 |
| 10.34 | 320480.1066 | 354305292.1152 | 0.9045 | 0.08128832 | 0.00337557 | 275700.1185 | 327325470.0740 | 0.8423 | 0.0008 |
| 10.4 | 1338613.8888 | 1030612382.2720 | 1.2989 | 0.05799261 | 0.00136799 | 1192228.0769 | 935147409.2196 | 1.2749 | 0.0013 |
| 11 | 103558.8997 | 113179399.0000 | 0.9150 | NA | NA | 92022.3945 | 102555238.0000 | 0.8973 | 0.0009 |
| 11.04 | 928946.7017 | 691079453.9823 | 1.3442 | 0.03664345 | 0.00566624 | 819188.8252 | 626900152.9823 | 1.3067 | 0.0013 |
| 11.07 | 1105137.8472 | 906609080.4784 | 1.2190 | 0.0712808 | 0.01163938 | 979536.3725 | 822778942.3228 | 1.1905 | 0.0012 |
| 11.1 | 23319.5037 | 62260527.9986 | 0.3745 | NA | NA | 20750.1013 | 56345513.9991 | 0.3683 | 0.0004 |
| 11.24 | 1316146.4193 | 694415284.8171 | 1.8953 | 0.02651646 | 0 | 1164509.1849 | 629986780.7023 | 1.8485 | 0.0018 |
| 11.28 | 1201426.5541 | 931061502.3287 | 1.2904 | 0.05322351 | 0.00852261 | 1056173.0926 | 844537900.7763 | 1.2506 | 0.0013 |
| 11.29 | 1842920.1578 | 682699823.9541 | 2.6995 | 0.00237535 | 0 | 1632307.4484 | 618985286.7724 | 2.6371 | 0.0026 |
| 11.3 | 1420922.9886 | 850479249.7803 | 1.6707 | 0.05212634 | 0 | 1205004.0980 | 775759338.2406 | 1.5533 | 0.0016 |
| 11.31 | 1390336.5797 | 1027660775.9793 | 1.3529 | 0.04164099 | 0 | 1187221.5723 | 934293174.0159 | 1.2707 | 0.0013 |
| 11.32 | 1222101.6726 | 884892267.7244 | 1.3811 | 0.06902643 | 0 | 1027107.6601 | 807639242.0877 | 1.2717 | 0.0013 |
| 11.35 | 1529217.3130 | 650023313.7876 | 2.3526 | 0.02784937 | 0.00596156 | 1333093.9837 | 586757993.0834 | 2.2720 | 0.0023 |
| 11.36 | 892278.7121 | 752127000.9707 | 1.1863 | 0.05433335 | 0.00249102 | 779793.7354 | 682330632.1652 | 1.1428 | 0.0011 |
| 11.38 | 1167292.6233 | 935037295.0031 | 1.2484 | 0.08345182 | 0 | 1024729.9165 | 848427394.1604 | 1.2078 | 0.0012 |
| 11.4 | 1550167.6305 | 938888194.4445 | 1.6511 | 0.03831404 | 0.00131046 | 1316327.6438 | 855167686.3333 | 1.5393 | 0.0015 |
| 11.41 | 1290046.5924 | 1001169672.8132 | 1.2885 | 0.0607909 | 0.00592976 | 1133389.5568 | 907955006.2810 | 1.2483 | 0.0012 |
| 11.43 | 1251054.1404 | 728244752.0951 | 1.7179 | 0.05102941 | 0 | 1064464.6092 | 665951836.1368 | 1.5984 | 0.0016 |
| 11.44 | 3025042.5173 | 1000010163.7477 | 3.0250 | 0.00122813 | 0 | 2665725.7719 | 909430508.3657 | 2.9312 | 0.0029 |
| 11.45 | 1552279.9066 | 957111940.2698 | 1.6218 | 0.04072221 | 0 | 1323161.6190 | 870700608.4110 | 1.5197 | 0.0015 |
| 11.46 | 1288811.9781 | 1009559061.8587 | 1.2766 | 0.06202942 | 0.00720309 | 1133857.0443 | 915732488.9093 | 1.2382 | 0.0012 |
| 11.52 | 1286891.8356 | 985278713.4951 | 1.3061 | 0.04382784 | 0.00224509 | 1145745.4558 | 893744970.7136 | 1.2820 | 0.0013 |
| 11.53 | 1456025.0440 | 805028247.7520 | 1.8087 | 0.01361191 | 0 | 1271562.4195 | 737881410.2954 | 1.7233 | 0.0017 |
| 11.54 | 1538679.7197 | 883557660.9466 | 1.7415 | 0.03425362 | 0 | 1359654.3038 | 802071611.8653 | 1.6952 | 0.0017 |
| 11.57 | 1459087.4707 | 1014332312.0006 | 1.4385 | 0.03812487 | 0 | 1291956.3602 | 920379188.5187 | 1.4037 | 0.0014 |
| 11.58 | 588607.0322 | 262984881.9869 | 2.2382 | 0.02497998 | 0.00184711 | 495059.5659 | 238120736.0413 | 2.0790 | 0.0021 |
| 11.64 | 920847.5573 | 448734821.0000 | 2.0521 | 0.02002107 | 0 | 809814.5330 | 406204458.1292 | 1.9936 | 0.0020 |
| 11.68 | 1991659.8701 | 1233039942.9602 | 1.6152 | 0.02315154 | 0 | 1759280.3160 | 1116462032.1993 | 1.5758 | 0.0016 |
| 11.73 | 205852.3227 | 171021321.0000 | 1.2037 | NA | NA | 180908.2409 | 154405759.0000 | 1.1716 | 0.0012 |
| 11.74 | 1214414.3909 | 480857855.0280 | 2.5255 | NA | NA | 1029859.6334 | 436826975.0769 | 2.3576 | 0.0024 |
| 3 | 994967.6568 | 699792683.2455 | 1.4218 | 0.0255754 | 0 | 849431.5770 | 641835786.0000 | 1.3234 | 0.0013 |
| 12.11 | 564155.5529 | 268970434.0025 | 2.0975 | 0.06314285 | 0.00779778 | 470974.6194 | 243457450.0535 | 1.9345 | 0.0019 |
| 12.19 | 1541050.0259 | 706522387.6960 | 2.1812 | 0.03634773 | 0 | 1303072.7332 | 641810609.2542 | 2.0303 | 0.0020 |
| 12.21 | 1172862.3144 | 798233744.3145 | 1.4693 | 0.03641122 | 0.00190126 | 1045060.4687 | 724661726.2922 | 1.4421 | 0.0014 |
| 12.23 | 1420846.2095 | 763746745.8819 | 1.8604 | 0.04436456 | 0.001323 | 1247967.8959 | 692255600.9586 | 1.8028 | 0.0018 |
| 12.25 | 1423465.2251 | 830960724.8497 | 1.7130 | 0.03408402 | 0 | 1225739.7436 | 759667334.0863 | 1.6135 | 0.0016 |
| 12.26 | 1336933.0113 | 893964314.3404 | 1.4955 | 0.05985936 | 0 | 1142093.6373 | 816730549.0851 | 1.3984 | 0.0014 |
| 12.32 | 1168149.0628 | 744258826.8680 | 1.5695 | 0.07143483 | 0.00568005 | 1026241.4830 | 674851256.9532 | 1.5207 | 0.0015 |
| 12.34 | 383191.4733 | 313688177.0713 | 1.2216 | 0.06603271 | 0.00550239 | 337229.1159 | 284084470.0301 | 1.1871 | 0.0012 |
| 12.39 | 1306595.6314 | 970146630.9712 | 1.3468 | 0.07979532 | 0.00582193 | 1165759.0726 | 880417670.1990 | 1.3241 | 0.0013 |
| 12.4 | 1325333.1966 | 741717348.0781 | 1.7868 | 0.04277385 | 0 | 1169316.1384 | 672587725.0112 | 1.7385 | 0.0017 |
| 12.43 | 1249585.6303 | 861922326.8511 | 1.4498 | 0.0367812 | 0 | 1110142.5931 | 782209784.2323 | 1.4192 | 0.0014 |
| 12.48 | 728434.8741 | 554808125.1517 | 1.3129 | 0.06521207 | 0 | 643097.7485 | 503110595.8450 | 1.2782 | 0.0013 |
| 12.49 | 383356.7257 | 354695004.9318 | 1.0808 | 0.0820187 | 0.00343278 | 330682.3899 | 327431456.9931 | 1.0099 | 0.0010 |
| 12.5 | 107260.3854 | 87373411.0025 | 1.2276 | NA | NA | 93226.7754 | 79947676.0018 | 1.1661 | 0.0012 |
| 12.52 | 1075735.8404 | 585196224.9371 | 1.8382 | 0.05287912 | 0.00073032 | 907051.5734 | 534685510.2358 | 1.6964 | 0.0017 |
| 12.61 | 1401736.9044 | 817322087.9210 | 1.7150 | 0.0289982 | 0.00133594 | 1239677.5336 | 742066833.8509 | 1.6706 | 0.0017 |
| 12.64 | 1326451.6722 | 664370829.8032 | 1.9966 | 0.03851355 | 0 | 1116664.7978 | 603662033.9735 | 1.8498 | 0.0018 |
| 12.71 | 1301669.8787 | 873636343.3157 | 1.4899 | 0.02267091 | 0 | 1157676.3018 | 792281849.8389 | 1.4612 | 0.0015 |
| 12.73 | 1507599.9766 | 737627652.9278 | 2.0438 | 0.03628319 | 0 | 1279365.7535 | 673220513.9177 | 1.9004 | 0.0019 |
| 12.75 | 230614.5519 | 165796283.0009 | 1.3910 | NA | NA | 203324.5444 | 149968983.9964 | 1.3558 | 0.0014 |
| 2 | 837337.8862 | 619647689.2328 | 1.3513 | 0.01957551 | 0 | 735339.4357 | 562062235.9731 | 1.3083 | 0.0013 |
| 5 | 1126147.9754 | 859148563.5473 | 1.3108 | 0.02671578 | 0 | 995856.1804 | 779489674.1299 | 1.2776 | 0.0013 |
| 6 | 1084705.6233 | 784705321.3596 | 1.3823 | 0.03129782 | 0 | 951347.0181 | 711844561.0450 | 1.3365 | 0.0013 |
| 7 | 1081745.9381 | 816267105.1292 | 1.3252 | 0.03732185 | 0.00069076 | 950281.7356 | 740447078.8911 | 1.2834 | 0.0013 |
| 77 | 914809.9526 | 743109683.9054 | 1.2311 | 0.07804464 | 0.00644009 | 769171.4903 | 682308358.0421 | 1.1273 | 0.0011 |
| 78 | 791611.4396 | 715769598.1577 | 1.1060 | 0.06941555 | 0.00231533 | 686194.7513 | 654995491.9166 | 1.0476 | 0.0010 |
| 8 | 828355.4325 | 644662927.3063 | 1.2849 | 0.03006725 | 0.00116327 | 706100.0263 | 592028766.0306 | 1.1927 | 0.0012 |
| 80 | 162330.8420 | 202392733.0000 | 0.8021 | NA | NA | 138077.9734 | 186964584.0000 | 0.7385 | 0.0007 |
| 84 | 944392.4557 | 743321781.0934 | 1.2705 | 0.04295232 | 0 | 834680.9284 | 674403950.7852 | 1.2377 | 0.0012 |
| 9 | 1336112.1741 | 1010681510.8764 | 1.3220 | 0.03070339 | 0 | 1183606.6726 | 916885908.4335 | 1.2909 | 0.0013 |
| 90 | 1293840.7944 | 962916412.6762 | 1.3437 | 0.08128266 | 0.00455041 | 1072147.4654 | 875690789.9316 | 1.2243 | 0.0012 |
| 91 | 942695.2270 | 684091058.8479 | 1.3780 | 0.03622671 | 0 | 823618.0604 | 620242132.9693 | 1.3279 | 0.0013 |
| 92 | 470134.0850 | 459484257.2065 | 1.0232 | 0.0706874 | 0.01298134 | 406405.2500 | 416544754.1972 | 0.9757 | 0.0010 |
| 93 | 1256444.3979 | 976928578.8021 | 1.2861 | 0.05854771 | 0 | 1100380.8673 | 886708219.8858 | 1.2410 | 0.0012 |
| 99 | 1015885.0640 | 788094656.0057 | 1.2890 | 0.1299526 | 0.0052458 | 846261.2382 | 721360534.9544 | 1.1731 | 0.0012 |
| VT1 | 1548250.2593 | 730652172.0100 | 2.1190 | 0.03310315 | 0.00282128 | 1331342.2730 | 660786737.0354 | 2.0148 | 0.0020 |
| VT10 | 1699211.5659 | 970088779.5584 | 1.7516 | 0.02684075 | 0.00270118 | 1407800.6845 | 879461442.3823 | 1.6008 | 0.0016 |
| VT100 | 614392.1429 | 649292863.9752 | 0.9462 | 0.1322243 | 0.00709152 | 517101.0333 | 597956375.8379 | 0.8648 | 0.0009 |
| VT101 | 1066859.9555 | 1035592625.4981 | 1.0302 | 0.128741 | 0.00727266 | 945233.9288 | 939809854.3293 | 1.0058 | 0.0010 |
| VT102 | 657016.0286 | 702651846.8946 | 0.9351 | 0.1903466 | 0.01557562 | 572704.4344 | 643296624.0440 | 0.8903 | 0.0009 |
| VT103 | 1308903.7353 | 1049837615.3260 | 1.2468 | 0.08505003 | 0 | 1105485.3851 | 953626161.9929 | 1.1592 | 0.0012 |
| VT104 | 1184229.8018 | 999832528.1759 | 1.1844 | 0.08941763 | 0.0030834 | 1001805.6695 | 911044051.0056 | 1.0996 | 0.0011 |
| VT107 | 1881674.1375 | 1011791765.8739 | 1.8597 | 0.03526324 | 0.00572164 | 1619761.9623 | 921107808.8960 | 1.7585 | 0.0018 |
| VT109 | 1105464.8622 | 963010197.4652 | 1.1479 | 0.0898673 | 0.00184678 | 974463.5203 | 873883550.9958 | 1.1151 | 0.0011 |
| VT110 | 1180882.3542 | 973307427.8705 | 1.2133 | 0.1046837 | 0.00183451 | 996050.9078 | 887946168.4147 | 1.1217 | 0.0011 |
| VT111 | 819684.7206 | 594221233.9740 | 1.3794 | 0.08591444 | 0.00595321 | 709866.9984 | 547358356.8715 | 1.2969 | 0.0013 |
| VT12 | 1248106.6404 | 709207978.2119 | 1.7599 | 0.05477448 | 0.00270042 | 1047257.0334 | 645369759.0353 | 1.6227 | 0.0016 |
| VT16 | 1663285.8496 | 1069831525.8119 | 1.5547 | 0.04740482 | 0.00580236 | 1417016.6339 | 969816353.9484 | 1.4611 | 0.0015 |
| VT20 | 113370.0241 | 114726505.0002 | 0.9882 | NA | NA | 96860.4791 | 103101848.9938 | 0.9395 | 0.0009 |
| VT24 | 1184722.7447 | 1033728577.7329 | 1.1461 | 0.0766437 | 0 | 1061713.9030 | 938253596.0100 | 1.1316 | 0.0011 |
| VT30 | 1411722.0670 | 927145343.3013 | 1.5227 | 0.04178236 | 0.00598354 | 1238325.5309 | 840641945.2060 | 1.4731 | 0.0015 |
| VT34 | 1331852.0543 | 1052022272.6376 | 1.2660 | 0.04929283 | 0 | 1175992.6862 | 954296753.9895 | 1.2323 | 0.0012 |
| VT4 | 1210076.1335 | 1024112952.6014 | 1.1816 | 0.09302165 | 0.00891193 | 1053706.6694 | 929355902.4509 | 1.1338 | 0.0011 |
| VT40 | 169857.0389 | 180254388.0177 | 0.9423 | NA | NA | 141459.4738 | 164951838.0071 | 0.8576 | 0.0009 |
| VT44 | 1141798.6847 | 1041132590.0788 | 1.0967 | 0.1142271 | 0.00225087 | 946641.5358 | 946347460.1116 | 1.0003 | 0.0010 |
| VT50 | 966441.9057 | 851764438.5074 | 1.1346 | 0.06562203 | 0.00472486 | 848825.3118 | 772406781.5553 | 1.0989 | 0.0011 |
| VT52 | 1062849.4271 | 973144455.1082 | 1.0922 | 0.08309617 | 0.00908068 | 939561.7810 | 883149204.3598 | 1.0639 | 0.0011 |
| VT56 | 1147987.3915 | 1014877244.2558 | 1.1312 | 0.1055216 | 0.00576762 | 1005752.2633 | 921320532.9320 | 1.0916 | 0.0011 |
| VT70 | 1021616.1737 | 366874887.0054 | 2.7846 | 0.03070803 | 0.00076372 | 836318.4712 | 328194193.9969 | 2.5482 | 0.0025 |
| VT8 | 1306963.9931 | 1072147968.2466 | 1.2190 | 0.09573419 | 0.00133267 | 1093221.2443 | 971163041.5111 | 1.1257 | 0.0011 |

**Table S8. Population nucleotide diversity (π)**

Genome-wide and mitochondrial nucleotide diversity (π) per population, allowing autosomal vs. mitochondrial diversity comparisons.

| Pop | genomewide pi |
|---|---|
| Westfield_MA | 0.0018 |
| Central_PA | 0.0019 |
| Rutland VT L1 | 0.0018 |
| Rutland VT L2 | 0.0016 |
| Copake NY | 0.0021 |
| Rockland_County_NY | 0.0020 |
| Mt_Washington_MA | 0.0023 |

**Table S9. Between-population ROH concordance**

Frequency-weighted Jaccard similarity of ROH locations between population pairs at 100 kb and 1 Mb window scales; higher values indicate more similar ROH landscapes.

| popA | popB | Weighted Jaccard 100kb | Weighted Jaccard 1mb |
|---|---|---|---|
| Rutland VT L2 | Rutland VT L1 | 0.505 | 0.697 |
| Rutland VT L2 | Central PA | 0.292 | 0.532 |
| Rutland VT L2 | Copake NY | 0.360 | 0.618 |
| Rutland VT L2 | Mt Washington MA | 0.403 | 0.613 |
| Rutland VT L2 | Westfield MA | 0.423 | 0.583 |
| Rutland VT L1 | Central PA | 0.328 | 0.589 |
| Rutland VT L1 | Copake NY | 0.405 | 0.682 |
| Rutland VT L1 | Mt Washington MA | 0.467 | 0.692 |
| Rutland VT L1 | Westfield MA | 0.467 | 0.656 |
| Central PA | Copake NY | 0.356 | 0.637 |
| Central PA | Mt Washington MA | 0.423 | 0.676 |
| Central PA | Westfield MA | 0.352 | 0.629 |
| Copake NY | Mt Washington MA | 0.485 | 0.723 |
| Copake NY | Westfield MA | 0.396 | 0.660 |
| Mt Washington MA | Westfield MA | 0.472 | 0.695 |

**Table S10. PCA scores (all PCs)**

Per-sample principal component scores (PC1–PC97) from genotype covariance (including sex-chromosome PCs if present), used for structure plots and covariate modeling.

| snake | PC1 | PC2 | PC3 | PC4 | PC5 | PC6 | PC7 | PC8 | PC9 | PC10 |
|---|---|---|---|---|---|---|---|---|---|---|
| 09.06SFD | 0.1028 | 0.0570 | 0.0724 | 0.0069 | -0.0519 | 0.1371 | 0.0748 | -0.0210 | 0.0688 | -0.0849 |
| 1 | -0.1191 | 0.0506 | 0.0514 | -0.0545 | -0.1322 | -0.1824 | -0.2590 | -0.0793 | -0.0761 | -0.2296 |
| 10 | -0.0911 | 0.0782 | -0.0656 | -0.0902 | -0.1208 | -0.1223 | -0.1935 | -0.1138 | -0.0170 | -0.2056 |
| 10.16 | -0.1055 | 0.0969 | 0.0353 | -0.0956 | 0.2859 | -0.0230 | -0.0238 | -0.0383 | -0.0566 | 0.0103 |
| 10.17 | 0.1015 | 0.0581 | -0.0507 | -0.0409 | -0.0646 | 0.0226 | 0.0196 | 0.0250 | 0.0330 | 0.0568 |
| 10.21 | 0.1024 | 0.0734 | -0.0889 | -0.0984 | 0.1668 | -0.0181 | 0.0034 | 0.0349 | 0.0274 | 0.0163 |
| 10.34 | -0.0490 | 0.1052 | -0.1879 | -0.1208 | 0.2267 | 0.0285 | 0.0047 | -0.1454 | 0.0282 | 0.0525 |
| 10.4 | 0.0976 | 0.0503 | 0.0458 | 0.0591 | -0.0985 | 0.0182 | -0.0174 | -0.0150 | -0.0184 | 0.1050 |
| 11 | 0.1081 | 0.0365 | -0.1285 | -0.0651 | -0.0348 | -0.0197 | -0.0402 | -0.0008 | 0.0219 | -0.0749 |
| 11.04 | 0.0995 | 0.0642 | -0.0239 | -0.0211 | 0.0299 | -0.0300 | -0.0289 | 0.0184 | -0.0140 | 0.0618 |
| 11.07 | 0.1028 | 0.0637 | 0.0553 | 0.0036 | -0.0414 | 0.1458 | 0.0824 | -0.0328 | 0.0735 | -0.0910 |
| 11.1 | 0.1144 | 0.0357 | -0.1017 | -0.0715 | 0.0734 | 0.0388 | 0.0298 | 0.0223 | 0.0201 | -0.0204 |
| 11.24 | 0.0992 | 0.0595 | -0.0156 | -0.0096 | -0.0429 | 0.0135 | 0.0100 | 0.0271 | 0.0203 | 0.0294 |
| 11.28 | 0.1012 | 0.0597 | 0.0276 | 0.0180 | 0.1531 | -0.0061 | -0.0243 | -0.0209 | -0.0542 | 0.0205 |
| 11.29 | 0.0494 | 0.0768 | 0.0523 | -0.0518 | 0.1479 | 0.0061 | 0.0292 | 0.0445 | 0.0310 | -0.0655 |
| 11.3 | -0.1058 | 0.0876 | 0.0996 | 0.0102 | 0.1796 | 0.0406 | -0.0038 | -0.0703 | -0.0852 | 0.0203 |
| 11.31 | -0.1007 | 0.0670 | 0.0765 | 0.0521 | -0.1161 | 0.0077 | -0.0286 | -0.0717 | -0.0929 | 0.2049 |
| 11.32 | -0.1082 | 0.0950 | 0.0948 | -0.0317 | 0.2975 | -0.0150 | -0.0567 | -0.0561 | -0.1301 | 0.0631 |
| 11.35 | 0.0901 | 0.0251 | 0.1471 | -0.0381 | 0.1545 | -0.0570 | -0.0138 | 0.1160 | -0.0286 | 0.0251 |
| 11.36 | 0.1016 | 0.0666 | 0.0031 | -0.0176 | 0.1594 | 0.0024 | -0.0108 | -0.0048 | -0.0253 | -0.0100 |
| 11.38 | 0.0975 | 0.0747 | 0.0288 | 0.0356 | 0.1987 | 0.0018 | -0.0278 | -0.0370 | -0.0783 | 0.0343 |
| 11.4 | -0.1118 | 0.0767 | 0.1269 | 0.0050 | 0.2496 | 0.0190 | -0.0538 | -0.0786 | -0.1429 | 0.0521 |
| 11.41 | 0.0997 | 0.0539 | 0.0751 | 0.0212 | 0.1314 | -0.0058 | -0.0284 | 0.0031 | -0.0613 | 0.0407 |
| 11.43 | -0.1014 | 0.0930 | 0.0313 | -0.0423 | 0.2635 | -0.0054 | -0.0341 | -0.0979 | -0.0777 | -0.0009 |
| 11.44 | -0.1109 | 0.0656 | 0.1244 | 0.0260 | -0.1289 | 0.0850 | 0.0159 | -0.0825 | -0.0210 | 0.0728 |
| 11.45 | -0.1134 | 0.0813 | 0.1809 | 0.0212 | -0.0792 | 0.2484 | 0.1313 | -0.1186 | 0.0959 | -0.2299 |
| 11.46 | 0.0930 | 0.0580 | 0.0473 | 0.0331 | 0.1496 | -0.0152 | -0.0288 | -0.0194 | -0.0601 | 0.0368 |
| 11.52 | 0.0970 | 0.0467 | 0.0460 | 0.0405 | -0.0807 | -0.0239 | -0.0172 | 0.0250 | -0.0333 | 0.0808 |
| 11.53 | -0.0821 | 0.0843 | -0.0055 | 0.0137 | -0.0998 | 0.0679 | 0.0490 | -0.0985 | 0.0163 | 0.1559 |
| 11.54 | 0.0949 | 0.0357 | 0.0911 | 0.0523 | -0.1016 | -0.0206 | -0.0108 | 0.0561 | -0.0297 | 0.1137 |
| 11.57 | 0.0982 | 0.0361 | 0.0613 | 0.0509 | -0.1080 | -0.0190 | -0.0465 | -0.0018 | -0.0337 | 0.0867 |
| 11.58 | -0.0769 | 0.0945 | -0.0971 | -0.1208 | -0.1125 | -0.0159 | 0.0320 | 0.0192 | 0.1001 | 0.1618 |
| 11.64 | 0.1011 | 0.0685 | -0.0131 | -0.0374 | 0.0131 | 0.0578 | 0.0456 | 0.0253 | 0.0741 | -0.0548 |
| 11.68 | 0.0881 | 0.0072 | 0.2166 | 0.0117 | -0.0719 | -0.0420 | 0.0135 | 0.1336 | -0.0103 | 0.0910 |
| 11.73 | 0.1077 | 0.0426 | -0.1298 | -0.0631 | 0.0065 | 0.0164 | 0.0092 | 0.0177 | 0.0453 | 0.0136 |
| 11.74 | -0.1199 | 0.0539 | 0.0439 | -0.1113 | -0.0048 | -0.0659 | -0.0240 | 0.0943 | 0.0548 | -0.0260 |
| 3 | -0.0884 | 0.0707 | -0.0662 | -0.0618 | -0.1272 | -0.1402 | -0.2081 | -0.1273 | -0.0420 | -0.2135 |
| 12.11 | -0.0951 | 0.0893 | -0.0986 | -0.1442 | -0.1310 | -0.0633 | -0.0368 | 0.0431 | 0.0886 | 0.2449 |
| 12.19 | -0.1203 | 0.0473 | 0.1041 | -0.0321 | -0.1310 | -0.0340 | -0.0371 | 0.0496 | -0.0506 | 0.1723 |
| 12.21 | 0.0999 | 0.0570 | 0.0295 | -0.0021 | -0.0114 | 0.0783 | 0.0654 | -0.0067 | 0.0638 | -0.0556 |
| 12.23 | 0.0947 | 0.0564 | 0.1700 | -0.0026 | -0.0402 | 0.1052 | 0.0860 | 0.0509 | 0.0684 | -0.0888 |
| 12.25 | -0.0915 | 0.0901 | 0.0813 | 0.0054 | -0.0504 | 0.1797 | 0.1031 | -0.1306 | 0.0584 | -0.0950 |
| 12.26 | -0.0947 | 0.0626 | 0.0372 | 0.0219 | -0.1487 | 0.0174 | -0.0191 | -0.0637 | -0.0580 | 0.2111 |
| 12.32 | 0.0995 | 0.0550 | 0.0892 | -0.0376 | -0.0571 | 0.1366 | 0.1003 | 0.0128 | 0.1030 | -0.1106 |
| 12.34 | 0.1043 | 0.0752 | -0.1103 | -0.0742 | 0.1293 | 0.0055 | 0.0170 | 0.0093 | 0.0389 | 0.0007 |
| 12.39 | 0.0956 | 0.0627 | 0.1348 | 0.0230 | -0.0572 | 0.1506 | 0.1031 | -0.0170 | 0.0854 | -0.1243 |
| 12.4 | 0.0926 | 0.0417 | 0.0437 | -0.0090 | -0.0842 | -0.0502 | 0.0234 | 0.1116 | -0.0413 | 0.0913 |
| 12.43 | 0.0995 | 0.0549 | 0.0034 | 0.0207 | -0.0825 | -0.0104 | -0.0460 | 0.0013 | -0.0225 | 0.0793 |
| 12.48 | 0.1046 | 0.0797 | -0.0532 | -0.0338 | 0.1136 | 0.0079 | -0.0050 | 0.0274 | -0.0060 | 0.0114 |
| 12.49 | -0.0569 | 0.0890 | -0.1727 | -0.1080 | -0.1197 | 0.0246 | 0.0663 | -0.0355 | -0.0423 | 0.3616 |
| 12.5 | -0.0463 | 0.0760 | -0.2244 | -0.1651 | -0.1569 | 0.0841 | 0.4240 | 0.2208 | -0.7599 | -0.2146 |
| 12.52 | -0.1097 | 0.0946 | 0.0268 | -0.1062 | -0.1315 | -0.0276 | 0.0100 | 0.1065 | 0.0630 | 0.1952 |
| 12.61 | 0.0969 | 0.0465 | 0.0678 | 0.0231 | -0.0977 | 0.0119 | -0.0060 | 0.0286 | -0.0156 | 0.1220 |
| 12.64 | -0.1137 | 0.0848 | 0.1737 | -0.0406 | -0.0824 | 0.2221 | 0.1629 | -0.0504 | 0.1552 | -0.2321 |
| 12.71 | 0.0991 | 0.0562 | 0.0268 | 0.0112 | -0.0638 | 0.0404 | 0.0057 | 0.0059 | 0.0113 | 0.0310 |
| 12.73 | -0.1361 | 0.0261 | 0.0796 | -0.1640 | 0.0545 | -0.1326 | 0.0576 | 0.3097 | 0.1270 | -0.0259 |
| 12.75 | 0.0998 | 0.0466 | -0.1588 | -0.0832 | 0.0215 | 0.0043 | 0.0568 | 0.0858 | 0.0782 | -0.0201 |
| 2 | 0.0982 | 0.0527 | -0.0696 | -0.0340 | -0.0697 | -0.1032 | -0.1087 | -0.0024 | 0.0039 | -0.0956 |
| 5 | 0.0984 | 0.0506 | -0.0384 | -0.0022 | -0.0684 | -0.0908 | -0.1210 | -0.0151 | -0.0220 | -0.0920 |
| 6 | 0.0963 | 0.0418 | -0.0257 | -0.0186 | -0.0703 | -0.0992 | -0.1346 | 0.0022 | -0.0104 | -0.1047 |
| 7 | 0.0955 | 0.0451 | -0.0145 | -0.0180 | -0.0811 | -0.1148 | -0.1375 | 0.0151 | -0.0060 | -0.1235 |
| 77 | -0.0931 | -0.1386 | -0.1252 | -0.0571 | 0.0145 | 0.0811 | -0.0362 | -0.1017 | -0.0248 | -0.0158 |
| 78 | -0.1238 | -0.0720 | -0.1638 | 0.0322 | 0.0195 | 0.4440 | -0.3352 | 0.2293 | -0.0427 | 0.0434 |
| 8 | -0.0787 | 0.0841 | -0.0973 | -0.0711 | -0.1064 | -0.1614 | -0.2282 | -0.1061 | -0.0359 | -0.1926 |
| 80 | -0.0489 | -0.0369 | -0.2365 | -0.1191 | 0.0004 | 0.1041 | 0.1321 | -0.3217 | 0.1677 | 0.0868 |
| 84 | 0.0967 | -0.1219 | -0.0941 | 0.0074 | -0.0024 | 0.0387 | -0.0053 | -0.0568 | -0.0112 | 0.0162 |
| 9 | 0.0972 | 0.0364 | 0.0198 | 0.0227 | -0.0911 | -0.1008 | -0.1343 | -0.0160 | -0.0367 | -0.0941 |
| 90 | -0.1262 | -0.1996 | 0.0349 | 0.0185 | -0.0101 | 0.0398 | 0.0045 | -0.1049 | -0.0426 | -0.0022 |
| 91 | 0.0956 | -0.1448 | -0.0812 | -0.0142 | -0.0033 | 0.0327 | 0.0053 | -0.0423 | 0.0010 | 0.0068 |
| 92 | 0.1042 | -0.0824 | -0.1342 | -0.0402 | 0.0009 | 0.0251 | 0.0051 | -0.0498 | 0.0164 | 0.0008 |
| 93 | 0.0966 | -0.1752 | -0.0179 | 0.0353 | -0.0141 | 0.0311 | -0.0071 | -0.0732 | -0.0344 | 0.0108 |
| 99 | -0.1056 | 0.0171 | -0.1130 | 0.2429 | 0.0312 | -0.1139 | 0.0911 | 0.0722 | 0.0669 | -0.0267 |
| VT1 | 0.0789 | -0.2704 | 0.1225 | -0.1084 | 0.0347 | -0.0826 | 0.0150 | 0.1614 | 0.0223 | -0.0116 |
| VT10 | -0.1775 | -0.2448 | 0.1519 | -0.1765 | 0.0369 | -0.1127 | 0.0585 | 0.2010 | 0.0727 | -0.0387 |
| VT100 | -0.0721 | 0.0399 | -0.1848 | 0.1712 | 0.0129 | -0.0524 | 0.1303 | -0.0090 | 0.0675 | -0.0150 |
| VT101 | 0.0831 | -0.0258 | -0.0239 | 0.2536 | 0.0160 | -0.0910 | 0.0685 | 0.0789 | 0.0162 | -0.0019 |
| VT102 | -0.1186 | 0.0285 | -0.2044 | 0.2192 | 0.0378 | 0.4127 | -0.3635 | 0.4148 | 0.0271 | -0.0571 |
| VT103 | -0.1196 | -0.0155 | -0.0243 | 0.2836 | 0.0032 | -0.0846 | 0.0825 | 0.0306 | 0.0140 | -0.0332 |
| VT104 | -0.1069 | 0.0032 | -0.0511 | 0.2672 | 0.0247 | -0.0994 | 0.1088 | 0.0203 | 0.0192 | -0.0018 |
| VT107 | -0.1189 | -0.0014 | -0.0601 | 0.2788 | 0.0194 | -0.1461 | 0.0919 | 0.0683 | 0.0312 | -0.0236 |
| VT109 | 0.0855 | -0.0087 | -0.0732 | 0.2321 | 0.0174 | -0.0887 | 0.0722 | 0.0634 | 0.0251 | -0.0015 |
| VT110 | -0.1070 | 0.0118 | -0.0813 | 0.2609 | 0.0142 | -0.1074 | 0.0871 | 0.0035 | 0.0310 | -0.0692 |
| VT111 | -0.0805 | 0.0363 | -0.2038 | 0.1403 | 0.0469 | -0.1480 | 0.1725 | 0.0612 | 0.1494 | -0.0525 |
| VT12 | -0.1362 | -0.1669 | 0.0218 | -0.1072 | 0.0126 | -0.0322 | 0.0174 | 0.0491 | 0.0471 | 0.0150 |
| VT16 | -0.1197 | -0.2023 | 0.0564 | 0.0209 | -0.0011 | 0.0355 | -0.0311 | -0.1364 | -0.0898 | 0.0082 |
| VT20 | 0.1050 | -0.0804 | -0.1737 | -0.1107 | 0.0047 | 0.0145 | 0.0333 | 0.0170 | 0.0586 | -0.0141 |
| VT24 | 0.0965 | -0.2016 | 0.0082 | 0.0480 | -0.0105 | 0.0349 | -0.0223 | -0.0880 | -0.0667 | 0.0311 |
| VT30 | 0.0884 | -0.2236 | 0.0380 | -0.0604 | 0.0228 | -0.0380 | 0.0155 | 0.0728 | 0.0140 | -0.0174 |
| VT34 | 0.0972 | -0.1881 | 0.0000 | 0.0363 | -0.0039 | 0.0359 | -0.0217 | -0.0769 | -0.0591 | 0.0136 |
| VT4 | 0.0985 | -0.2109 | -0.0009 | 0.0348 | 0.0010 | 0.0202 | -0.0214 | -0.0739 | -0.0533 | 0.0217 |
| VT40 | -0.0632 | -0.0511 | -0.2478 | -0.1626 | 0.0190 | 0.0375 | 0.1307 | -0.0553 | 0.2130 | -0.0788 |
| VT44 | -0.1089 | -0.1915 | 0.0016 | 0.0148 | 0.0095 | 0.0406 | -0.0129 | -0.1268 | -0.0819 | 0.0225 |
| VT50 | 0.0944 | -0.1722 | -0.0391 | -0.0372 | -0.0024 | -0.0098 | -0.0000 | 0.0039 | 0.0023 | -0.0066 |
| VT52 | 0.0954 | -0.1632 | -0.0328 | 0.0170 | -0.0076 | 0.0183 | -0.0193 | -0.0417 | -0.0247 | 0.0027 |
| VT56 | 0.1009 | -0.2115 | 0.0170 | 0.0399 | -0.0106 | 0.0346 | -0.0231 | -0.0817 | -0.0699 | 0.0237 |
| VT70 | -0.1568 | -0.1839 | 0.0460 | -0.2322 | 0.0312 | -0.0991 | 0.0854 | 0.2522 | 0.1350 | -0.0330 |
| VT8 | -0.1216 | -0.1700 | 0.0304 | 0.0649 | -0.0075 | 0.0793 | -0.0112 | -0.1222 | -0.0783 | 0.0364 |

### Table S11. Autosomal PCA scores

An autosomal PCA run, providing per-sample scores across PCs, includes only the autosomal genome to examine population structure.

| snake | PC1 | PC2 | PC3 | PC4 | PC5 | PC6 | PC7 | PC8 | PC9 | PC10 | UMAP1 | UMAP2 |
|---|---|---|---|---|---|---|---|---|---|---|---|---|
| 09.06SFD | -0.0670 | 0.1124 | 0.0011 | 0.1247 | -0.1322 | -0.1772 | -0.1288 | 0.0967 | 0.0344 | -0.0222 | -2.0561 | 4.0222 |
| 1 | -0.0342 | 0.0007 | 0.0125 | 0.0263 | 0.2008 | 0.2077 | -0.2767 | 0.0050 | -0.0664 | 0.0147 | 0.8331 | 5.2438 |
| 10 | -0.0636 | -0.0586 | 0.0892 | 0.0470 | 0.1420 | 0.1037 | -0.2174 | -0.0065 | -0.0722 | -0.0338 | 0.5512 | 5.0179 |
| 10.16 | -0.0821 | 0.0110 | 0.0260 | -0.2494 | 0.0002 | 0.0024 | -0.0261 | -0.0308 | -0.1040 | 0.0499 | 1.5878 | 2.0843 |
| 10.17 | -0.0792 | -0.0224 | 0.0814 | 0.0990 | -0.0018 | -0.0136 | 0.1208 | -0.0041 | 0.0383 | 0.0215 | -0.8328 | 1.3887 |
| 10.21 | -0.1123 | -0.0747 | 0.1483 | -0.1626 | -0.0699 | -0.0107 | 0.0525 | 0.0486 | 0.0449 | 0.0436 | 0.6536 | 1.8675 |
| 10.34 | -0.1371 | -0.1175 | 0.1198 | -0.1175 | -0.0460 | -0.0495 | 0.0035 | -0.0322 | -0.1112 | -0.0082 | 0.9254 | 2.6539 |
| 10.4 | -0.0587 | 0.0492 | -0.0386 | 0.1600 | -0.0261 | 0.0756 | 0.1193 | -0.0948 | -0.0105 | 0.0305 | -2.3577 | 0.2496 |
| 11 | -0.1398 | -0.1221 | 0.1289 | 0.0364 | 0.0232 | -0.0105 | -0.0564 | 0.0243 | 0.0459 | -0.2268 | -0.1346 | 3.0163 |
| 11.04 | -0.0792 | -0.0115 | 0.0432 | -0.0185 | -0.0481 | 0.0916 | 0.0825 | -0.0169 | 0.0190 | 0.0102 | 0.0010 | 0.8660 |
| 11.07 | -0.0707 | 0.0951 | 0.0166 | 0.1112 | -0.1452 | -0.1939 | -0.0997 | 0.0738 | 0.0164 | -0.0276 | -1.9990 | 3.4730 |
| 11.1 | -0.1718 | -0.0901 | 0.1241 | -0.0587 | -0.0399 | -0.0688 | 0.0038 | -0.0107 | 0.0101 | 0.8645 | 0.3364 | 2.1288 |
| 11.24 | -0.0716 | -0.0192 | 0.0371 | 0.1006 | -0.0311 | -0.0084 | 0.0815 | -0.0018 | 0.0476 | -0.0065 | -0.6775 | 0.9728 |
| 11.28 | -0.0705 | 0.0270 | -0.0018 | -0.1396 | -0.1714 | 0.0959 | -0.0097 | -0.0407 | -0.0206 | -0.0493 | 1.5180 | 1.0121 |
| 11.29 | -0.0777 | 0.0569 | 0.0326 | -0.1354 | -0.0755 | -0.0263 | -0.0275 | 0.0730 | 0.0412 | 0.0187 | 0.7749 | 1.4632 |
| 11.3 | -0.0624 | 0.0639 | -0.0730 | -0.1635 | -0.0373 | 0.0204 | 0.0150 | -0.1088 | -0.0472 | -0.0438 | 2.2059 | 1.0850 |
| 11.31 | -0.0480 | 0.0379 | -0.0780 | 0.1058 | 0.0615 | 0.0303 | 0.0941 | -0.1339 | -0.1044 | 0.0316 | -2.3666 | 0.8218 |
| 11.32 | -0.0788 | 0.0541 | -0.0329 | -0.2795 | -0.0521 | 0.0630 | -0.0254 | -0.1441 | -0.1046 | -0.0633 | 2.1718 | 1.8573 |
| 11.35 | -0.0454 | 0.1587 | -0.0249 | -0.1981 | -0.0445 | 0.1204 | 0.0518 | 0.0166 | 0.2194 | -0.0137 | 1.1505 | 0.3390 |
| 11.36 | -0.0841 | 0.0124 | 0.0305 | -0.1594 | -0.1492 | 0.0565 | -0.0242 | -0.0142 | -0.0181 | -0.0734 | 1.2448 | 1.2478 |
| 11.38 | -0.0867 | 0.0139 | -0.0256 | -0.2006 | -0.2045 | 0.0920 | 0.0209 | -0.0827 | -0.0609 | -0.0605 | 1.9386 | 1.3009 |
| 11.4 | -0.0626 | 0.0716 | -0.0715 | -0.2304 | -0.0422 | 0.0593 | -0.0197 | -0.1525 | -0.1037 | -0.0566 | 2.2273 | 1.5990 |
| 11.41 | -0.0658 | 0.0788 | -0.0208 | -0.0974 | -0.1629 | 0.1206 | 0.0615 | -0.0362 | 0.1139 | -0.0433 | 1.3875 | 0.5583 |
| 11.43 | -0.0786 | 0.0126 | -0.0056 | -0.2175 | -0.0346 | -0.0003 | -0.0397 | -0.0374 | -0.1185 | 0.0169 | 1.7793 | 1.6575 |
| 11.44 | -0.0502 | 0.0924 | -0.0750 | 0.0983 | 0.0576 | -0.0545 | 0.0225 | -0.1280 | -0.1384 | 0.0050 | -2.6610 | 1.5238 |
| 11.45 | -0.0582 | 0.1738 | -0.0876 | 0.0547 | -0.0240 | -0.2517 | -0.2021 | 0.0282 | -0.1047 | -0.0134 | -2.9326 | 3.1723 |
| 11.46 | -0.0741 | 0.0264 | -0.0318 | -0.1363 | -0.1271 | 0.1097 | 0.0267 | -0.0420 | 0.0091 | 0.0026 | 1.9219 | 0.7525 |
| 11.52 | -0.0560 | 0.0374 | -0.0347 | 0.1122 | 0.0034 | 0.0806 | 0.1193 | -0.0243 | 0.0538 | 0.0080 | -1.2594 | -0.0991 |
| 11.53 | -0.0686 | -0.0010 | -0.0174 | 0.0929 | 0.0518 | -0.0644 | 0.1058 | -0.0917 | -0.1468 | 0.0226 | -2.1468 | 1.5966 |
| 11.54 | -0.0445 | 0.0938 | -0.0774 | 0.1469 | 0.0254 | 0.0848 | 0.1525 | -0.0444 | 0.0736 | 0.0444 | -2.0381 | -0.2928 |
| 11.57 | -0.0433 | 0.0559 | -0.0342 | 0.1629 | -0.0134 | 0.0941 | 0.0750 | -0.0596 | -0.0095 | 0.0229 | -1.8558 | 0.3575 |
| 11.58 | -0.0878 | -0.0594 | 0.0977 | 0.0610 | 0.1262 | -0.0467 | 0.1039 | -0.0630 | -0.0895 | 0.0188 | -1.3353 | 1.6478 |
| 11.64 | -0.0803 | 0.0100 | 0.0672 | 0.0067 | -0.0780 | -0.0961 | -0.0076 | 0.0937 | 0.0514 | 0.0045 | -0.9096 | 2.8597 |
| 11.68 | -0.0269 | 0.2304 | -0.0874 | 0.0561 | 0.0935 | 0.0770 | 0.1603 | 0.0026 | 0.1567 | 0.0535 | -1.7008 | -0.5626 |
| 11.73 | -0.0967 | -0.1191 | 0.1376 | 0.0116 | -0.0259 | -0.0549 | 0.0253 | 0.0319 | 0.0218 | 0.0066 | -0.0536 | 2.2238 |
| 11.74 | -0.0410 | 0.0342 | 0.0131 | -0.0648 | 0.1783 | -0.0049 | -0.0670 | 0.0199 | 0.0089 | 0.0343 | 0.0553 | 0.1069 |
| 12.11 | -0.0804 | -0.0567 | 0.0976 | 0.0479 | 0.1703 | -0.0442 | 0.1183 | -0.0809 | -0.0992 | 0.0161 | -1.6572 | 1.3617 |
| 12.19 | -0.0329 | 0.0811 | -0.0525 | 0.0598 | 0.1717 | 0.0374 | 0.1169 | -0.1237 | -0.0591 | 0.0277 | -2.5372 | 0.6876 |
| 12.21 | -0.0714 | 0.0594 | 0.0232 | 0.0527 | -0.1193 | -0.1065 | -0.0259 | 0.0713 | 0.0098 | -0.0002 | -1.6463 | 3.1508 |
| 12.23 | -0.0687 | 0.2074 | -0.0521 | 0.0481 | -0.0629 | -0.1538 | -0.0934 | 0.1029 | 0.1055 | 0.0108 | -2.5843 | 3.6841 |
| 12.25 | -0.0661 | 0.0725 | -0.0460 | 0.0403 | -0.0176 | -0.1598 | -0.0654 | -0.0419 | -0.1368 | -0.0219 | -2.5223 | 2.5505 |
| 12.26 | -0.0404 | 0.0302 | -0.0454 | 0.0992 | 0.0678 | 0.0013 | 0.1030 | -0.1407 | -0.1101 | 0.0152 | -2.6408 | 1.1404 |
| 12.32 | -0.0656 | 0.1477 | 0.0212 | 0.0957 | -0.0705 | -0.2188 | -0.1034 | 0.1281 | 0.0741 | 0.0010 | -2.7839 | 3.7905 |
| 12.34 | -0.1190 | -0.0961 | 0.1250 | -0.1212 | -0.0813 | -0.0235 | 0.0483 | 0.0548 | 0.0264 | 0.0285 | 0.7845 | 2.0799 |
| 12.39 | -0.0730 | 0.1772 | -0.0502 | 0.0901 | -0.0880 | -0.1888 | -0.1528 | 0.0812 | 0.0157 | -0.0181 | -2.5838 | 3.8190 |
| 12.4 | -0.0528 | 0.0633 | -0.0043 | 0.0818 | 0.0707 | 0.0589 | 0.2139 | 0.0048 | 0.1389 | -0.0388 | -1.3292 | -0.3912 |
| 12.43 | -0.0589 | 0.0005 | 0.0105 | 0.1298 | -0.0099 | 0.0900 | 0.1022 | -0.0422 | 0.0112 | -0.0015 | -1.1262 | 0.3203 |
| 12.48 | -0.1018 | -0.0406 | 0.0712 | -0.0909 | -0.1302 | 0.0367 | 0.0569 | -0.0082 | 0.1247 | -0.0589 | 0.4596 | 1.0350 |
| 12.49 | -0.1029 | -0.0874 | 0.0960 | 0.0676 | 0.0771 | -0.0773 | 0.1455 | -0.0850 | -0.0514 | -0.0732 | -1.0341 | 2.0892 |
| 12.5 | -0.1391 | -0.1141 | 0.1289 | 0.0389 | 0.0692 | -0.1189 | 0.1494 | -0.0383 | 0.0183 | -0.2993 | -0.4735 | 2.5414 |
| 12.52 | -0.0708 | 0.0279 | 0.0127 | 0.0305 | 0.1853 | -0.0463 | 0.1203 | -0.0583 | -0.0416 | 0.0101 | -1.6564 | 1.2622 |
| 12.61 | -0.0505 | 0.0801 | -0.0274 | 0.1449 | 0.0165 | 0.0713 | 0.1675 | -0.0964 | 0.0212 | 0.0493 | -2.3057 | -0.0459 |
| 12.64 | -0.0652 | 0.1655 | -0.0405 | 0.0374 | 0.0311 | -0.2764 | -0.1688 | 0.0506 | -0.0678 | 0.0017 | -2.4518 | 3.0074 |
| 12.71 | -0.0706 | 0.0475 | 0.0015 | 0.1116 | -0.0470 | -0.0088 | 0.0531 | -0.0214 | 0.0098 | -0.0093 | -1.3843 | 0.7681 |
| 12.73 | -0.0240 | 0.0744 | 0.0032 | -0.1652 | 0.2762 | -0.0839 | 0.0694 | 0.0831 | 0.1627 | -0.0818 | 0.3319 | -0.7903 |
| 12.75 | -0.1045 | -0.1328 | 0.1577 | -0.0129 | 0.0230 | -0.1141 | 0.0760 | 0.1068 | 0.1401 | -0.1848 | 0.1044 | 2.8166 |
| 2 | -0.0618 | -0.0760 | 0.0966 | 0.0846 | 0.0492 | 0.1549 | -0.1507 | 0.1053 | 0.0656 | 0.0113 | 1.7530 | 5.4872 |
| 3 | -0.0525 | -0.0745 | 0.0663 | 0.0525 | 0.1235 | 0.1292 | -0.2125 | 0.0077 | -0.0930 | -0.0215 | 0.7922 | 4.9556 |
| 5 | -0.0529 | -0.0579 | 0.0595 | 0.0911 | 0.0281 | 0.1905 | -0.1599 | 0.0549 | 0.0563 | -0.0124 | 1.4076 | 5.8522 |
| 6 | -0.0487 | -0.0439 | 0.0700 | 0.0821 | 0.0492 | 0.2016 | -0.1956 | 0.0807 | 0.0614 | 0.0014 | 1.0536 | 5.8361 |
| 7 | -0.0521 | -0.0309 | 0.0574 | 0.0956 | 0.0467 | 0.2007 | -0.1877 | 0.0610 | 0.0861 | -0.0017 | 1.5399 | 5.3583 |
| 77 | 0.1646 | -0.0860 | 0.1011 | -0.0155 | -0.0019 | -0.0593 | -0.0367 | -0.0822 | -0.0607 | 0.0067 | 1.4381 | -4.0284 |
| 78 | 0.1352 | -0.1296 | 0.0286 | 0.0094 | 0.0026 | -0.1529 | -0.1809 | -0.5038 | 0.2716 | -0.0060 | 0.7444 | -3.9382 |
| 8 | -0.0660 | -0.0864 | 0.0704 | 0.0466 | 0.1271 | 0.1236 | -0.1968 | 0.0045 | -0.0781 | -0.0018 | 1.2813 | 4.7846 |
| 80 | 0.1043 | -0.1399 | 0.1623 | 0.0086 | -0.0245 | -0.1061 | 0.0255 | -0.0067 | -0.1433 | 0.0472 | 1.5755 | -4.7858 |
| 84 | 0.1343 | -0.0832 | 0.0763 | 0.0414 | -0.0865 | -0.0192 | 0.0059 | -0.0318 | -0.0334 | -0.0005 | 1.6020 | -4.3442 |
| 9 | -0.0421 | -0.0041 | 0.0249 | 0.1107 | 0.0234 | 0.2300 | -0.2068 | 0.0510 | 0.0792 | 0.0043 | 1.4015 | 5.6338 |
| 90 | 0.1733 | 0.0272 | -0.0396 | -0.0385 | 0.0235 | -0.0348 | -0.0500 | -0.0659 | -0.1268 | -0.0199 | 1.4447 | -2.9847 |
| 91 | 0.1539 | -0.0622 | 0.0964 | 0.0362 | -0.0701 | -0.0242 | 0.0266 | 0.0184 | -0.0178 | 0.0032 | 2.2161 | -4.2632 |
| 92 | 0.1309 | -0.1031 | 0.1433 | 0.0484 | -0.0891 | -0.0384 | 0.0358 | 0.0389 | -0.0284 | -0.0047 | 2.0064 | -4.5533 |
| 93 | 0.1668 | 0.0015 | 0.0310 | 0.0686 | -0.1243 | 0.0288 | 0.0110 | 0.0020 | -0.0623 | -0.0047 | 2.8240 | -4.4751 |
| 99 | 0.0093 | -0.2055 | -0.2248 | -0.0289 | 0.0337 | -0.0436 | 0.0051 | 0.1043 | -0.0013 | 0.0162 | -2.8846 | -4.5760 |
| VT1 | 0.2074 | 0.1745 | 0.0576 | -0.1161 | 0.1225 | 0.0558 | 0.0628 | 0.1212 | 0.2126 | 0.0504 | 1.9940 | -2.3481 |
| VT10 | 0.1868 | 0.1552 | 0.0032 | -0.1896 | 0.2959 | -0.0542 | 0.0071 | 0.0870 | 0.0751 | 0.0338 | 1.6620 | -1.9176 |
| VT100 | -0.0155 | -0.2147 | -0.1179 | -0.0026 | -0.0090 | -0.0749 | 0.0259 | 0.0579 | -0.0471 | -0.0165 | -2.1491 | -4.7509 |
| VT101 | 0.0295 | -0.1325 | -0.2708 | 0.0131 | -0.0603 | 0.0213 | 0.0543 | 0.1774 | 0.1016 | 0.0095 | -2.3134 | -3.9710 |
| VT102 | 0.0090 | -0.2206 | -0.1435 | 0.0090 | 0.0025 | -0.1762 | -0.1824 | -0.4241 | 0.5195 | 0.0364 | -0.7098 | -3.9631 |
| VT103 | 0.0326 | -0.1407 | -0.2805 | -0.0195 | 0.0395 | -0.0156 | -0.0018 | 0.0696 | -0.0306 | 0.0133 | -2.1636 | -4.6253 |
| VT104 | 0.0184 | -0.1582 | -0.2498 | -0.0309 | -0.0012 | -0.0214 | 0.0288 | 0.0895 | -0.0226 | 0.0142 | -2.8435 | -4.5144 |
| VT107 | 0.0245 | -0.1628 | -0.2703 | -0.0237 | 0.0545 | -0.0129 | 0.0131 | 0.1092 | -0.0116 | 0.0073 | -2.9809 | -4.0896 |
| VT109 | 0.0176 | -0.1828 | -0.2318 | 0.0158 | -0.0689 | 0.0172 | 0.0651 | 0.1449 | 0.0552 | 0.0225 | -1.9534 | -4.3128 |
| VT110 | 0.0151 | -0.1668 | -0.2301 | -0.0098 | 0.0308 | -0.0216 | -0.0282 | 0.0938 | -0.0479 | 0.0137 | -2.6930 | -4.0098 |
| VT111 | -0.0066 | -0.2235 | -0.1186 | -0.0333 | 0.0440 | -0.0698 | 0.0768 | 0.1686 | -0.0101 | -0.0033 | -2.6782 | -5.0000 |
| VT12 | 0.1460 | 0.0380 | 0.0282 | -0.0811 | 0.1374 | -0.0549 | -0.0144 | 0.0061 | -0.0680 | 0.0160 | 1.2967 | -2.6995 |
| VT16 | 0.1673 | 0.0449 | -0.0354 | -0.0168 | 0.0099 | 0.0133 | -0.0510 | -0.0715 | -0.1105 | 0.0034 | 1.1734 | -3.2031 |
| VT20 | 0.1309 | -0.1346 | 0.2307 | 0.0137 | -0.0450 | -0.0796 | 0.1040 | 0.1554 | 0.0992 | 0.0258 | 2.1123 | -4.9371 |
| VT24 | 0.1825 | 0.0337 | 0.0184 | 0.0694 | -0.1440 | 0.0648 | 0.0123 | -0.0414 | -0.0559 | 0.0028 | 2.8784 | -3.6918 |
| VT30 | 0.1906 | 0.0941 | 0.0849 | -0.0319 | 0.0029 | 0.0271 | 0.0459 | 0.1004 | 0.1294 | 0.0157 | 2.4220 | -2.9457 |
| VT34 | 0.1762 | 0.0306 | 0.0285 | 0.0676 | -0.1610 | 0.0487 | -0.0039 | -0.0305 | -0.0368 | -0.0005 | 3.0028 | -4.0691 |
| VT4 | 0.1932 | 0.0164 | 0.0446 | 0.0638 | -0.1430 | 0.0748 | 0.0349 | 0.0219 | 0.0028 | 0.0163 | 3.2969 | -4.1067 |
| VT40 | 0.0877 | -0.1470 | 0.1779 | -0.0336 | 0.0478 | -0.1490 | 0.0378 | -0.0009 | -0.0900 | -0.0352 | 1.2773 | -4.8180 |
| VT44 | 0.1669 | 0.0021 | -0.0165 | -0.0327 | 0.0051 | -0.0089 | -0.0344 | -0.0810 | -0.1026 | -0.0077 | 1.4644 | -3.5585 |
| VT50 | 0.1701 | 0.0012 | 0.1003 | 0.0071 | -0.0522 | 0.0143 | 0.0422 | 0.0669 | 0.0516 | 0.0050 | 2.4195 | -3.5911 |
| VT52 | 0.1562 | -0.0116 | 0.0514 | 0.0472 | -0.1009 | 0.0455 | -0.0134 | -0.0076 | -0.0328 | 0.0146 | 2.3085 | -3.8359 |
| VT56 | 0.1915 | 0.0417 | 0.0327 | 0.0829 | -0.1733 | 0.0791 | 0.0347 | -0.0121 | -0.0212 | 0.0089 | 3.0232 | -3.7192 |
| VT70 | 0.1538 | 0.0678 | 0.1054 | -0.1805 | 0.2905 | -0.1093 | 0.0380 | 0.0743 | 0.0697 | 0.0270 | 1.1544 | -2.1620 |
| VT8 | 0.1499 | -0.0006 | -0.0607 | -0.0143 | -0.0134 | -0.0187 | -0.0447 | -0.1336 | -0.1295 | -0.0136 | 1.8931 | -3.2088 |

**Table S12. PCA variance partitioning**

Variance explained and ANOVA summaries attributing PC variation to sequencing coverage and Town (population), with F- and P-values and cumulative $R^2$ per PC.

| PC | Predictor | PercentVar | F value | P value |
|---|---|---|---|---|
| PC1 | Coverage (1st) | 0.1383759 | 119.2389 | 4.12532e-18 |
| PC1 | Population (2nd) | 0.7583402 | 108.9106 | 8.297245e-39 |
| PC1 | Sum (=R^2) | 0.8967161 | 110.3861 | 4.466119e-41 |
| PC2 | Coverage (1st) | 0.2305026 | 77.50853 | 9.56489e-14 |
| PC2 | Population (2nd) | 0.5048202 | 28.2917 | 1.103982e-18 |
| PC2 | Sum (=R^2) | 0.7353229 | 35.32268 | 4.2342e-23 |
| Overall | Coverage differences among Populations (one-way ANOVA) | NA | 5.534508 | 0.00017029 |
| NA | NA | | | |
| UMAP | Predictor | PropVar | F_value | P_value |
| UMAP1 | Coverage (1st) | 0.00038784 | 0.07716487 | 0.7818212 |
| UMAP1 | Population (2nd) | 0.5522925 | 18.31428 | 9.650744e-14 |
| UMAP1 | Sum (=R^2) | 0.5526803 | 15.70898 | 2.945518e-13 |
| UMAP2 | Coverage (1st) | 0.05432331 | 38.31369 | 1.808133e-08 |
| UMAP2 | Population (2nd) | 0.8194875 | 96.3294 | 9.504129e-37 |
| UMAP2 | Sum (=R^2) | 0.8738108 | 88.04144 | 3.115823e-37 |
| Overall | Coverage differences among Populations (one-way ANOVA) | NA | 4.608565 | 0.0003993 |

**Table S13. GWAS significant loci near ALDH4A1**

Top association results for color-morph GWAS on chr16, including genotype counts (cases/controls), genotype-specific penetrance, allele frequencies, positions, effect sizes (β), SE, and p-values (–log10 P).

| SNP | chr16:7387613 | chr16:7387777 | chr16:7411568 | chr16:7411601 | chr16:7411644 | chr16:7414099 |
|---|---|---|---|---|---|---|
| Cases_AA | 15 | 14 | 16 | 12 | 18 | 17 |
| Cases_AB | 8 | 8 | 6 | 9 | 7 | 6 |
| Cases_BB | 2 | 3 | 3 | 4 | 0 | 2 |
| Controls_AA | 1 | 1 | 2 | 0 | 3 | 2 |
| Controls_AB | 15 | 15 | 12 | 10 | 11 | 12 |
| Controls_BB | 8 | 8 | 10 | 14 | 10 | 10 |
| Penetrance_AA | 0.938 | 0.933 | 0.889 | 1 | 0.857 | 0.895 |
| Penetrance_AB | 0.348 | 0.348 | 0.333 | 0.474 | 0.389 | 0.333 |
| Penetrance_BB | 0.2 | 0.273 | 0.231 | 0.222 | 0 | 0.167 |
| Major allele | A | T | T | A | A | A |
| Minor allele | G | C | C | G | G | G |
| Frequency | 0.439301 | 0.461734 | 0.438117 | 0.53963 | 0.37421 | 0.419927 |
| P | 4.6169e-10 | 3.2831e-10 | 1.5372e-08 | 1.4044e-08 | 4.8253e-09 | 3.3705e-09 |
| beta | -20.58566 | -21.51563 | -16.47237 | -11.65007 | -31.94682 | -15.79936 |
| SE | 46.16613 | 40.70615 | 20.91315 | 7.259528 | 108.96 | 17.05073 |
| high_WT/HE/HO | 15/14/9 | 15/14/9 | 15/12/8 | 2014-12-10 | 19/14/4 | 16/12/8 |
| emIter | 5 | 4 | 10 | 6 | 12 | 3 |
| minusLog10P | 9.33565 | 9.483716 | 7.81327 | 7.852509 | 8.316476 | 8.472306 |

**Table S14. Kraken2 classification summary**

Per-sample metagenomic classification totals by high-level taxonomic “Type” (e.g., bacteria), including percent classified/unclassified and read counts.

| snake | Type | Percent classified root | Percent unclassified | Classified reads | Unclassified reads | Total reads |
|---|---|---|---|---|---|---|
| 09.06SFD | bacteria | 38.2800 | 0.000 | 899,325 | 1 | 899,326 |
| 10.16 | bacteria | 46.5000 | 0.000 | 1,262,201 | 1 | 1,262,202 |
| 10.17 | bacteria | 60.0600 | 0.000 | 1,189,171 | 1 | 1,189,172 |
| 10.21 | bacteria | 71.0900 | 0.000 | 2,112,636 | 2 | 2,112,638 |
| 10.34 | bacteria | 62.6700 | 0.000 | 1,382,967 | 9 | 1,382,976 |
| 10.4 | bacteria | 30.3200 | 0.000 | 530,560 | 5 | 530,565 |
| 10 | bacteria | 44.2900 | 0.000 | 838,453 | 1 | 838,454 |
| 11.04 | bacteria | 44.7400 | 0.000 | 1,024,391 | 1 | 1,024,392 |
| 11.07 | bacteria | 44.8500 | 0.000 | 1,116,622 | 1 | 1,116,623 |
| 11.1 | bacteria | 81.7500 | 0.000 | 1,189,714 | 8 | 1,189,722 |
| 11.24 | bacteria | 45.8900 | 0.000 | 1,035,852 | 3 | 1,035,855 |
| 11.28 | bacteria | 35.0500 | 0.000 | 802,918 | 1 | 802,919 |
| 11.29 | bacteria | 47.4300 | 0.000 | 1,450,925 | 1 | 1,450,926 |
| 11.3 | bacteria | 45.4000 | 0.000 | 1,256,843 | 20 | 1,256,863 |
| 11.31 | bacteria | 45.6700 | 0.000 | 1,527,602 | 2 | 1,527,604 |
| 11.32 | bacteria | 42.3000 | 0.000 | 1,097,813 | 1 | 1,097,814 |
| 11.35 | bacteria | 36.8700 | 0.000 | 916,338 | 1 | 916,339 |
| 11.36 | bacteria | 46.7200 | 0.000 | 1,364,057 | 1 | 1,364,058 |
| 11.38 | bacteria | 30.9000 | 0.000 | 524,548 | 2 | 524,550 |
| 11.41 | bacteria | 32.7600 | 0.000 | 816,712 | 1 | 816,713 |
| 11.43 | bacteria | 47.4700 | 0.000 | 1,198,985 | 1 | 1,198,986 |
| 11.44 | bacteria | 35.6300 | 0.000 | 899,291 | 4 | 899,295 |
| 11.45 | bacteria | 26.1800 | 0.000 | 451,974 | 1 | 451,975 |
| 11.46 | bacteria | 31.3800 | 0.000 | 761,918 | 1 | 761,919 |
| 11.4 | bacteria | 29.5500 | 0.000 | 583,362 | 1 | 583,363 |
| 11.52 | bacteria | 36.3600 | 0.000 | 956,970 | 1 | 956,971 |
| 11.53 | bacteria | 41.9800 | 0.000 | 845,122 | 1 | 845,123 |
| 11.54 | bacteria | 39.7500 | 0.000 | 900,773 | 10 | 900,783 |
| 11.57 | bacteria | 29.9100 | 0.000 | 632,647 | 1 | 632,648 |
| 11.58 | bacteria | 76.3900 | 0.000 | 3,094,741 | 1 | 3,094,742 |
| 11.64 | bacteria | 58.6100 | 0.000 | 1,050,598 | 1 | 1,050,599 |
| 11.68 | bacteria | 32.6700 | 0.000 | 2,298,551 | 2 | 2,298,553 |
| 11.73 | bacteria | 78.3800 | 0.000 | 2,116,154 | 2 | 2,116,156 |
| 11.74 | bacteria | 67.3700 | 0.000 | 2,647,916 | 14 | 2,647,930 |
| 11 | bacteria | 63.0800 | 0.000 | 464,616 | 4 | 464,620 |
| 12.11 | bacteria | 58.0200 | 0.000 | 824,125 | 4 | 824,129 |
| 12.19 | bacteria | 39.9800 | 0.000 | 836,608 | 4 | 836,612 |
| 12.21 | bacteria | 45.2400 | 0.000 | 1,231,702 | 1 | 1,231,703 |
| 12.23 | bacteria | 38.2500 | 0.000 | 989,467 | 5 | 989,472 |
| 12.25 | bacteria | 38.7700 | 0.000 | 788,177 | 1 | 788,178 |
| 12.26 | bacteria | 39.3000 | 0.000 | 860,066 | 1 | 860,067 |
| 12.32 | bacteria | 51.6900 | 0.000 | 1,561,631 | 1 | 1,561,632 |
| 12.34 | bacteria | 69.7800 | 0.000 | 1,734,912 | 1 | 1,734,913 |
| 12.39 | bacteria | 38.9000 | 0.000 | 1,003,619 | 28 | 1,003,647 |
| 12.4 | bacteria | 51.5100 | 0.000 | 1,594,616 | 2 | 1,594,618 |
| 12.43 | bacteria | 35.5800 | 0.000 | 651,965 | 1 | 651,966 |
| 12.48 | bacteria | 57.3800 | 0.000 | 1,592,698 | 2 | 1,592,700 |
| 12.49 | bacteria | 58.4000 | 0.000 | 826,865 | 1 | 826,866 |
| 12.5 | bacteria | 70.6000 | 0.000 | 777,116 | 2 | 777,118 |
| 12.51 | bacteria | 71.0400 | 0.000 | 141,436 | 1 | 141,437 |
| 12.52 | bacteria | 49.5700 | 0.000 | 1,237,090 | 21 | 1,237,111 |
| 12.61 | bacteria | 39.4700 | 0.000 | 799,837 | 2 | 799,839 |
| 12.64 | bacteria | 50.8600 | 0.000 | 1,357,217 | 2 | 1,357,219 |
| 12.71 | bacteria | 43.1500 | 0.000 | 1,066,129 | 22 | 1,066,151 |
| 12.73 | bacteria | 69.7300 | 0.000 | 8,478,776 | 1 | 8,478,777 |
| 12.74 | bacteria | 91.1800 | 0.000 | 3,218,617 | 1 | 3,218,618 |
| 12.75 | bacteria | 76.6700 | 0.000 | 2,443,810 | 10 | 2,443,820 |
| 162019 | bacteria | 95.4000 | 0.000 | 15,939,070 | 6 | 15,939,076 |
| 186035 | bacteria | 75.5400 | 0.000 | 352,297 | 2 | 352,299 |
| 186036 | bacteria | 77.7200 | 0.000 | 356,036 | 1 | 356,037 |
| 186037 | bacteria | 81.8600 | 0.000 | 335,789 | 2 | 335,791 |
| 19192 | bacteria | 96.0800 | 0.000 | 25,153,008 | 3 | 25,153,011 |
| 1 | bacteria | 35.8700 | 0.000 | 889,529 | 1 | 889,530 |
| 27544 | bacteria | 96.0600 | 0.000 | 17,236,172 | 1 | 17,236,173 |
| 2 | bacteria | 47.5000 | 0.000 | 1,116,598 | 1 | 1,116,599 |
| 3 | bacteria | 42.0300 | 0.000 | 850,503 | 18 | 850,521 |
| 5 | bacteria | 44.1000 | 0.000 | 1,137,735 | 1 | 1,137,736 |
| 600.1 | bacteria | 98.7100 | 0.000 | 13,037,917 | 1 | 13,037,918 |

**Table S14. Kraken2 classification summary**

Per-sample metagenomic classification totals by high-level taxonomic “Type” (e.g., bacteria), including percent classified/unclassified and read counts.

| snake | Type | Percent classified root | Percent unclassified | Classified reads | Unclassified reads | Total reads |
|---|---|---|---|---|---|---|
| 6439 | bacteria | 96.4900 | 0.000 | 6,443,788 | 6 | 6,443,794 |
| 6462 | bacteria | 96.0500 | 0.000 | 22,479,054 | 2 | 22,479,056 |
| 6 | bacteria | 40.7700 | 0.000 | 804,500 | 1 | 804,501 |
| 77 | bacteria | 37.4100 | 0.000 | 613,164 | 1 | 613,165 |
| 78 | bacteria | 39.6600 | 0.000 | 685,802 | 1 | 685,803 |
| 7 | bacteria | 45.4100 | 0.000 | 1,397,089 | 1 | 1,397,090 |
| 80 | bacteria | 62.4200 | 0.000 | 488,939 | 1 | 488,940 |
| 81 | bacteria | 81.6800 | 0.000 | 14,702,981 | 2 | 14,702,983 |
| 82303 | bacteria | 95.9200 | 0.000 | 4,872,199 | 1 | 4,872,200 |
| 84 | bacteria | 41.0400 | 0.000 | 718,689 | 3 | 718,692 |
| 8 | bacteria | 51.8800 | 0.000 | 1,442,724 | 1 | 1,442,725 |
| 90 | bacteria | 26.6400 | 0.000 | 513,090 | 1 | 513,091 |
| 91 | bacteria | 43.1100 | 0.000 | 789,487 | 1 | 789,488 |
| 92 | bacteria | 52.6300 | 0.000 | 728,225 | 1 | 728,226 |
| 93 | bacteria | 22.8500 | 0.000 | 413,670 | 1 | 413,671 |
| 99 | bacteria | 42.5200 | 0.000 | 910,158 | 21 | 910,179 |
| 9 | bacteria | 36.1600 | 0.000 | 995,455 | 43 | 995,498 |
| R-4404 | bacteria | 96.0000 | 0.000 | 9,915,220 | 1 | 9,915,221 |
| VT100 | bacteria | 73.9900 | 0.000 | 3,212,514 | 4 | 3,212,518 |
| VT101 | bacteria | 30.7100 | 0.000 | 741,119 | 1 | 741,120 |
| VT102 | bacteria | 26.4100 | 0.000 | 287,480 | 1 | 287,481 |
| VT103 | bacteria | 29.4600 | 0.000 | 690,800 | 26 | 690,826 |
| VT104 | bacteria | 34.1400 | 0.000 | 800,327 | 1 | 800,328 |
| VT105 | bacteria | 56.2800 | 0.000 | 13,209,370 | 1 | 13,209,371 |
| VT107 | bacteria | 33.9700 | 0.000 | 836,121 | 1 | 836,122 |
| VT109 | bacteria | 31.1100 | 0.000 | 569,089 | 16 | 569,105 |
| VT10 | bacteria | 46.4900 | 0.000 | 5,835,048 | 1 | 5,835,049 |
| VT110 | bacteria | 31.7800 | 0.000 | 598,349 | 23 | 598,372 |
| VT111 | bacteria | 54.7500 | 0.000 | 1,547,227 | 1 | 1,547,228 |
| VT12 | bacteria | 49.6400 | 0.000 | 1,608,232 | 1 | 1,608,233 |
| VT16 | bacteria | 32.1900 | 0.000 | 916,591 | 1 | 916,592 |
| VT1 | bacteria | 57.0000 | 0.000 | 12,159,051 | 2 | 12,159,053 |
| VT20 | bacteria | 72.4900 | 0.000 | 1,951,946 | 2 | 1,951,948 |
| VT24 | bacteria | 22.0800 | 0.000 | 365,667 | 1 | 365,668 |
| VT30 | bacteria | 46.0800 | 0.000 | 2,695,237 | 1 | 2,695,238 |
| VT34 | bacteria | 29.9100 | 0.000 | 683,030 | 1 | 683,031 |
| VT40 | bacteria | 61.6600 | 0.000 | 1,433,230 | 1 | 1,433,231 |
| VT44 | bacteria | 32.9800 | 0.000 | 782,895 | 1 | 782,896 |
| VT4 | bacteria | 30.0400 | 0.000 | 687,230 | 2 | 687,232 |
| VT50 | bacteria | 43.9300 | 0.000 | 1,673,318 | 1 | 1,673,319 |
| VT52 | bacteria | 42.1300 | 0.000 | 1,064,280 | 38 | 1,064,318 |
| VT56 | bacteria | 25.8000 | 0.000 | 473,390 | 1 | 473,391 |
| VT70 | bacteria | 53.7100 | 0.000 | 2,789,192 | 2 | 2,789,194 |
| VT8 | bacteria | 25.0000 | 0.000 | 573,660 | 18 | 573,678 |
| 09.06SFD | fungus | 3.5000 | 96.500 | 82,216 | 2,266,957 | 2,349,173 |
| 10.16 | fungus | 3.4200 | 96.580 | 92,764 | 2,621,443 | 2,714,207 |
| 10.17 | fungus | 3.6900 | 96.310 | 73,155 | 1,906,726 | 1,979,881 |
| 10.21 | fungus | 3.0200 | 96.980 | 89,870 | 2,881,777 | 2,971,647 |
| 10.34 | fungus | 3.5900 | 96.410 | 79,326 | 2,127,479 | 2,206,805 |
| 10.4 | fungus | 3.8600 | 96.140 | 67,610 | 1,682,277 | 1,749,887 |
| 10 | fungus | 4.5000 | 95.500 | 85,236 | 1,807,678 | 1,892,914 |
| 11.04 | fungus | 3.4000 | 96.600 | 77,962 | 2,211,808 | 2,289,770 |
| 11.07 | fungus | 2.9600 | 97.040 | 73,589 | 2,416,012 | 2,489,601 |
| 11.1 | fungus | 3.7400 | 96.260 | 54,493 | 1,400,856 | 1,455,349 |
| 11.24 | fungus | 3.6000 | 96.400 | 81,317 | 2,175,827 | 2,257,144 |
| 11.28 | fungus | 3.6700 | 96.330 | 84,155 | 2,206,398 | 2,290,553 |
| 11.29 | fungus | 3.0400 | 96.960 | 92,949 | 2,966,315 | 3,059,264 |
| 11.3 | fungus | 3.0500 | 96.950 | 84,497 | 2,683,989 | 2,768,486 |
| 11.31 | fungus | 3.2800 | 96.720 | 109,825 | 3,234,889 | 3,344,714 |
| 11.32 | fungus | 3.4600 | 96.540 | 89,902 | 2,505,240 | 2,595,142 |
| 11.35 | fungus | 3.4200 | 96.580 | 84,908 | 2,400,643 | 2,485,551 |
| 11.36 | fungus | 3.4400 | 96.560 | 100,411 | 2,818,951 | 2,919,362 |
| 11.38 | fungus | 3.9600 | 96.040 | 67,220 | 1,630,245 | 1,697,465 |
| 11.41 | fungus | 3.3100 | 96.690 | 82,543 | 2,410,129 | 2,492,672 |
| 11.43 | fungus | 5.3400 | 94.660 | 134,994 | 2,390,876 | 2,525,870 |
| 11.44 | fungus | 3.4900 | 96.510 | 88,041 | 2,435,788 | 2,523,829 |
| 11.45 | fungus | 3.9700 | 96.030 | 68,470 | 1,657,872 | 1,726,342 |
| 11.46 | fungus | 3.2400 | 96.760 | 78,659 | 2,349,379 | 2,428,038 |
| 11.4 | fungus | 4.0600 | 95.940 | 80,251 | 1,894,219 | 1,974,470 |
| 11.52 | fungus | 3.4500 | 96.550 | 90,711 | 2,541,552 | 2,632,263 |

**Table S14. Kraken2 classification summary**

Per-sample metagenomic classification totals by high-level taxonomic “Type” (e.g., bacteria), including percent classified/unclassified and read counts.

| snake | Type | Percent classified root | Percent unclassified | Classified reads | Unclassified reads | Total reads |
|---|---|---|---|---|---|---|
| 11.53 | fungus | 4.1400 | 95.860 | 83,425 | 1,929,606 | 2,013,031 |
| 11.54 | fungus | 3.3500 | 96.650 | 75,906 | 2,189,921 | 2,265,827 |
| 11.57 | fungus | 3.1400 | 96.860 | 66,393 | 2,048,815 | 2,115,208 |
| 11.58 | fungus | 2.5600 | 97.440 | 103,552 | 3,947,687 | 4,051,239 |
| 11.64 | fungus | 3.8100 | 96.190 | 68,330 | 1,724,212 | 1,792,542 |
| 11.68 | fungus | 2.6500 | 97.350 | 186,747 | 6,848,054 | 7,034,801 |
| 11.73 | fungus | 14.7500 | 85.250 | 398,335 | 2,301,597 | 2,699,932 |
| 11.74 | fungus | 2.8200 | 97.180 | 110,758 | 3,819,849 | 3,930,607 |
| 11 | fungus | 7.2400 | 92.760 | 53,308 | 683,216 | 736,524 |
| 12.11 | fungus | 4.8100 | 95.190 | 68,333 | 1,352,188 | 1,420,521 |
| 12.19 | fungus | 4.1300 | 95.870 | 86,431 | 2,006,266 | 2,092,697 |
| 12.21 | fungus | 3.1800 | 96.820 | 86,526 | 2,636,206 | 2,722,732 |
| 12.23 | fungus | 3.2900 | 96.710 | 85,127 | 2,501,416 | 2,586,543 |
| 12.25 | fungus | 3.6200 | 96.380 | 73,599 | 1,959,521 | 2,033,120 |
| 12.26 | fungus | 3.6800 | 96.320 | 80,440 | 2,107,978 | 2,188,418 |
| 12.32 | fungus | 3.8500 | 96.150 | 116,356 | 2,904,995 | 3,021,351 |
| 12.34 | fungus | 3.7900 | 96.210 | 94,295 | 2,391,969 | 2,486,264 |
| 12.39 | fungus | 3.2300 | 96.770 | 83,430 | 2,496,374 | 2,579,804 |
| 12.4 | fungus | 3.0900 | 96.910 | 95,638 | 2,999,900 | 3,095,538 |
| 12.43 | fungus | 3.7800 | 96.220 | 69,330 | 1,763,219 | 1,832,549 |
| 12.48 | fungus | 4.5500 | 95.450 | 126,282 | 2,649,332 | 2,775,614 |
| 12.49 | fungus | 5.8200 | 94.180 | 82,337 | 1,333,438 | 1,415,775 |
| 12.5 | fungus | 5.1500 | 94.850 | 56,739 | 1,043,949 | 1,100,688 |
| 12.51 | fungus | 15.8600 | 84.140 | 31,577 | 167,504 | 199,081 |
| 12.52 | fungus | 3.5900 | 96.410 | 89,511 | 2,405,920 | 2,495,431 |
| 12.61 | fungus | 5.1100 | 94.890 | 103,524 | 1,922,862 | 2,026,386 |
| 12.64 | fungus | 3.6100 | 96.390 | 96,340 | 2,572,338 | 2,668,678 |
| 12.71 | fungus | 4.0800 | 95.920 | 100,883 | 2,369,652 | 2,470,535 |
| 12.73 | fungus | 30.2900 | 69.710 | 3,683,011 | 8,476,464 | 12,159,475 |
| 12.74 | fungus | 3.2400 | 96.760 | 114,392 | 3,415,587 | 3,529,979 |
| 12.75 | fungus | 5.3600 | 94.640 | 170,735 | 3,016,902 | 3,187,637 |
| 162019 | fungus | 1.1800 | 98.820 | 197,747 | 16,510,054 | 16,707,801 |
| 186035 | fungus | 15.5500 | 84.450 | 72,515 | 393,861 | 466,376 |
| 186036 | fungus | 10.2400 | 89.760 | 46,895 | 411,192 | 458,087 |
| 186037 | fungus | 8.3700 | 91.630 | 34,324 | 375,888 | 410,212 |
| 19192 | fungus | 1.4500 | 98.550 | 380,302 | 25,798,252 | 26,178,554 |
| 1 | fungus | 3.8000 | 96.200 | 94,198 | 2,385,909 | 2,480,107 |
| 27544 | fungus | 1.4300 | 98.570 | 257,201 | 17,685,422 | 17,942,623 |
| 2 | fungus | 3.9900 | 96.010 | 93,723 | 2,256,833 | 2,350,556 |
| 3 | fungus | 4.3900 | 95.610 | 88,863 | 1,934,927 | 2,023,790 |
| 5 | fungus | 3.5900 | 96.410 | 92,716 | 2,487,131 | 2,579,847 |
| 600.1 | fungus | 0.5900 | 99.410 | 78,392 | 13,129,762 | 13,208,154 |
| 6439 | fungus | 1.3900 | 98.610 | 93,057 | 6,585,386 | 6,678,443 |
| 6462 | fungus | 1.3800 | 98.620 | 324,050 | 23,080,264 | 23,404,314 |
| 6 | fungus | 3.3600 | 96.640 | 66,256 | 1,906,842 | 1,973,098 |
| 77 | fungus | 4.3100 | 95.690 | 70,676 | 1,568,283 | 1,638,959 |
| 78 | fungus | 2.5300 | 97.470 | 43,705 | 1,685,469 | 1,729,174 |
| 7 | fungus | 3.4800 | 96.520 | 107,066 | 2,969,434 | 3,076,500 |
| 80 | fungus | 5.3100 | 94.690 | 41,607 | 741,690 | 783,297 |
| 81 | fungus | 1.1400 | 98.860 | 204,675 | 17,795,601 | 18,000,276 |
| 82303 | fungus | 4.2800 | 95.720 | 217,642 | 4,861,911 | 5,079,553 |
| 84 | fungus | 5.8900 | 94.110 | 103,174 | 1,647,858 | 1,751,032 |
| 8 | fungus | 3.6300 | 96.370 | 101,010 | 2,679,916 | 2,780,926 |
| 90 | fungus | 3.7300 | 96.270 | 71,935 | 1,854,181 | 1,926,116 |
| 91 | fungus | 6.0100 | 93.990 | 110,093 | 1,721,399 | 1,831,492 |
| 92 | fungus | 4.9900 | 95.010 | 69,043 | 1,314,572 | 1,383,615 |
| 93 | fungus | 3.4400 | 96.560 | 62,306 | 1,748,181 | 1,810,487 |
| 99 | fungus | 4.9100 | 95.090 | 105,173 | 2,035,404 | 2,140,577 |
| 9 | fungus | 3.4500 | 96.550 | 95,037 | 2,657,560 | 2,752,597 |
| R-4404 | fungus | 1.3600 | 98.640 | 140,697 | 10,187,161 | 10,327,858 |
| VT100 | fungus | 4.3000 | 95.700 | 186,634 | 4,155,475 | 4,342,109 |
| VT101 | fungus | 4.8200 | 95.180 | 116,420 | 2,296,838 | 2,413,258 |
| VT102 | fungus | 2.8600 | 97.140 | 31,143 | 1,057,362 | 1,088,505 |
| VT103 | fungus | 3.5500 | 96.450 | 83,149 | 2,261,513 | 2,344,662 |
| VT104 | fungus | 3.9100 | 96.090 | 91,742 | 2,252,312 | 2,344,054 |
| VT105 | fungus | 0.7200 | 99.280 | 169,790 | 23,301,205 | 23,470,995 |
| VT107 | fungus | 4.0600 | 95.940 | 100,008 | 2,361,040 | 2,461,048 |
| VT109 | fungus | 3.9800 | 96.020 | 72,795 | 1,756,634 | 1,829,429 |
| VT10 | fungus | 2.7700 | 97.230 | 347,265 | 12,203,889 | 12,551,154 |
| VT110 | fungus | 4.6300 | 95.370 | 87,213 | 1,795,459 | 1,882,672 |

**Table S14. Kraken2 classification summary**

Per-sample metagenomic classification totals by high-level taxonomic “Type” (e.g., bacteria), including percent classified/unclassified and read counts.

| snake | Type | Percent classified root | Percent unclassified | Classified reads | Unclassified reads | Total reads |
|---|---|---|---|---|---|---|
| VT111 | fungus | 4.5300 | 95.470 | 128,149 | 2,697,674 | 2,825,823 |
| VT12 | fungus | 2.8800 | 97.120 | 93,406 | 3,146,248 | 3,239,654 |
| VT16 | fungus | 3.2100 | 96.790 | 91,412 | 2,755,863 | 2,847,275 |
| VT1 | fungus | 1.4400 | 98.560 | 306,389 | 21,024,557 | 21,330,946 |
| VT20 | fungus | 2.6900 | 97.310 | 72,333 | 2,620,247 | 2,692,580 |
| VT24 | fungus | 3.0400 | 96.960 | 50,276 | 1,605,743 | 1,656,019 |
| VT30 | fungus | 4.3100 | 95.690 | 251,876 | 5,596,702 | 5,848,578 |
| VT34 | fungus | 3.3900 | 96.610 | 77,399 | 2,206,365 | 2,283,764 |
| VT40 | fungus | 2.7600 | 97.240 | 64,207 | 2,260,051 | 2,324,258 |
| VT44 | fungus | 3.4000 | 96.600 | 80,686 | 2,292,838 | 2,373,524 |
| VT4 | fungus | 3.0000 | 97.000 | 68,643 | 2,219,108 | 2,287,751 |
| VT50 | fungus | 3.0600 | 96.940 | 116,548 | 3,692,148 | 3,808,696 |
| VT52 | fungus | 3.3300 | 96.670 | 84,222 | 2,442,164 | 2,526,386 |
| VT56 | fungus | 2.9900 | 97.010 | 54,898 | 1,780,023 | 1,834,921 |
| VT70 | fungus | 4.8400 | 95.160 | 251,306 | 4,941,985 | 5,193,291 |
| VT8 | fungus | 3.0100 | 96.990 | 69,093 | 2,225,940 | 2,295,033 |

**Table S15. Microbiome diversity and dispersion**

Per-sample α-diversity (Shannon), within/between-population components, and distances to group centroids used in PERMANOVA/PERMDISP analyses.

| snake | Between sample mean | Within sample mean | shannon (alpha diversity) | dist to centroid |
|---|---|---|---|---|
| 09.06SFD | 0.6341 | 0.5675 | 2.8858 | 0.3681 |
| 1 | 0.7248 | 0.2432 | 2.0477 | 0.0778 |
| 10 | 0.7387 | 0.3285 | 3.0047 | 0.2514 |
| 10.16 | 0.6357 | 0.5613 | 2.7684 | 0.3564 |
| 10.17 | 0.6119 | 0.5312 | 2.3340 | 0.3211 |
| 10.21 | 0.7814 | 0.7039 | 1.4953 | 0.5852 |
| 10.34 | 0.6929 | 0.6464 | 2.1311 | 0.4962 |
| 10.4 | 0.6499 | 0.2635 | 2.4034 | 0.0733 |
| 11 | 0.7399 | 0.5917 | 2.5165 | 0.5713 |
| 11.04 | 0.6362 | 0.5652 | 2.6502 | 0.3648 |
| 11.07 | 0.6398 | 0.5534 | 2.6985 | 0.3491 |
| 11.1 | 0.7798 | 0.7028 | 1.4762 | 0.5778 |
| 11.24 | 0.6250 | 0.5566 | 2.9906 | 0.3521 |
| 11.28 | 0.6439 | 0.5618 | 2.8136 | 0.3629 |
| 11.29 | 0.6690 | 0.5872 | 2.4785 | 0.4010 |
| 11.3 | 0.6511 | 0.5664 | 2.2943 | 0.3645 |
| 11.31 | 0.6770 | 0.2978 | 2.0531 | 0.1634 |
| 11.32 | 0.5979 | 0.5016 | 2.7519 | 0.2568 |
| 11.35 | 0.5974 | 0.5020 | 2.7488 | 0.2554 |
| 11.36 | 0.6642 | 0.6226 | 2.3472 | 0.4576 |
| 11.38 | 0.5681 | 0.4940 | 2.8935 | 0.2386 |
| 11.4 | 0.6471 | 0.5788 | 2.6331 | 0.3788 |
| 11.41 | 0.6660 | 0.5978 | 2.2915 | 0.4133 |
| 11.43 | 0.6135 | 0.5459 | 2.6660 | 0.3340 |
| 11.44 | 0.6401 | 0.3159 | 2.0735 | 0.2000 |
| 11.45 | 0.5987 | 0.5184 | 3.1068 | 0.2898 |
| 11.46 | 0.6598 | 0.5917 | 2.4280 | 0.3948 |
| 11.52 | 0.6495 | 0.3636 | 2.0707 | 0.2629 |
| 11.53 | 0.6434 | 0.2605 | 2.0676 | 0.0897 |
| 11.54 | 0.6972 | 0.3575 | 1.8980 | 0.2610 |
| 11.57 | 0.6360 | 0.2591 | 2.4106 | 0.0600 |
| 11.58 | 0.7867 | 0.5601 | 1.8839 | 0.5062 |
| 11.64 | 0.6975 | 0.6009 | 1.8904 | 0.4415 |
| 11.68 | 0.6503 | 0.7368 | 2.0590 | 0.6947 |
| 3 | 0.7075 | 0.3141 | 2.3701 | 0.2263 |
| 12.11 | 0.7098 | 0.6915 | 2.0182 | 0.5929 |
| 12.19 | 0.6559 | 0.5718 | 2.2861 | 0.4253 |
| 12.21 | 0.6745 | 0.6048 | 2.2796 | 0.4239 |
| 12.23 | 0.6447 | 0.5635 | 3.0087 | 0.3677 |
| 12.25 | 0.6054 | 0.5190 | 2.7677 | 0.2894 |
| 12.26 | 0.6685 | 0.5693 | 2.2259 | 0.4071 |
| 12.32 | 0.6657 | 0.6142 | 2.2774 | 0.4514 |
| 12.34 | 0.6731 | 0.5870 | 1.9343 | 0.4222 |
| 12.39 | 0.6717 | 0.5738 | 2.1944 | 0.4137 |
| 12.4 | 0.6861 | 0.6481 | 2.1538 | 0.5360 |
| 12.43 | 0.6360 | 0.5614 | 2.6596 | 0.4002 |
| 12.48 | 0.6096 | 0.5452 | 2.6269 | 0.3329 |
| 12.49 | 0.6877 | 0.6445 | 2.0139 | 0.5394 |
| 12.5 | 0.7275 | 0.6621 | 2.0073 | 0.5620 |
| 12.51 | 0.7698 | 0.7804 | 3.4741 | 0.6793 |
| 12.52 | 0.6579 | 0.5811 | 2.5356 | 0.4366 |
| 12.61 | 0.6531 | 0.2625 | 2.3006 | 0.0710 |
| 12.64 | 0.6997 | 0.6322 | 2.1046 | 0.4862 |
| 12.71 | 0.6447 | 0.2612 | 2.1364 | 0.0810 |
| 12.73 | 0.6651 | 0.6243 | 1.7207 | 0.5018 |
| 12.74 | 0.8407 | 0.7611 | 0.5918 | 0.7150 |
| 12.75 | 0.7170 | 0.6390 | 1.8157 | 0.5503 |
| 162019 | 0.8889 | 0.4795 | 0.3313 | 0.2947 |
| 186035 | 0.6271 | 0.6993 | 2.5723 | 0.6182 |
| 186036 | 0.7441 | 0.4381 | 1.7763 | 0.1305 |
| 186037 | 0.7162 | 0.5222 | 1.4873 | 0.3137 |
| 19192 | 0.8882 | 0.4790 | 0.3293 | 0.2939 |
| 2 | 0.7462 | 0.2410 | 1.8895 | 0.0741 |
| 5 | 0.7527 | 0.2448 | 1.6227 | 0.0946 |
| 6 | 0.7531 | 0.2520 | 1.6115 | 0.1068 |
| 7 | 0.7186 | 0.3704 | 2.4020 | 0.3042 |
| 77 | 0.6468 | 0.6694 | 2.0286 | 0.5014 |

**Table S15. Microbiome diversity and dispersion**

Per-sample α-diversity (Shannon), within/between-population components, and distances to group centroids used in PERMANOVA/PERMDISP analyses.

| snake | Between sample mean | Within sample mean | shannon (alpha diversity) | dist to centroid |
|---|---|---|---|---|
| 78 | 0.6313 | 0.5533 | 2.4190 | 0.3168 |
| 8 | 0.7590 | 0.2423 | 1.7866 | 0.0865 |
| 80 | 0.6095 | 0.4995 | 2.3013 | 0.2318 |
| 81 | 0.8237 | 0.7437 | 2.2783 | 0.6920 |
| 82303 | 0.8369 | 0.9087 | 0.4384 | 0.8261 |
| 84 | 0.6010 | 0.5397 | 3.2521 | 0.3032 |
| 9 | 0.6919 | 0.3278 | 2.4334 | 0.2375 |
| 90 | 0.5619 | 0.4747 | 2.9273 | 0.1772 |
| 91 | 0.5813 | 0.4673 | 2.8829 | 0.1627 |
| 92 | 0.6124 | 0.5662 | 2.6012 | 0.3553 |
| 93 | 0.6502 | 0.6910 | 2.2678 | 0.5287 |
| 99 | 0.5770 | 0.4684 | 2.6597 | 0.1782 |
| VT1 | 0.6967 | 0.6649 | 3.2609 | 0.5248 |
| VT10 | 0.6277 | 0.5783 | 2.6640 | 0.3966 |
| VT100 | 0.7614 | 0.7973 | 1.4972 | 0.6936 |
| VT101 | 0.6627 | 0.6014 | 2.4080 | 0.4284 |
| VT102 | 0.6922 | 0.6434 | 2.3199 | 0.4782 |
| VT103 | 0.6278 | 0.4863 | 2.8087 | 0.2840 |
| VT104 | 0.6260 | 0.4884 | 2.8632 | 0.2890 |
| VT105 | 0.8472 | 0.8224 | 4.3604 | 0.7143 |
| VT107 | 0.6261 | 0.4862 | 2.8356 | 0.2823 |
| VT109 | 0.6258 | 0.4891 | 2.8191 | 0.2918 |
| VT110 | 0.6242 | 0.4941 | 2.8458 | 0.2970 |
| VT111 | 0.7080 | 0.6475 | 1.8165 | 0.5037 |
| VT12 | 0.6448 | 0.6433 | 2.8111 | 0.5000 |
| VT16 | 0.6186 | 0.5335 | 2.4501 | 0.3253 |
| VT20 | 0.7217 | 0.7573 | 1.7322 | 0.6434 |
| VT24 | 0.5572 | 0.4896 | 2.7898 | 0.2909 |
| VT30 | 0.6197 | 0.5700 | 2.8485 | 0.3809 |
| VT34 | 0.5564 | 0.4893 | 2.8772 | 0.2933 |
| VT4 | 0.5691 | 0.4996 | 2.7665 | 0.2870 |
| VT40 | 0.6476 | 0.5907 | 2.6491 | 0.4127 |
| VT44 | 0.5612 | 0.4856 | 3.0736 | 0.2815 |
| VT50 | 0.6784 | 0.6202 | 2.5623 | 0.4486 |
| VT52 | 0.5718 | 0.5288 | 2.4343 | 0.3508 |
| VT56 | 0.5526 | 0.4819 | 2.8164 | 0.2820 |
| VT70 | 0.6664 | 0.6706 | 2.0219 | 0.5320 |
| VT8 | 0.6129 | 0.5540 | 2.6958 | 0.3640 |

**Table S16. Ophidiomyces ophidiicola read counts**

Per-sample pathogen detection metrics: total reads, reads masked as low complexity, percent low complexity, and final filtered O. ophidiicola reads.

| snake | Total Reads | Reads in LowComplexity | Fraction in LowComplexity | Final Oo Reads | Oo burden |
|---|---|---|---|---|---|
| 12.51 | 10,200 | 3,543 | 0.3474 | 6,657 | 0.0126 |
| 186037 | 34,719 | 20,963 | 0.6038 | 13,756 | 0.0053 |
| 186036 | 41,978 | 25,592 | 0.6097 | 16,386 | 0.0033 |
| 186035 | 56,246 | 36,358 | 0.6464 | 19,888 | 0.0074 |
| 11 | 92,578 | 59,072 | 0.6381 | 33,506 | -0.0004 |
| 80 | 137,184 | 92,196 | 0.6721 | 44,988 | 0.0001 |
| 12.5 | 122,775 | 72,620 | 0.5915 | 50,155 | 0.0068 |
| VT102 | 212,263 | 161,740 | 0.7620 | 50,523 | -0.0062 |
| 11.1 | 131,566 | 72,573 | 0.5516 | 58,993 | 0.0102 |
| VT24 | 353,668 | 283,771 | 0.8024 | 69,897 | -0.0015 |
| 92 | 245,698 | 173,231 | 0.7051 | 72,467 | -0.0043 |
| 12.49 | 224,805 | 151,057 | 0.6719 | 73,748 | -0.0029 |
| 12.11 | 236,318 | 162,466 | 0.6875 | 73,852 | -0.0039 |
| 11.73 | 205,456 | 130,872 | 0.6370 | 74,584 | -0.0017 |
| 78 | 275,342 | 198,234 | 0.7200 | 77,108 | -0.0043 |
| 10.4 | 411,826 | 329,831 | 0.8009 | 81,995 | -0.0022 |
| VT56 | 386,747 | 303,111 | 0.7837 | 83,636 | -0.0011 |
| 11.64 | 287,995 | 204,266 | 0.7093 | 83,729 | -0.0044 |
| 11.38 | 393,243 | 308,350 | 0.7841 | 84,893 | -0.0040 |
| 93 | 399,032 | 313,896 | 0.7866 | 85,136 | -0.0028 |
| 77 | 358,590 | 271,784 | 0.7579 | 86,806 | -0.0050 |
| 12.74 | 125,964 | 36,442 | 0.2893 | 89,522 | 0.0069 |
| 10 | 303,584 | 213,477 | 0.7032 | 90,107 | -0.0053 |
| 10.17 | 266,793 | 175,821 | 0.6590 | 90,972 | -0.0029 |
| 84 | 373,390 | 280,581 | 0.7514 | 92,809 | -0.0045 |
| 91 | 362,971 | 268,967 | 0.7410 | 94,004 | -0.0049 |
| 12.43 | 407,173 | 311,409 | 0.7648 | 95,764 | -0.0046 |
| 12.34 | 276,301 | 179,156 | 0.6484 | 97,145 | -0.0009 |
| 11.57 | 437,566 | 339,487 | 0.7759 | 98,079 | -0.0015 |
| 90 | 419,786 | 321,644 | 0.7662 | 98,142 | -0.0021 |
| 11.45 | 463,147 | 364,804 | 0.7877 | 98,343 | -0.0022 |
| 12.61 | 401,460 | 301,801 | 0.7518 | 99,659 | -0.0039 |
| 11.54 | 407,614 | 307,172 | 0.7536 | 100,442 | -0.0033 |
| VT109 | 443,536 | 338,936 | 0.7642 | 104,600 | -0.0023 |
| 11.4 | 520,878 | 414,916 | 0.7966 | 105,962 | -0.0030 |
| 6 | 417,936 | 311,055 | 0.7443 | 106,881 | -0.0029 |
| VT8 | 497,988 | 387,119 | 0.7774 | 110,869 | 0.0015 |
| 12.75 | 260,412 | 149,255 | 0.5731 | 111,157 | 0.0029 |
| VT34 | 497,414 | 385,647 | 0.7753 | 111,767 | -0.0004 |
| 10.34 | 342,462 | 227,120 | 0.6632 | 115,342 | 0.0022 |
| 12.25 | 431,825 | 316,192 | 0.7322 | 115,633 | -0.0023 |
| 11.53 | 436,775 | 320,912 | 0.7347 | 115,863 | -0.0029 |
| VT4 | 470,583 | 353,930 | 0.7521 | 116,653 | 0.0002 |
| 11.58 | 257,542 | 140,724 | 0.5464 | 116,818 | -0.0013 |
| 3 | 418,762 | 301,128 | 0.7191 | 117,634 | -0.0033 |
| VT110 | 507,597 | 388,314 | 0.7650 | 119,283 | -0.0017 |
| VT40 | 309,712 | 189,944 | 0.6133 | 119,768 | 0.0043 |
| 11.28 | 501,094 | 380,628 | 0.7596 | 120,466 | -0.0025 |
| 10.21 | 322,005 | 199,926 | 0.6209 | 122,079 | 0.0003 |
| 11.24 | 433,322 | 309,537 | 0.7143 | 123,785 | -0.0024 |
| 99 | 469,848 | 345,173 | 0.7346 | 124,675 | -0.0026 |
| VT52 | 476,280 | 350,873 | 0.7367 | 125,407 | -0.0006 |
| VT44 | 511,443 | 385,141 | 0.7530 | 126,302 | 0.0004 |
| VT101 | 524,816 | 397,955 | 0.7583 | 126,861 | 0.0005 |
| VT20 | 280,142 | 152,517 | 0.5444 | 127,625 | 0.0077 |
| 11.46 | 529,407 | 401,193 | 0.7578 | 128,214 | -0.0009 |
| 12.26 | 485,879 | 357,369 | 0.7355 | 128,510 | -0.0009 |
| 11.41 | 517,873 | 388,503 | 0.7502 | 129,370 | -0.0007 |
| 11.04 | 450,846 | 320,583 | 0.7111 | 130,263 | -0.0022 |
| 11.07 | 472,345 | 341,995 | 0.7240 | 130,350 | -0.0006 |
| 12.19 | 448,094 | 316,445 | 0.7062 | 131,649 | -0.0020 |
| 2 | 439,259 | 307,235 | 0.6994 | 132,024 | -0.0016 |
| 12.64 | 445,592 | 313,324 | 0.7032 | 132,268 | -0.0031 |
| 12.23 | 465,741 | 333,163 | 0.7153 | 132,578 | -0.0023 |
| 09.06SFD | 510,663 | 377,875 | 0.7400 | 132,788 | -0.0009 |
| 12.71 | 504,579 | 371,629 | 0.7365 | 132,950 | -0.0018 |
| VT103 | 554,888 | 420,047 | 0.7570 | 134,841 | 0.0010 |
| 12.52 | 459,712 | 323,840 | 0.7044 | 135,872 | -0.0020 |

**Table S16. Ophidiomyces ophidiicola read counts**

Per-sample pathogen detection metrics: total reads, reads masked as low complexity, percent low complexity, and final filtered O. ophidiicola reads.

| snake | Total Reads | Reads in LowComplexity | Fraction in LowComplexity | Final Oo Reads | Oo burden |
|---|---|---|---|---|---|
| 11.43 | 471,870 | 335,408 | 0.7108 | 136,462 | -0.0015 |
| VT104 | 530,938 | 394,301 | 0.7426 | 136,637 | 0.0003 |
| 11.35 | 476,115 | 336,827 | 0.7074 | 139,288 | -0.0025 |
| 11.44 | 538,698 | 398,561 | 0.7399 | 140,137 | 0.0010 |
| 10.16 | 478,889 | 337,194 | 0.7041 | 141,695 | -0.0026 |
| 12.39 | 532,762 | 388,627 | 0.7295 | 144,135 | 0.0002 |
| 12.21 | 511,848 | 365,629 | 0.7143 | 146,219 | -0.0010 |
| 12.48 | 459,710 | 312,654 | 0.6801 | 147,056 | 0.0010 |
| 11.32 | 572,348 | 425,177 | 0.7429 | 147,171 | -0.0011 |
| VT16 | 590,404 | 442,876 | 0.7501 | 147,528 | 0.0025 |
| 5 | 531,034 | 383,449 | 0.7221 | 147,585 | -0.0004 |
| 1 | 576,331 | 427,131 | 0.7411 | 149,200 | -0.0009 |
| 11.3 | 523,553 | 373,530 | 0.7135 | 150,023 | -0.0002 |
| 11.52 | 589,679 | 439,250 | 0.7449 | 150,429 | 0.0001 |
| VT107 | 602,288 | 450,541 | 0.7480 | 151,747 | 0.0005 |
| 12.4 | 503,983 | 350,309 | 0.6951 | 153,674 | -0.0014 |
| 8 | 494,376 | 339,261 | 0.6862 | 155,115 | 0.0002 |
| VT100 | 373,060 | 216,938 | 0.5815 | 156,122 | -0.0006 |
| 9 | 613,269 | 455,416 | 0.7426 | 157,853 | 0.0007 |
| 11.31 | 578,299 | 416,893 | 0.7209 | 161,406 | 0.0017 |
| 11.29 | 509,518 | 346,607 | 0.6803 | 162,911 | -0.0009 |
| VT111 | 527,635 | 363,222 | 0.6884 | 164,413 | 0.0013 |
| 11.36 | 557,594 | 392,757 | 0.7044 | 164,837 | -0.0006 |
| VT12 | 493,669 | 325,224 | 0.6588 | 168,445 | -0.0008 |
| 7 | 566,499 | 394,869 | 0.6970 | 171,630 | -0.0002 |
| 12.32 | 631,246 | 456,111 | 0.7226 | 175,135 | 0.0007 |
| 11.74 | 485,207 | 286,569 | 0.5906 | 198,638 | 0.0019 |
| 6439 | 244,743 | 34,021 | 0.1390 | 210,722 | NA |
| VT50 | 838,534 | 599,138 | 0.7145 | 239,396 | 0.0009 |
| R-4404 | 333,135 | 51,255 | 0.1539 | 281,880 | NA |
| 162019 | 331,998 | 48,778 | 0.1469 | 283,220 | -0.0011 |
| VT70 | 748,882 | 451,029 | 0.6023 | 297,853 | 0.0026 |
| 82303 | 383,410 | 65,237 | 0.1701 | 318,173 | 0.0062 |
| VT30 | 1,304,186 | 947,438 | 0.7265 | 356,748 | 0.0035 |
| 81 | 454,017 | 75,275 | 0.1658 | 378,742 | 0.0027 |
| 27544 | 443,277 | 62,484 | 0.1410 | 380,793 | NA |
| VT105 | 479,147 | 95,576 | 0.1995 | 383,571 | -0.0017 |
| 11.68 | 1,590,706 | 1,171,927 | 0.7367 | 418,779 | 0.0155 |
| 12.73 | 1,509,506 | 1,002,467 | 0.6641 | 507,039 | 0.0045 |
| 6462 | 614,156 | 76,037 | 0.1238 | 538,119 | NA |
| 19192 | 648,545 | 83,813 | 0.1292 | 564,732 | 0.0035 |
| VT10 | 2,283,575 | 1,454,073 | 0.6368 | 829,502 | 0.0108 |
| VT1 | 2,174,662 | 1,254,135 | 0.5767 | 920,527 | 0.0092 |
